# A promiscuous mechanism to phase separate eukaryotic carbon fixation in the green lineage

**DOI:** 10.1101/2024.04.09.588658

**Authors:** James Barrett, Mihris I.S. Naduthodi, Yuwei Mao, Clément Dégut, Sabina Musiał, Aidan Salter, Mark C. Leake, Michael J. Plevin, Alistair J. McCormick, James N. Blaza, Luke C.M. Mackinder

**Affiliations:** Department of Biology, University of York; York, YO10 5DD, UK; Centre for Novel Agricultural Products (CNAP), Department of Biology, University of York; York, YO10 5DD, UK; Institute of Molecular Plant Sciences, School of Biological Sciences, University of Edinburgh; Edinburgh, EH9 3BF, UK; Centre for Engineering Biology, University of Edinburgh; Edinburgh, EH9 3BF, UK.; School of Physics, Engineering and Technology, University of York; York, YO10 5DD, UK.; York Structural Biology Laboratory, The University of York; York, YO10 5DD, UK; Department of Chemistry, York Structural Biology Laboratory; York, YO10 5DD, UK

## Abstract

CO_2_ fixation is commonly limited by inefficiency of the CO_2_-fixing enzyme Rubisco. Eukaryotic algae concentrate and fix CO_2_ in phase-separated condensates called pyrenoids, which complete up to one-third of global CO_2_ fixation. Condensation of Rubisco in pyrenoids is dependent on interaction with disordered linker proteins that show little conservation between species. We developed a sequence-independent bioinformatic pipeline to identify linker proteins in green algae. We report the linker from *Chlorella* and demonstrate that it binds a conserved site on the Rubisco large subunit. We show the *Chlorella* linker phase separates *Chlamydomonas* Rubisco and that despite their separation by ∼800 million years of evolution, the *Chlorella* linker can support the formation of a functional pyrenoid in *Chlamydomonas*. This cross-species reactivity extends to plants, with the *Chlorella* linker able to drive condensation of some native plant Rubiscos *in vitro* and *in planta*. Our results represent an exciting frontier for pyrenoid engineering in plants, which is modelled to increase crop yields.

---

As the primary gateway between atmospheric carbon dioxide (CO_2_) and organic carbon, ribulose-1,5-bisphosphate carboxylase/oxygenase (Rubisco) fixes approximately 400 gigatons of CO_2_ annually^1^. Despite this huge global productivity, Rubisco as an enzyme is catalytically slow^2^. Its Archean origin^3^ and exceptionally slow evolutionary trajectory^4^, has meant Rubisco has failed to significantly overcome the supposed trade-off between specificity for its substrate (CO_2_ or O_2_) and catalytic rate^4,5^. These shortfalls mean Rubisco is often limiting for photosynthesis in most plants (i.e. C_3_ plants)^6–11^. Accordingly, C_3_ plants compensate by producing and maintaining large amounts of Rubisco^12,13^. By mass, Rubiscos in aquatic phototrophs (algae and cyanobacteria) are ∼20 times more efficient^14^. These Rubiscos benefit from operating in biophysical CO_2_-concentrating mechanisms (CCMs) that increase the CO_2_:O_2_ ratio at their active site. A large proportion of aquatic CO_2_-fixation occurs in the pyrenoid^15^, a sub-compartment of the chloroplast found in most eukaryotic algae and some basal land plants that is the centerpiece of their biophysical CCMs^16^. Pyrenoid formation is underpinned by biomolecular condensation of Rubisco by disordered linker proteins^15,17^ (Fig. 1a). Engineering pyrenoid-based CCMs in crop plants that operate C_3_ photosynthesis is a promising avenue to increase their primary productivity and reduce nitrogen and water usage^18,19^. While significant progress has been made in the characterization and transfer of pyrenoid components from the model alga *Chlamydomonas reinhardtii* to the model C_3_ plant *Arabidopsis thaliana*^15,20–24^, our knowledge of pyrenoids from other species remains limited. By characterizing pyrenoids from other species, we hope to expand the toolbox available for plant pyrenoid engineering and gain insight into the commonalties and differences between pyrenoid assembly.

**Fig. 1.**
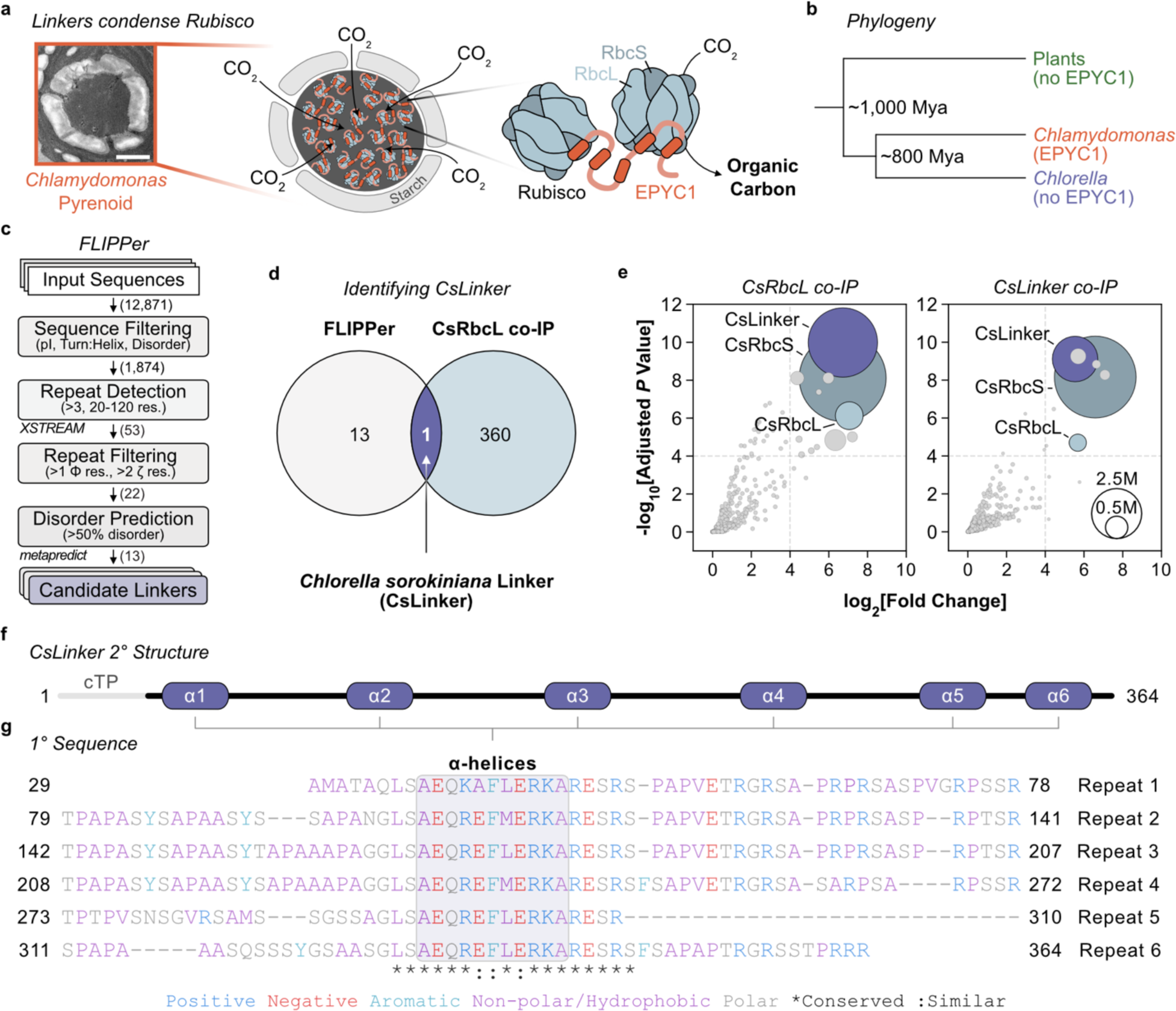
Identification of the *Chlorella sorokiniana* linker protein (CsLinker). **a**, Transmission electron micrograph (TEM) of the *Chlamydomonas reinhardtii* pyrenoid, with adjacent schematic of Rubisco condensation in the pyrenoid by interaction of EPYC1 helices with the Rubisco small subunits (RbcSs). Condensed Rubisco fixes CO2 to organic carbon. Scale bar = 1 μm. **b**, Phylogeny of *Chlamydomonas*, *Chlorella* and plants. Estimated divergence points from a time-calibrated phylogeny28. **c**, Schematic representation of the Fast Linker Identification Pipeline for Pyrenoids (FLIPPer) used to identify candidate linkers that share features with EPYC1. Where relevant, the program used is indicated. The number of sequences remaining after each filtering step of the *Chlorella sorokiniana* UTEX1230 genome is indicated. Abbreviations: pI = isoelectric point, res. = residue, Φ = hydrophobic, ζ = electrostatic. **d**, Venn diagram demonstrating identification of CsLinker from FLIPPer and CsRbcL co-immunoprecipitation (co-IP). **e**, Reciprocal co-IP experiments performed using antibodies raised to the Rubisco large subunit (left) and CsLinker (right). Dashed lines indicate arbitrary significance thresholds (−log10[Adjusted *P* Value] > 4, log2[Fold Change] > 4), above which points are sized according to their summed abundance (M = millions) from three replicates as per the inset key. **f**, Predicted secondary structure of CsLinker from AlphaFold modelling (Extended Data Fig. 1). The predicted chloroplast transit peptide (cTP) and α-helices (α1-6) are indicated. **g**, Primary sequence alignment of the six repeat regions of CsLinker, colored by residue property.

Here, we identify and characterize CsLinker, a pyrenoid linker protein from the green alga *Chlorella* which is an ancient relative of *Chlamydomonas*. Using biochemical and structural approaches, we demonstrate that CsLinker is functionally analogous to linker proteins from other organisms, despite low sequence identity. Crucially, and in contrast to the *Chlamydomonas* linker EPYC1, we show that CsLinker binds to the Rubisco large subunit (RbcL). The high conservation of the binding interface on the RbcL supports functional cross-reactivity of *Chlamydomonas* Rubisco and CsLinker as well as CsLinker-mediated condensation of native plant Rubiscos both *in vitro* and *in planta*. These findings represent a significant advance towards the engineering of synthetic pyrenoids and overcome a major hurdle for the future engineering of pyrenoid-based CCMs in plants.

## Results

### A Fast Linker Identification Pipeline for Pyrenoids (FLIPPer) identifies the pyrenoid linker in ***Chlorella sorokiniana***

The *Chlamydomonas* pyrenoid linker protein EPYC1 is not conserved outside closely related species^15^ (Fig. 1b). To identify functional analogues in other pyrenoid-containing species of the green lineage, we developed a sequence homology-independent Fast Linker Identification Pipeline for Pyrenoids (FLIPPer). FLIPPer searches for proteins with features key to the function of EPYC1, namely: i) largely disordered, with ii) repeating structured elements, that are iii) spaced at a feasible length scale for cross-linking Rubiscos (20-120 residues; ∼2-8 nm^25,26^) (Fig. 1c). Amongst our analyses, we identified 13 proteins with these features in the genome of the unicellular Trebouxiophyte *Chlorella sorokiniana* UTEX1230^27^ (Chlorella hereafter) (Extended Data Fig. 1), which diverged from *Chlamydomonas* ∼800 million years ago (Mya)^28^ (Fig. 1b). To validate the identity of the Chlorella linker, we completed co-immunoprecipitation (co-IP) experiments on Chlorella cells grown in low CO_2_, using an antibody specific to the Rubisco large subunit (RbcL) (Fig. 1e, Extended Data Fig. 2a,b and Supplementary Table 1). Of the 360 proteins enriched relative to control co-IPs (*Chlamydomonas* BST2 antibody), only 1 was shared amongst the FLIPPer outputs (Fig. 1d). This protein, CSI2_123000012064, was the most significantly enriched in the RbcL co-IP experiment alongside both subunits of the Chlorella Rubisco holoenzyme (CsRbcL and CsRbcS) (Fig. 1e, left). We called this protein CsLinker. We subsequently completed reciprocal co-IPs using a CsLinker antibody (Extended Data Fig. 2c,d), which enriched both CsRbcL and CsRbcS, indicating *ex vivo* complex formation between CsLinker and Chlorella Rubisco (Fig. 1e, right). We confirmed the CsLinker gene sequence, gene model and predicted chloroplast transit peptide (cTP) by PCR and sequencing and through mapping of peptides identified in mass spectrometry experiments (Extended Data Fig. 3). We further confirmed the low CO_2_ inducibility of CsLinker through analysis of available RNA-seq data, and by western blotting (Extended Data Fig. 4 and Supplementary Table 2). Whilst AlphaFold 2 modelling demonstrated CsLinker and EPYC1 share clear structural analogy, their primary sequences share little similarity (Fig. 1g, 25% identity; Extended Data Fig. 5), reflecting their independent origins.

### CsLinker is a *bona fide* pyrenoid linker protein

Having demonstrated *ex vivo* interaction between CsLinker and CsRubisco (Fig. 1e), we sought to confirm both were abundant components of the Chlorella pyrenoid *in vivo*. Using the RbcL antibody, we completed immunoelectron microscopy and observed CsRubisco almost exclusively localized in the pyrenoid (96.2 ± 3.5 % S.D., n=6) (Fig. 2a, Extended Data Fig. 6 and Supplementary Table 3). We observed no immuno-labelling using the CsLinker antibody, likely due to poor antibody antigenicity (Extended Data Fig. 6c). To determine the abundance of CsLinker and CsRubisco *in vivo*, we completed absolute quantification mass spectrometry using standard curves of recombinant CsLinker and CsRubisco purified from Chlorella. Accounting for the chloroplast volume^29^, we measured the concentration of CsLinker and CsRubisco to be 4.58 ± 0.53 μM S.D. and 2.16 ± 0.04 μM S.D. respectively (Fig. 2b, Extended Data Fig. 7 and Supplementary Tables 4-7), demonstrating CsLinker is highly abundant in the chloroplast. We validated the Rubisco quantification by western blotting (2.42 μM; asterisk in Fig. 2b, Extended Data Fig. 2b and Supplementary Table 8). Alongside the localization of CsRubisco in the pyrenoid (Fig. 2a), the two-fold abundance of CsLinker over CsRubisco *in vivo* (Fig. 2b) and their *ex vivo* interaction (Fig. 1e) indicate CsLinker likely interacts with CsRubisco as an abundant component of the Chlorella pyrenoid *in vivo*.

**Fig. 2.**
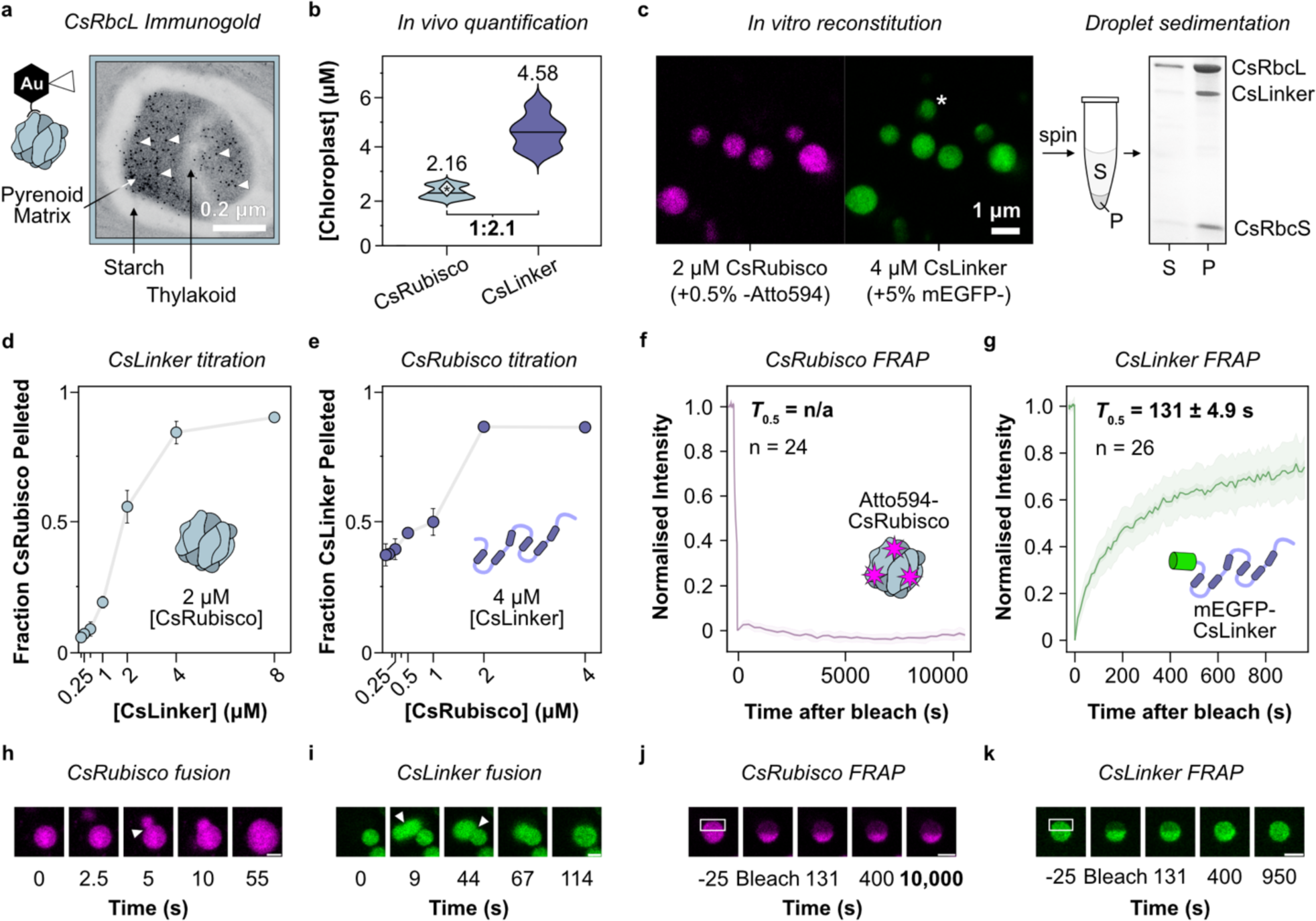
CsLinker phase separates Rubisco at physiological conditions. **a**, Representative immunogold TEM of the Chlorella pyrenoid after primary incubation with the RbcL antibody. A subset of gold nanoparticles are indicated by white arrowheads. **b**, Violin plot from absolute quantification of CsRubisco holoenzyme (derived from CsRbcL) and CsLinker *in vivo* (n = 3). The ratio between CsRubisco and CsLinker is indicated below. (See Extended Data Fig. 7 for details). The asterisked point indicates the independent quantification from western blotting (Extended Data Fig. 2b). **c**, Confocal fluorescence microscopy image of droplets in the *in vitro* reconstitution of the Chlorella pyrenoid formed at the measured chloroplast concentrations of CsRubisco and CsLinker (2 μM and 4 μM, respectively) (left panel). Asterisk indicates a droplet that settled between imaging of the two channels. Atto594-CsRubisco and mEGFP-CsLinker were incorporated at 0.5% and 5% molar concentrations respectively. Droplets can be sedimented by centrifugation and the composition of the pellet (P), relative to the supernatant (S) analyzed by SDS-PAGE (right panel). This experiment format was used to generate datapoints in d and e. **d**, Titration droplet sedimentation assays with CsRubisco fixed at the measured chloroplast concentration. **e**, Titration droplet sedimentation assays with fixed CsLinker. Where visible, error bars represent the S.D. from two replicates. **f**, Average full-scale normalized half-FRAP recovery curve of Atto594-CsRubisco in droplets formed as in c. The mean, S.E.M and S.D. are represented by the line, the smaller shader region, and the larger shaded region respectively. **g**, Average full-scale normalized half-FRAP recovery curve of mEGFP-CsLinker in droplets with the same composition. **h**, Time series of 0.05% (molar ratio) Atto594-CsRubisco labelled droplets formed as in c, undergoing fusion (arrow) and relaxation. **i**, 5% mEGFP-CsLinker labelled droplets undergoing consecutive fusions. **j**, Time series of a representative CsRubisco FRAP experiment, as quantified in f. The white box indicates the region bleached. **k**, Representative CsLinker FRAP experiment, as quantified in g. Scale bars for panels h-k = 1 μm.

The *Chlamydomonas* pyrenoid is a liquid-liquid phase-separated (LLPS) biomolecular condensate *in vivo*^26^, which is essential to Rubisco packaging and pyrenoid function^15^. *In vitro*, reconstituted pyrenoids formed by mixing purified linker EPYC1 and *Chlamydomonas* Rubisco demonstrate similar properties to *in vivo*^30^. Accordingly, we sought to understand whether mixing CsLinker and CsRubisco gives rise to similar emergent properties *in vitro*. When mixed at the chloroplast concentrations we measured (Fig. 2b), we observed demixing into micron-scale droplets that was dependent on, and incorporated both, CsLinker and CsRubisco (Fig. 2c, Extended Data Fig. 8a,b). To assess the relative occupancy of the components within droplets, we completed reciprocal titration in droplet sedimentation assays (Fig. 2c, right). By separately fixing both components at their chloroplast concentrations, we observed requirement of an approximately two-fold excess of CsLinker to fully demix both components (Fig. 2d,e, Extended Data Fig. 8c,d). This observation agreed with the approximately two-fold abundance of CsLinker we measured *in vivo* (Fig. 2b), and previous observations in *Chlamydomonas*^30^. Using the same ratio, we observed a critical global concentration (0.3-0.5 μM) and salt-dependency for droplet formation (Extended Data Fig. 9); both key indicators of LLPS^31^. To further understand the droplet properties, we completed fluorescence recovery after photobleaching (FRAP) experiments, which allowed us to monitor the mobility of labelled CsLinker and CsRubisco. Strikingly, we observed largely different mobilities for CsRubisco and CsLinker in droplets. Whilst mEGFP-CsLinker exhibited exchange with the dilute phase in whole FRAP experiments (*T*_0.5_ = 201 ± 14.3 s S.E.M.; Extended Data Fig. 10a and Supplementary Table 9) and internal mixing in half-FRAP experiments (*T*_0.5_ = 131 ± 4.89 s S.E.M.; Fig. 2g,k), Atto594-CsRubisco demonstrated no mixing or exchange over hour timescales (Fig. 2f,j). However, both CsLinker and CsRubisco appeared to undergo internal rearrangement over second timescales upon droplet fusion (Fig. 2h,i), suggesting CsRubisco is not immobilized in droplets, akin to observations of the *Phaeodactylum tricornutum* pyrenoid reconstitution^17^. To ensure our observations were not set-up specific, we confirmed the mobility of *Chlamydomonas* Rubisco (CrRubisco) and EPYC1 in the *Chlamydomonas* pyrenoid reconstitution using the same strategy. Consistent with previous *in vitro*^30^ and *in vivo*^26^ observations, both components were highly mobile (EPYC1-mEGFP half-FRAP *T*_0.5_ = 22 ± 6.3 s S.E.M., Atto594-CrRubisco half-FRAP *T*_0.5_ = 55 ± 5.1 s S.E.M.; Extended Data Fig. 10d,e). Taken together these results demonstrate the ability of CsLinker to phase separate Chlorella Rubisco *in vitro*, and alongside our *ex vivo* and *in vivo* observations, indicate this process likely underpins pyrenoid formation *in vivo*, analogous to EPYC1 in *Chlamydomonas*.

### CsLinker binds to the Rubisco large subunit (RbcL)

Previously characterized Rubisco-condensing linker proteins from algal pyrenoids^17,32^ and bacterial carboxysomes^33,34^ utilize structured regions to bind different regions on Rubisco. We hypothesized that the predicted α-helical regions in CsLinker may bind to Rubisco and that this interaction would involve a previously uncharacterized interface. To characterize the interaction, we produced fragments of CsLinker encompassing entire repeat sequences centered on predicted α-helix 3 (α3) and α-helices 3 and 4 (α3-α4) (Fig. 1f,g, Extended Data Fig. 11a). Using 2D NMR spectroscopy, we confirmed the only stable structure in the fragments to be α-helices of approximately 10 residues, in line with structural predictions (Fig. 1f, Extended Data Fig. 5a, 11). The similarity of the NMR spectra of α3 and α3-α4 indicated the individual repeat regions have similar overall properties (Extended Data Fig. 11d), consistent with their similar residue composition and the lack of stable tertiary structure. We next tested the ability of the fragments to interact with CsRubisco. In line with the dependency of LLPS on multivalent interactions for cross-linking, the single helix α3 fragment was unable to phase-separate CsRubisco, but did demonstrate concentration-dependent mobility shift of CsRubisco in native PAGE experiments (Extended Data Fig. 12a-c). Surprisingly, the α3-α4 double repeat fragment was also unable to induce LLPS but did demonstrate formation of a stable higher-order complex with CsRubisco that had a half-occupancy (*K*_0.5_) of ∼1.2 μM (CI_95_: 0.93-1.42 μM) (Fig. 3a, Extended Data Fig. 12d-h and Supplementary Table 10). To measure the affinity of the fragments for CsRubisco, we used surface plasmon resonance (SPR) experiments in which CsRubisco was immobilized as bait, and the α3 and α3-α4 fragments were used as prey (Fig. 3b, left). We measured a similar affinity (*K*_D_) of the α3-α4 fragment for CsRubisco (1.21 μM [CI_95_: 1.10–1.33 μM]) as we observed for the *K*_0.5_ of the α3-α4-CsRubisco complex by native PAGE (Fig. 3a, Extended Data Fig. 13 and Supplementary Table 11). The *K*_D_ of the α3 fragment was ∼100-fold higher (103 μM [CI_95_: 80–133 μM]) than α3-α4, consistent with cooperative binding of the two α-helical regions in α3-α4 to the same Rubisco rather than other Rubiscos in solution. This explained the lack of LLPS, and indicated the higher-order complex observed by native PAGE likely consists of single Rubiscos bound by α3-α4.

**Fig. 3.**
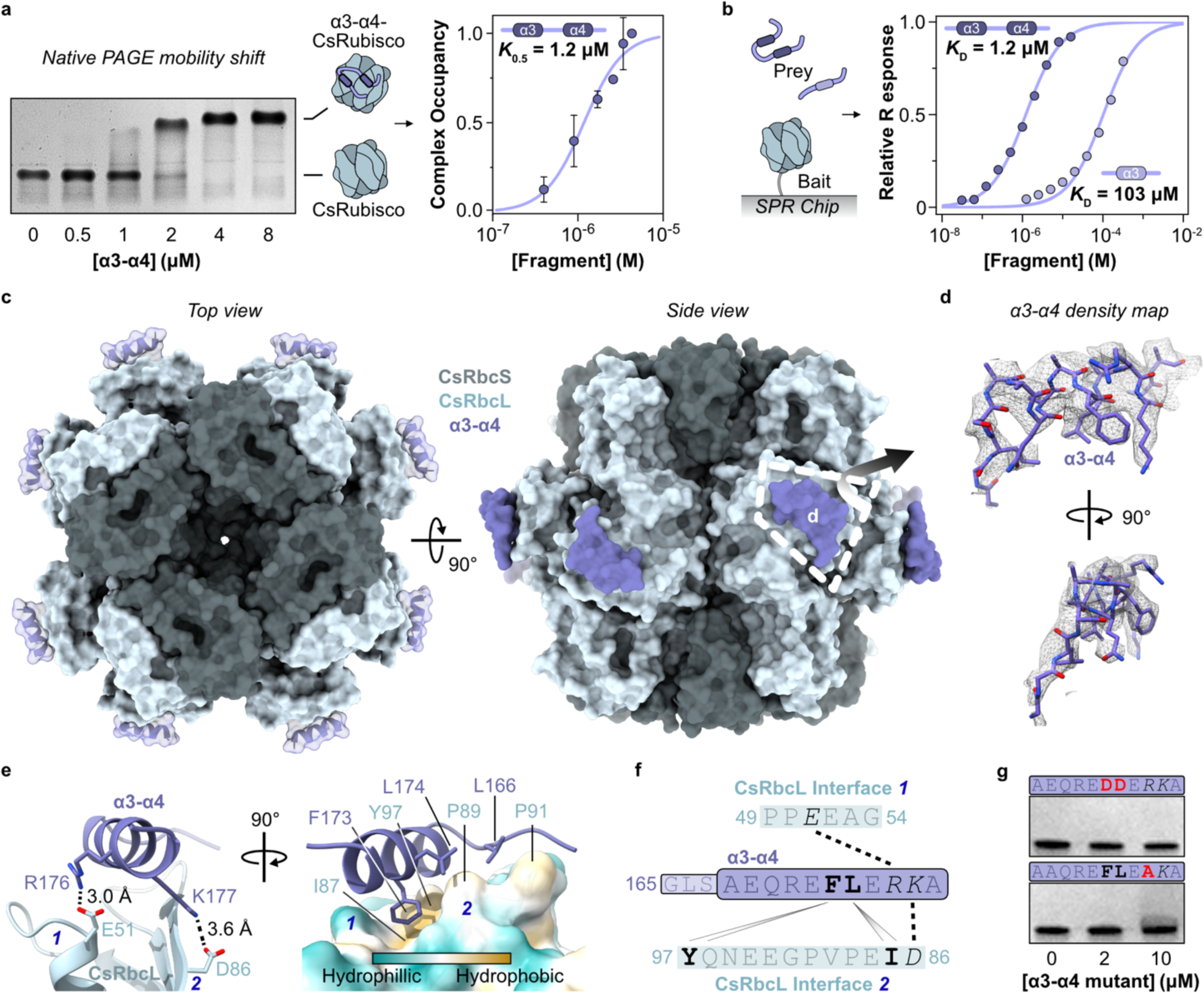
CsLinker binds to the Rubisco large subunit (RbcL). **a**, α3-α4 forms a stable complex with CsRubisco. Representative native PAGE gel shift assay (left) demonstrating formation of higher order α3-α4-CsRubisco complex. Quantification of replicate data (right) taken from Extended data Fig. 12h; n=2, error bars = S.D. **b**, Schematic of surface plasmon resonance (SPR) experiments (left) used to determine binding alinity of α3 and α3-α4 for CsRubisco (right). SPR response normalized to *B*max value obtained from fit of raw data; n=3, error bars = S.D. **c**, Top (left) and side (right) views of surface representations of cryo-EM determined structure of the α3-α4-CsRubisco complex. The modelled α-helix of α3 is superimposed on each CsRbcL based on the coordinates built into one axis of the C1 map, as shown in d. **d**, Density map of the α3-α4 region in the C1 complex map, carved with a radius of 2 from the built coordinates, at a contour level of 0.033. **e**, Molecular interactions at the interface. Shortest range electrostatic interactions highlighted by PDBePISA (left) and residues contributing to the hydrophobic interface (right). The surface of CsRbcL is colored according to hydrophobicity. Residues are numbered according to their position in the full length CsLinker. **f**, Map of the interactions between the α-helices of α3-α4 and the two interfaces on CsRbcL. Dashes indicate salt bridges, with wedges representing significant contributions to the hydrophobic interface, with italicised and bolded residues contributing the same interfaces, respectively. **g**, Native PAGE gel shift assays showing that mutation of α3-α4 disrupts binding to CsRubisco. The sequence of the α-helices in each fragment is provided above each image.

To determine where CsLinker binds CsRubisco, we elucidated the 3D structure of the α3-α4-CsRubisco complex using cryogenic electron microscopy (cryo-EM) (Supplementary Table 12). We prepared samples of α3-α4 and CsRubisco such that the solution concentration of CsLinker repeats was comparable to the value we measured in the chloroplast (32 μM in this experiment, 27.5 μM *in vivo*; Fig. 2b). This concentration also saturated the α3-α4-CsRubisco complex (Fig. 3a). The 2.4 Å map and model we obtained with D4 reconstruction (PDB: 8Q04) was highly similar to a previously determined cryo-EM structure of *Chlamydomonas* Rubisco^32^ (Cα RMSD of RbcL = 1.106 Å and RbcS = 0.937 Å, Extended Data Fig. 14k). During processing we observed low-resolution density additional to the core subunits at the equatorial region of CsRubisco in the RbcL (Extended Data Fig. 14d). We hypothesized this additional density corresponded to the helical regions of α3-α4 and that sub-stoichiometric binding meant many binding sites were unoccupied, giving rise to poorly resolved density where the linker binds. Using a symmetry expansion approach^35^, with a soft featureless mask around the additional density (Extended Data Fig. 15a), we found ∼23% of the 8 CsRbcL sites to be occupied by additional density in the symmetry expanded sub-particles (Extended Data Fig. 15c,f). By focusing on the sub-particles bound by α3-α4, we obtained a map of the additional density which possessed clearly helical nature (Fig. 3d, Extended Data Fig. 15e; PDB: 8Q05). We built the helical region of α3 into the density and numbered residues according to their position in the full-length protein (Fig. 1g, Fig. 3e, Extended Data Fig. 15h,i). Each CsLinker binding site is contained within the N-terminal region of a single CsRbcL subunit, utilizing two salt bridges on a hydrophobic interface (Fig. 3e,f and Supplementary Table 13). The two salt bridges are between Arg176 of α3-α4 and Glu51 of the CsRbcL, and between Lys177 of α3-α4 and Asp86 of the third beta-sheet (βC) in CsRbcL (Extended Data Fig. 15j). Phe173 of α3-α4 dominates the hydrophobic interaction in a pocket formed by Ile87 and Tyr97 of the CsRbcL. As the α3-α4 fragment contains a single residue difference between α3 and α4 (Fig. 1g), the residue at helix position 7 could be either a Leu (174) or Met (240). The map resolution did not allow distinction between the two, though when Leu174 was built into this density, it was also positioned to contribute to the hydrophobic interaction. We validated the binding interface by site-directed mutagenesis (SDM) of the α3-α4 fragment; selecting substitutions that disrupted the hydrophobic and salt bridge interfaces. In both cases, disruption of α3-α4-CsRubisco complex formation by native PAGE was observed (Fig. 3g, Extended Data Fig. 16c).

### CsLinker complements pyrenoid formation in *Chlamydomonas*

The Rubisco large subunit (RbcL) is highly conserved across the green lineage^36^, in contrast to the small subunit (RbcS) which shows much higher sequence variation^37^ (Extended Data Fig. 17). Given the interaction of CsLinker with the RbcL (Fig. 3c), we wondered whether this affords CsLinker increased cross-reactivity for other Rubiscos relative to EPYC1, which binds the RbcS^32,38^. We first considered cross-reactivity of CsLinker between Chlorella and *Chlamydomonas* Rubiscos, where the CsLinker interaction interfaces are almost totally conserved (Fig. 4a). CsLinker phase separated *Chlamydomonas* Rubisco (CrRubisco) with a similar efficiency as its cognate CsRubisco (Fig. 4h), and with a similar efficiency as EPYC1 for its cognate CrRubisco (Fig. 4b, Extended Data Fig. 21a,d). By contrast, EPYC1 was unable to demix CsRubisco in the reciprocal experiment (Extended Data Fig. 21a,c), in line with the lack of conservation of EPYC1-interacting residues in CsRbcS (Extended Data Fig. 17). SPR experiments with α3 and α3-α4 showed comparable *K*_D_ values for CrRubisco as the cognate interaction (108 μM [CI_95_: 102–115 μM] and 1.30 μM [CI_95_: 1.16–1.46 μM] respectively; Extended Data Fig. 13). A similar α3-α4-CrRubisco complex was also observed by native PAGE, albeit with apparently less stability (Extended Data Fig. 18c). Together, these data are consistent with CsLinker binding the same conserved interface in both Rubiscos.

**Fig. 4.**
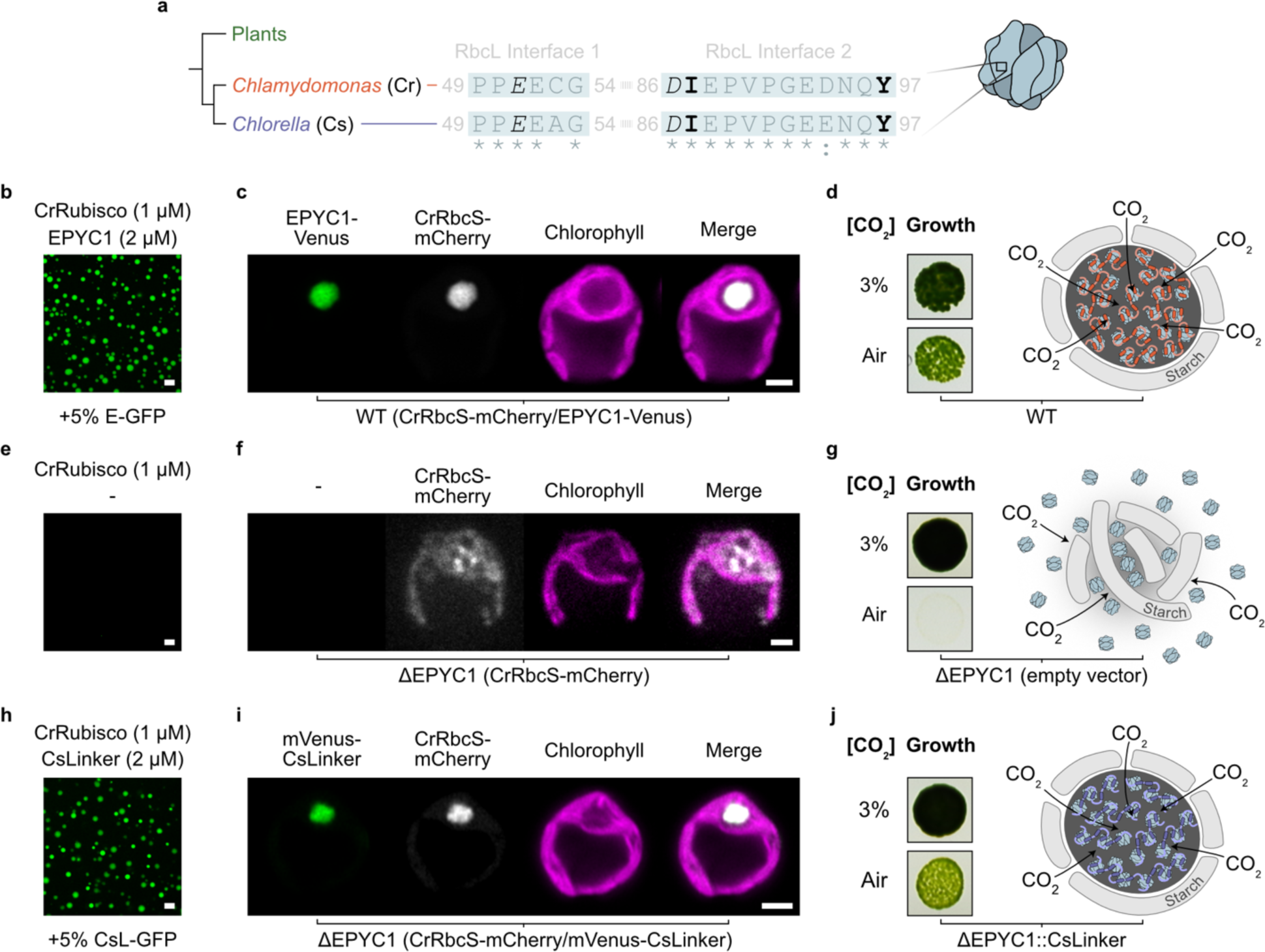
CsLinker can functionally replace EPYC1 in the *Chlamydomonas* pyrenoid. **a**, Alignment of the CsLinker-binding interface sequences from Chlorella (Cs) RbcL and the equivalent region of the *Chlamydomonas* (Cr) RbcL. Interacting residues are shown in black and stylized by interaction type according to Fig. 3f. Conserved residues are indicated by asterisks. **b**, Confocal fluorescence microscopy image of droplets formed with *Chlamydomonas* Rubisco (CrRubisco) and EPYC1, in which 5% (molar ratio) of the EPYC1 was GFP-tagged (E-GFP). Scale bar = 5 μm. **c**, Confocal fluorescence microscopy image of WT *Chlamydomonas* (CC-4533) expressing EPYC1-Venus and CrRbcS-mCherry. Scale bar = 2 μm. **d**, Growth phenotype of WT *Chlamydomonas* grown on TP minimal media under elevated (3%) and ambient (Air) levels of CO2 (left). Schematic representation of the pyrenoid (right). **e**, Confocal fluorescence microscopy image of CrRubisco alone, scale bar = 5 μm. **f**, Confocal fluorescence microscopy image of ΔEPYC1 *Chlamydomonas* strain expressing CrRbcS-mCherry. Scale bar = 2 μm. **g**, Growth phenotype of ΔEPYC1*Chlamydomonas* strain. Schematic representation of pyrenoid region (right). **h**, Droplets formed with *Chlamydomonas* Rubisco (CrRubisco) and CsLinker, in which 5% (molar ratio) of the CsLinker was GFP-tagged (CsL-GFP). Scale bar = 5 μm. **i**, Confocal fluorescence microscopy image of ΔEPYC1 *Chlamydomonas* strain expressing CrRbcS-mCherry and mVenus-CsLinker. Scale bar = 2 μm. **j**, Growth phenotype of ΔEPYC1 *Chlamydomonas* strain complemented with untagged CsLinker. Schematic representation of pyrenoid (right).

LLPS of Rubisco in the *Chlamydomonas* pyrenoid is essential to its function, allowing growth at atmospheric CO_2_ levels^15^ (Fig. 4d,g). Given the *in vitro* cross-reactivity of CsLinker and the functional analogy with EPYC1, we next considered whether CsLinker could replace EPYC1 in the pyrenoid of *Chlamydomonas*. We utilized a previously characterized *Chlamydomonas* strain which lacks EPYC1 (ΔEPYC1) and accordingly does not form pyrenoids. This strain has Rubisco distributed throughout the chloroplast and exhibits reduced growth under ambient CO_2_ conditions (air)^15^ (Fig. 4f,g). We expressed mVenus-tagged CsLinker in the chloroplast of ΔEPYC1, and subsequently co-expressed mCherry-tagged CrRbcS in the resulting strain to create a ΔEPYC1 (CrRbcS-mCherry/mVenus-CsLinker) line. In line with our observations *in vitro* (Fig. 4h), mVenus-CsLinker expression led to the formation of a micron-scale condensate at the canonical pyrenoid position that contained both CrRbcS and CsLinker (Fig. 4i). Although Rubisco partitioning and condensate size was reduced compared with WT, it was significantly increased over the background ΔEPYC1 strain, suggesting *in vivo* condensation of CrRubisco by CsLinker (Extended Data Fig. 19). To avoid any phenotypic impact of tagging either CsLinker or CrRubisco, we expressed untagged CsLinker in the ΔEPYC1 background and confirmed expression by western blotting (Extended Data Fig. 19i). In line with the visual recovery of Rubisco condensation by mVenus-CsLinker, introduction of untagged CsLinker restored the growth of the resulting ΔEPYC1::CsLinker strain in air to almost wild-type levels (Fig. 4j and Extended Data Fig. 20). The functional complementation of EPYC1 by CsLinker presents a compelling example of functional LLPS-driven organelle assembly complementation by a protein with little sequence similarity and a different binding interface, across an approximately 800 My evolutionary gap.

### CsLinker condenses native plant Rubisco *in vitro* and *in planta*

Encouraged by the functional cross-reactivity of CsLinker we observed with *Chlamydomonas* Rubisco, we sought to understand the extent of cross-reactivity for Rubiscos in the green lineage. We next demonstrated CsLinker was able to demix Rubisco from the multicellular Ulvophyte seaweed, *Ulva mutabilis* (Um), which retains all four CsLinker-interacting residues (Fig. 5a). Notably, the efficiency of phase separation was lower (Extended Data Fig. 21), SPR experiments demonstrated higher *K*_D_ values of the α3 and α3-α4 fragments for UmRubisco (162 μM [CI_95_: 158–167 μM] and 1.56 μM [CI_95_: 1.40–1.72 μM] respectively; Extended Data Fig. 13d), and native PAGE assays showed little gel-shift (Extended Data Fig. 18f). We attribute the reduced affinity and concomitant phase separation efficiency to the reduction of the *Ulva* RbcL interface 2 by one residue (Fig. 5a). This change would disrupt a potential hydrogen bond network with the Gln residue of the CsLinker helices, though the resolution in this region of the α3-α4-CsRubisco complex map did not allow distinctive assignment of this network (Extended Data Fig. 15k, 22c).

**Fig. 5.**
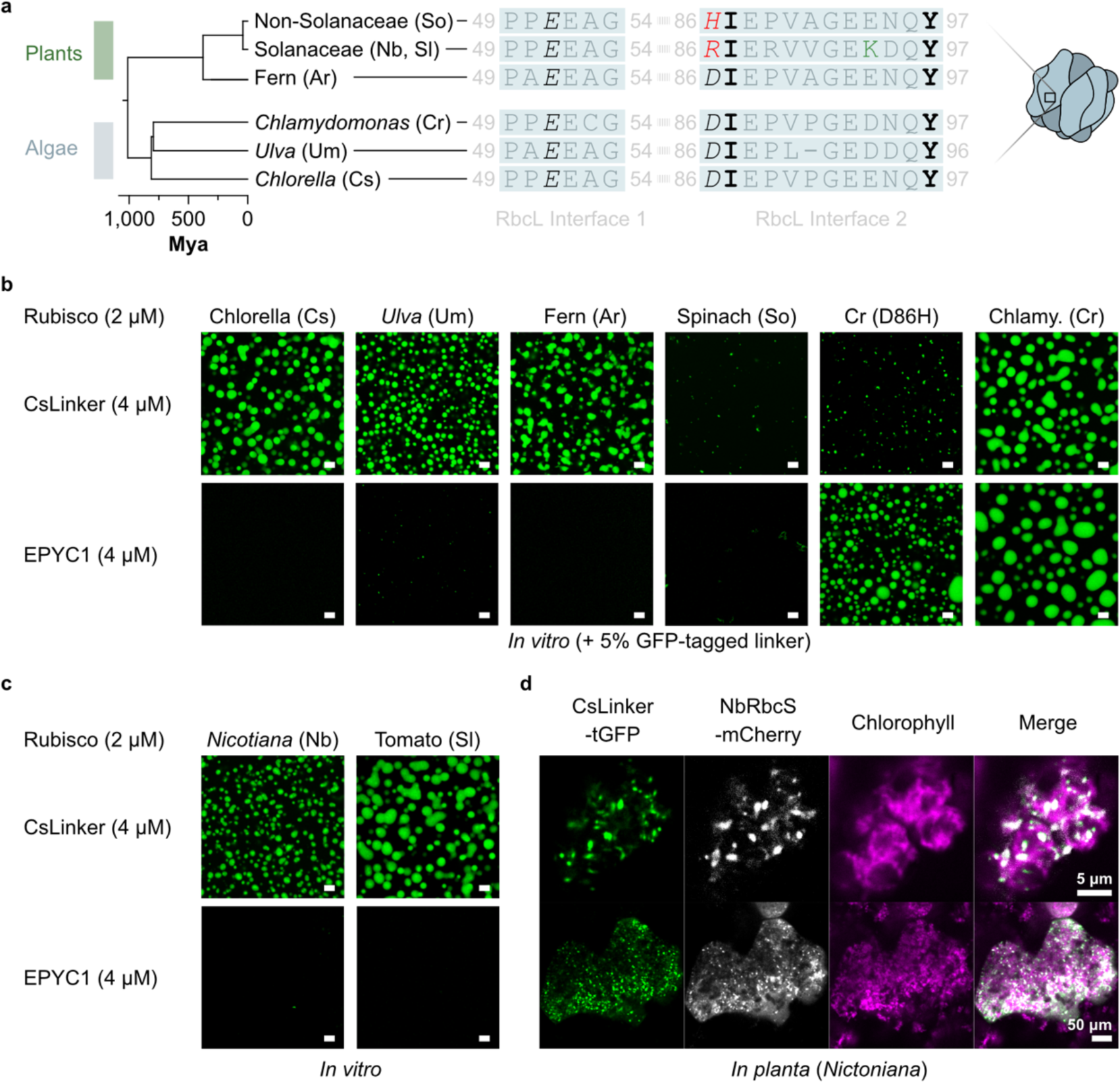
CsLinker condenses native plant Rubisco *in vitro* and *in planta*. **a**, Alignment of the CsLinker-binding interface sequences from algal and plant RbcLs. Interacting residues are shown in black and stylized by interaction type according to Fig. 3f. Substitutions of interacting residues are shown in red. **b**, Confocal fluorescence microscopy images of droplets formed with dilerent Rubiscos and either CsLinker or EPYC1. Scale bars = 5 μm. **c**, Images of droplets formed with Solanaceae Rubisco and either CsLinker or EPYC1. **d**, Confocal fluorescence microscopy images of CsLinker-tGFP and NbRbcS-mCherry transiently expressed in *Nicotiana benthamiana* chloroplasts, condensed into Rubisco puncta *in planta*.

Motivated by the wider goal of engineering pyrenoids in C_3_ angiosperm crop plants to enhance CO_2_-fixation^18,19^, we next wondered whether the cross-reactivity of CsLinker extends to plant Rubiscos. We analyzed RbcL sequences from major plant groups for conservation of CsLinker-interacting residues and found early diverged, non-flowering ferns were phylogenetically the closest plant group to angiosperms to contain all four key residues (Fig. 5a, Extended Data Fig. 22a). Accordingly, Rubisco from the fern *Adiantum raddianum* (Ar) demonstrated phase-separation with CsLinker *in vitro* (Fig. 5b), with a similar efficiency to the cognate pairing (Extended Data Fig. 21). Most angiosperm crops have a substitution of Asp86 for His86 in their RbcLs (Extended Data Fig. 22b). We tested the effect of this substitution (D86H) using Rubisco purified from Spinach (*Spinacia oleracea*; So), which has RbcL interfaces representative of the consensus angiosperm sequence (Extended Data Fig. 22a,b). We observed sub-micron aggregates of Spinach Rubisco at the same concentrations used for the cognate CsLinker interaction, but never observed droplet formation (Fig. 5b and Extended Data Fig. 21). To confirm this effect was predominated by the D86H substitution, we made the same substitution in the RbcL of CrRubisco that we previously demonstrated phase separates with CsLinker (Fig. 4h, Fig. 5b). The D86H mutation disrupted droplet formation of CrRubisco, which instead demonstrated similar behavior to the Spinach Rubisco (Fig. 5b). Notably, the D86H mutated CrRubisco still phase-separated with its cognate linker EPYC1 (Fig. 5b). Complex formation of D86H CrRubisco with α3-α4 was also disrupted, as assessed by native PAGE (Extended Data Fig. 18i). These data suggest substitution of the D86 residue severely impacts the affinity and phase separation propensity of CsLinker for Rubiscos of most angiosperm plants which possess a His in this position, including most of the key C_3_ crop plants (rice, wheat, soybean) (Extended Data Fig. 22b).

Amongst angiosperm RbcL sequences, there is some variation in the sequences of the CsLinker-interacting interfaces (Extended Data Fig. 22b). RbcLs of the nightshade family (Solanaceae), which contains some of the most widely consumed plants (potato, tomato, eggplant, pepper, tobacco), possess a distinct sequence composition at interface 2, due to co-evolution with their specific Rubisco activase which also binds this region^39^. Solanaceae RbcLs possess an Arg at position 86 whilst retaining the other three key interacting residues, but also display other unique sequence features in interface 2 (Fig. 5a). Given the charge inversion at position 86 (D86R) and presumed subsequent salt bridge disruption (Fig. 3e), we predicted Solanaceae Rubiscos would not phase separate with CsLinker. Surprisingly, Rubisco from the model Solanaceous plant *Nicotiana benthamiana* (NbRubisco; Tobacco) was readily phase-separated *in* v*itro* (Fig. 5c). As this behavior was specific to CsLinker (and not EPYC1), we hypothesized the interaction between CsLinker and NbRbcL is based on a similar binding interface as CsRbcL. Analysis of the Rubisco structure from *Nicotiana tabacum* (PDB: 1EJ7^40^) indicated that Lys94 of the RbcL (green in Fig. 5a) is favorably positioned to form a salt bridge with Glu169 of the helix of CsLinker (Extended Data Fig. 22e), that could compensate for the loss of the D86 salt bridge. Given that all key nightshade plants have conserved CsLinker-interacting interfaces (Extended Data Fig. 22b), we were interested whether CsLinker could phase separate Rubisco from other widely consumed members of this family. In line with the results of *Nicotiana*, Tomato Rubisco (*Solanum lycopersicum*) readily demixed with CsLinker *in vitro* (Fig. 5c and Extended Data Fig. 21). Finally, to explore if the *in vitro* observation of Solanaceae Rubisco condensation extended *in planta*, we transiently co-expressed TurboGFP-tagged CsLinker and mCherry-tagged NbRbcS in *Nicotiana benthamiana.* Strikingly, micron-scale condensates were observed in each chloroplast, which contained both Rubisco and CsLinker, representing the first phase separation of native plant Rubisco *in planta* (Fig. 5d and Extended Data Fig. 23). The condensation of native plant Rubisco represents a significant step forward in the goal of engineering pyrenoids in plants and provides prospect that ongoing engineering of CsLinker could allow phase separation of native non-Solanaceae crop Rubiscos.

## Discussion

Our in-depth characterization of a pyrenoid linker protein from Chlorella has yielded significant insight into LLPS driven organelle assembly and provided exciting frontiers for future plant pyrenoid engineering approaches to enhance photosynthesis.

Identification that CsLinker binds to the RbcL and structural characterization of the RbcL binding site has allowed us to make three general observations. Firstly, green lineage pyrenoid linker proteins likely evolved separately and convergently, as previously proposed^15,32,41^, and their binding region was not constrained to the small subunit of Rubisco. Their different sequence composition, binding affinity and binding region suggests the general physical properties of linkers also likely converged at linker-specific optima, which is supported by ongoing modelling efforts^42,43^. Secondly, given that Chlorella and *Chlamydomonas* diverged approximately 800 Mya, the capacity to functionally exchange EPYC1 for CsLinker in the *Chlamydomonas* pyrenoid provides a compelling example that the conservation of functionally critical physicochemical properties in intrinsically disordered proteins can be more important than conservation of a specific primary sequence. This example showcases the algal pyrenoid as a tractable model to explore the evolution of biomolecular condensation and LLPS evolution *in vitro* and *in vivo* with clear biological fitness readouts – often a limitation in LLPS systems^31^. We anticipate that our work will enable future studies to systematically test how the physical properties of phase-separating proteins (e.g. sticker number, sticker binding affinity, spacer length and spacer flexibility^44^) impact biological fitness. Thirdly, the high conservation of the CsLinker binding site in the green lineages RbcLs, enabled cross-reactivity of CsLinker with plant Rubiscos and allowed us to demonstrate phase separation of unmodified plant Rubisco for the first time. This finding overcomes a major future barrier for pyrenoid engineering in plants, which is currently dependent on genetic replacement of the multiple host plant RbcS proteins with those of *Chlamydomonas*^23,24,45^.

Whilst our results are encouraging, we acknowledge several areas that will require future work to address. Firstly, we presume that our demonstration of EPYC1 replacement in *Chlamydomonas* is dependent on retention of the native RbcS which is responsible for organizing other essential pyrenoid features through interaction with Rubisco-binding motif (RBM)-containing proteins^21,46^. Future approaches to engineer functional pyrenoids in plants without Rubisco engineering will be dependent on the development of approaches that circumvent RbcS interaction entirely, either by replacing RBMs in *Chlamydomonas* proteins or by identifying and characterizing analogous RbcL-binding parts, possibly from Chlorella. Secondly, although *in vivo* expression of CsLinker in both *Chlamydomonas* and *Nicotiana* resulted in Rubisco condensation, the partitioning of Rubisco in the condensate was lower than WT and multiple condensates were observed *in planta*. Whilst the level of CsLinker expression likely plays an important role, these observations also suggest a degree of tunability in the phase separation of Rubisco that is dependent on specific features of the interaction proteins when the interface is not fully conserved. Future studies could likely exploit this tunability to expand the cross-functionality of CsLinker with other non-Solanaceae Rubiscos, making use of the molecular details of the interaction outlined here.

Moving forward, as additional pyrenoids and their corresponding assembly proteins from diverse algae are characterized we expect our understanding of their evolution, the underlying principles of pyrenoid assembly and our ability to model pyrenoid systems to rapidly advance. We envision that this knowledge will provide an expanded parts list for pyrenoids and give us new tools to predict and modulate pyrenoid properties and thereby accelerate future efforts to engineer pyrenoids in plants.

## Methods

### Strains and culture conditions

*Chlorella sorokiniana* UTEX1230 (SAG 211-8k), *Chlamydomonas reinhardtii* WT (CC-4533), ΔEPYC1 (CC-5360) and resulting strains were maintained on 1.5% agar Tris-acetate-phosphate (TAP) medium with revised trace elements47 plates in low light (∼10 μmol photons m-2 s-1). For Rubisco extraction, growth, immunoelectron microscopy and confocal microscopy experiments, strains were grown in Erlenmeyer flasks in Tris-phosphate (TP) media in medium light (∼50 μmol photons m-2 s-1) under ambient CO2 to a density of ∼0.5-1x10^7^ cells mL^-1^.

### FLIPPer and bioinformatic analysis

FLIPPer was originally built using IUPred2A48 as the disorder prediction and filtering software, which was used in the initial identification of CsLinker. For licensing reasons, the pipeline was subsequently rebuilt using metapredict V249 instead and the outputs were largely unchanged. FLIPPer is available (github.com/james-r-barrett/FLIPPer) and was used with default settings thresholded based on EPYC1 sequence analysis. Briefly, input sequences are physicochemically filtered, repeats detected with XSTREAM50 and filtered to contain interacting residues before disorder prediction and filtering with metapredict. Output sequence structures were manually examined for repeated helical regions by AlphaFold 251 prediction in ColabFold v1.552. Dilerential gene expression analysis was completed according to ref.53, using publicly available data (PRJNA343632).

### Co-Immunoprecipitation Mass Spectrometry

Polyclonal rabbit antibodies were raised to Rubisco (EVWKEIKFEFETIDTL-cooh) and CsLinker (PTPVSNSGVRSAMSSG-amide) peptides (YenZym Antibodies LLC, USA). The control rabbit antibody was raised to *Chlamydomonas* BST2 (PDLDSINAAAPNGNGSHNGN-amide). 1x10^9^ Chlorella cells grown in TP sparged with 0.01% CO2 were lysed by ultrasonication in 10 mL IP buler (20 mM Tris-HCl pH 8.0, 50 mM NaCl, 0.1 mM EDTA, 12.5% Glycerol (w/v), 5 mM DTT, 1x cOmplete protease inhibitor tablet / 50 mL). 1 mL of clarified lysate (30 minutes at 50,000 *g*) was applied to 200 μL of protein A Dynabeads™ (Invitrogen) loaded with 32 μg of respective antibodies that had been blocked with BSA (2 mg mL-1) for 1 hour and washed 3 times with IP buler. The lysate was incubated for 3 hours before washing. Bound protein was trypsin-digested from the beads, post reduced with DTE, and alkylated with iodoacetamide prior to UPLC separation by 60SPD EvoSep One (EvoSep, Denmark) method and acquisition by PASEF-DIA using a timsTOF HT (Bruker, USA). Data were searched using DIA-NN54 and filtered to 1% FDR with a minimum of two unique peptides.

### Immunoelectron Microscopy

Cells were fixed in 1% glutaraldehyde, 2% formaldehyde in 0.1 M sodium cacodylate (pH 7.4) for 1 hour before resuspension to 2% (w/v) agar. Agar blocks were dehydrated using a 10-50% ethanol gradient at 4°C, followed by 70-100% at -20°C, then infiltrated with LR White resin containing 0.5% benzoin methyl ether (London Resin Company, UK) and polymerised in gelatin capsules for 24 hours at -20°C and 24 hours at -10°C under UV light. 70 nm sections were cut using a Leica UCT7 ultramicrotome with a Diatome knife and mounted on nickel grids. All imaging was completed using an FEI Tecnai T12 BioTWIN TEM operating at 120kV with Ceta CCD camera.

For immunolabelling, grids were blocked with 3% BSA (w/v) in PBS for 30 minutes prior to primary incubation with either purified antibody (0.09 μg mL-1) or pre-immune serum (1:5,000 dilution) for 1 hour at 30°C in a humidity chamber. Secondary incubation was completed with a 1:40 dilution of Goat anti-Rabbit IgG 10 nm gold conjugate for a further hour (Merck).

### Absolute quantification mass spectrometry

6x10^6^ Chlorella cells grown in TP sparged with 0.01% CO2 were boiled in 50 μL of Laemmli buler in triplicate for the biological samples. The standards were separately boiled in duplicate in the same volume. Both the samples and standards were trypsin-digested from SDS-PAGE gels, post reduced with DTE, and alkylated with iodoacetamide prior to separation by UPLC with a 25 cm PepMap column (ThermoFisher Scientific) and 1 hour acquisition by DDA using an Orbitrap Fusion Lumos Tribrid (ThermoFisher Scientific). Technical replicate injections were used for the biological samples, after which the values were averaged. LC-MS chromatograms were aligned using Progenesis QI and the MS2 spectrum was searched using Mascot with a 1% FDR. Matches were mapped to MS1 intensity using Progenesis QI and summed at protein level. The external calibration curve was used to calculate protein abundance in the biological samples (Extended Data Fig. 7).

### Cloning, plasmids, and strains

For *E. coli*, the mature CsLinker sequence was predicted by TargetP 2.055, codon-optimised in Geneious Prime using the *E. coli* K-12 codon usage table and synthesised (TWIST Bioscience). The sequence was ligation-independent cloned (LIC) into pET His6 GFP TEV LIC cloning vector (addgene #29663) to produce the His-mEGFP-TEV-CsLinker plasmid. The mEGFP-CsLinker fusion was subsequently PCR-amplified and LIC cloned into pET MBP His6 LIC cloning vector (addgene #37237) to produce His-mEGFP-TEV-CsLinker-TEV-MBP-His plasmid. α3 (residues 142-207) and α3-α4 (residues 142-272) fragments were PCR-amplified from this plasmid and Gibson assembled back into the PCR-amplified backbone to produce His-mEGFP-TEV-α3-TEV-MBP-His and His-mEGFP-TEV-α3-α4-TEV-MBP-His respectively. SDM constructs were constructed by introducing nucleotide exchanges in the primers used to amplify the α3-α4 fragment from His-mEGFP-TEV-α3-α4-TEV-MBP-His. The SDM sequences were Gibson-assembled back into the backbone to create His-mEGFP-TEV-α3(F173DL174D)-α4(F239DM240D)-TEV-MBP-His and His-mEGFP-TEV-α3(E169AR176A)-α4(E235AR242A)-TEV-MBP-His constructs.

For *Chlamydomonas* chloroplast expression, the mature CsLinker was codon optimised in Geneious Prime using the *Chlamydomonas* chloroplast codon usage table (Kazusa) and synthesised (TWIST Bioscience). The mVenus sequence was codon optimised using Codon Usage Optimizer v0.92 (https://github.com/khai-/CUO). Sequences were PCR-amplified and golden gate assembled into pME_Cp_2_098 (Gift from René Inckemann and Tobias Erb). The ΔRbcL plasmid was created by Gibson assembly of the aadA gene amplified from CC-5168 into plasmid P-67 cpDNA EcoRI 14 (Chlamydomonas Resource Centre) to replace the RbcL CDS in frame. The RbcL_D86H plasmid was created by KLD site-directed mutagenesis (New England Biolabs) of the P-67 cpDNA EcoRI 14 plasmid (Chlamydomonas Resource Centre).

For *Nicotiana benthamiana* transient expression, the mature CsLinker sequence was codon optimised and synthesised using GeneArt and the *Nicotiana benthamiana* codon usage table (ThermoFisher Scientific). The sequence was golden gate assembled into pICH47732 with a 35S promoter, AtRbcS1A transit peptide, C-terminal Turbo-GFP and HSP+NOS double terminator according to 56. The NbRbcS was synthesised with AtRbcS1A transit peptide by IDT and assembled into pICH47751 with AtRbcS1A promoter, C-terminal mCherry tag and OCS terminator.

### Rubisco Extraction

Rubisco was purified from *Chlamydomonas* and Chlorella according to ref.57 with the addition of a 16.5 hour, 37,000 rpm 10-30% sucrose gradient ultra-centrifugation step performed in an SW41-Ti rotor prior to anion exchange using a HiTrap 5 mL Q XL column (Cytiva). The same method was used for *Ulva*, Spinach, *Adiantum*, *Nicotiana* and Tomato, except lysis was completed by manual agitation in a blender. Rubisco was labelled using an Atto594 protein labelling kit (Sigma-Aldrich), according to manufacturer’s instructions.

### *E. coli* protein purification

All constructs were purified from *E. coli* BL21 (DE3) strains harbouring respective plasmids. Cells were grown to OD600 of 0.5-0.8 in Luria Broth or ^15^NH4Cl minimal media for the NMR samples, before induction with 1 mM isopropyl β-d-1-thiogalactopyranoside (IPTG) for 3 h at 37 °C. Pellets were snap-frozen before ultrasonic lysis in high salt buler (50 mM Tris-HCl, 500 mM NaCl, 25 mM imidazole, pH 8.0) with 2 mM phenylmethanesulfonylfluoride (PMSF). Soluble protein was applied to an IMAC column (HisTrap™ FF Crude 5 mL - Cytiva), washed with high salt buler and eluted with a linear gradient to 500 mM Imidazole in high salt buler. For untagged CsLinker, α3, α3-α4, α3(F173DL174D)-α4(F239DM240D) and α3(E169AR176A)-α4(E235AR242A) constructs, the N-terminal mEGFP and C-terminal MBP were cleaved overnight with TEV protease produced according to refs.58. The cleaved solution was passed over an IMAC column equilibrated with high salt buler and the untagged flow-through was collected. Size-exclusion chromatography (SEC) was completed on the flow-through using a HiLoad® 16/600 Superdex® 75 pg (Cytiva) equilibrated with 50 mM Tris-HCl, 500 mM NaCl, pH 8.0 buler. For the mEGFP-CsLinker, no TEV cleavage or second IMAC was completed, but protein was exposed to SEC.

### Droplet sedimentation assay

All assays were completed in 5 μL reaction volumes in which Rubisco and linker were mixed and incubated at room temperature for 15 minutes. Sedimentation was performed at 10,000 g for 10 minutes. Band intensity was quantified in Fiji.

### *In vitro* confocal microscopy and fluorescence recovery after photobleaching (FRAP)

All *in vitro* confocal and FRAP experiments were completed on either a Zeiss LSM880 or Zeiss LSM980 confocal microscope with a 63x 1.4 numerical aperture (NA) Plan-Apo oil-immersion lens (Carl Zeiss, Germany) in 5 μL reaction volumes formed in µ-Slide 15 Well 3D coverslips (ibidi, Germany). For FRAP experiments, the sample volumes were overlaid with 30 μL of ibidi Anti-Evaporation oil.

FRAP experiments were completed on droplets with diameter ∼1 μM, where half of the droplet was bleached in half FRAP experiments. 15 pre-bleach images were taken before bleaching (100% 488 nm intensity, 1 cycle for mEGFP, 100% 488 nm + 561 nm intensity, 5 cycles for Atto594). Bleach depth was consistently 60-75%. FRAP images were processed in Fiji. Briefly, fluorescence images were translationally stabilised using the Image Stabilizer plugin (4 pyramid levels, 0.99 template update coelicient) output of the brightfield images and the mean gray value in the bleached, unbleached and background ROIs was measured. Background values were subtracted from bleached and unbleached values before photobleach normalisation using the unbleached references was completed. Full-scale normalisation was completed using the average pre-bleach intensity. An exponential model was fitted to the post-bleach data (y(t) = A * (1-e-kt), where A is the plateau, k is fitted and t is post-bleach time).

### Native PAGE gel-shift assay

Rubisco and fragments were mixed in 5 μL reaction volumes and incubated at room temperature for 30 minutes. 1.6 μL of loading buler (80 mM Tris-HCl pH 8.0, 200 mM NaCl, 40% glycerol) was added prior to loading onto 4–20% Mini-PROTEAN® TGX™ gels. Electrophoresis was completed for 4 hours at 100 V at 4 °C.

### Surface plasmon resonance (SPR)

All SPR experiments were completed in triplicate on a Biacore™ T100 fitted with a T200 upgrade kit with a sensor temperature of 25 °C. Immobilisation of Rubisco was completed according to ref.32 and experiments were completed with modifications. The analyte was injected at 15 μL min-1 for 30 seconds followed by 360 seconds of dissociation. After each replicate set the chip was washed with 1M NaCl in running buler, after which the chip was washed for 360 seconds with running buler. Binding to the reference chip was negligible. Fitting of the reference-subtracted curves was completed using hill equation (y(x) = *B*max*x / (*K*D + x), where *B*max is the maximum specific binding, *K*D is the apparent dissociation constant, and x is the analyte concentration.

### Single-particle cryogenic electron microscopy data collection and image processing

Chlorella Rubisco and the α3-α4 fragment were mixed in a buler compatible with cryo-EM experiments (200 mM sorbitol, 50 mM HEPES, 50 mM KOAc, 2 mM Mg(OAc)2·4H2O and 1 mM CaCl2 at pH 6.8) at final concentrations of 0.5 μM and 16 μM respectively and incubated at room temperature for 10 minutes. α3-α4 had the same apparent *K*0.5 for CsRubisco in this buler as the Tris buler used for native PAGE experiments (Extended Data Fig. 12f). 2.5 μL of solution was applied to R1.2/1.3 Cu 400-mesh grids (Quantifoil, Germany) that had been glow-discharged for 60 seconds with a current of 15 mA in a PELCO easiGlow system. Using an FEI Mark IV Vitrobot (ThermoFisher Scientific) with chamber at 4 °C and 95% relative humidity, the grids were blotted for 8 seconds with a blot force of -5 before rapid plunge-freezing in liquid ethane.

Data was collected on a 200 kV Glacios cryo-electron microscope equipped with a Falcon IV direct electron detector at the University of York. Automated data collection was performed using EPU in AFIS mode (ThermoFisher Scientific). A nominal magnification of 240,000x and electron fluence of 50 (e-/Å2) with a calibrated pixel size of 0.574 Å2 was used during collection in which each exposure was 6.52 seconds. The 100 μm objective aperture was inserted and the C2 aperture was 50 μm. A range of defocus value were used (-1.8, -1.6, -1.4, -1.2, -1.0, -0.8 μm).

Relion3 was used for processing and 3D reconstruction59,60. Of the 1568 EER frames, 32 were grouped to give a fluence/frame of 1.02 e-/Å2. Relion’s implementation of MotionCor2 was used for motion correction prior to CTF estimation with CTFFIND4, assuming a spherical aberration of 2.7 mm61. Initially 465 particles were manually picked and reference-free 2D classification was completed. 237,035 particles were autopicked using selected 2D class averages. Particles were extracted with 2x binning and 2D classification was re-completed. 2D classes presenting clear Rubisco structure were selected and a C1 symmetry 3D classification was completed. A single 3D class with clear secondary structures was used for auto-refinement with D4 symmetry (73,962 particles). After CTF refinement and Bayesian polishing in Relion62 a 20 Å low-pass filtered mask of the 3D refined map, expanded by 10 pixels with a soft edge of 6 pixels was used for solvent masking and resolution estimation. This map has D4 symmetry and represented the CsRubisco holoenzyme (EMD-18049).

In the 3D class used for the D4 map, additional low-resolution density was observed on the equator of Rubisco, suggesting a sub-stoichiometrically bound partner (Extended Data Fig. 14d). The predicted helix of α3 (Fig. 1f) was built into one region of additional density in UCSF Chimera (Extended Data Fig. 15a)63. The soft, featureless mask was created from these coordinates by lowpass filtering to 20 Å, extension by 5 pixels and softening by 6 pixels. This mask was used for two rounds of C1 3D classification with a D4 symmetry expanded particle dataset created from the 73,962 polished particles used for the D4 map (591,696 elective particles). 133,171 symmetry expanded particles were used for the 3D reconstruction of the C1 CsRubisco-α3-α4 map, which was processed as for the D4 map (EMB-18050).

### Single-particle cryogenic electron microscopy model building, fitting and refinement

A holoenzyme model of Chlorella Rubisco was built in UCSF Chimera using AlphaFold 2 structural prediction of the CsRbcL and CsRbcS sequences from *Chlorella sorokiniana* UTEX1230. This model was rigid-body fitted into the CsRubisco holoenzyme map using UCSF Chimera. Flexible fitting was performed in COOT 0.964, using one of each CsRbcL and CsRbcS chain. Real-space refinement was completed in Phenix65. The coordinates of this refinement were applied to the other 7 CsRbcL/CsRbcS chains. For α3-α4 model building, the AlphaFold 2 predicted helix of α3 was manually built into the additional density present in the C1 CsRubisco-α3-α4 map in COOT. The side chains of residue E169, R171, E172, E175 and R179 were removed prior to refinement in Phenix, due to a lack of supporting density. Both the structures derived from the C1 and D4 maps were validated using MolProbity66. Figures were created in UCSF Chimera and UCSF ChimeraX67.

### NMR Spectroscopy

Spectra were recorded at 10 °C on a Bruker Advance Neo 700Mhz spectrometer equipped with a TCI Prodigy CryoProbe (Bruker). Samples were analysed in buler containing 15 mM sodium phosphate pH 8.0, 150 mM NaCl and 5% D2O. Protein concentrations were 920 µM for α3 and 500 μM for α3-α4. Spectra were processed using TopSpin (Bruker) and analysed using CCPN Analysis68.

### *Chlamydomonas* Transformation

Chloroplast transformations were completed by particle bombardment with a Biolistic PDS-1000/He Particle Delivery System (Bio-Rad). Per bombardment, 0.5 mg of 550 nm gold nanoparticles (Seashell Technologies Inc.) were incubated with 1 μg of plasmid DNA and prepared according to manufacturer’s instructions. 1x107 cells were plated in a 4 cm diameter on TAP plates and placed ∼9 cm below a 1100 psi rupture disk. After firing under vacuum conditions, cells were recovered for ∼24 hours before re-plating to selection conditions. Transformants into the ΔEPYC1 strain for RbcL knockout and CsLinker reintroduction were re-plated to TAP plates containing 100 μg mL-1 spectinomycin under low light (∼10 μmol photons m-2 s-1). Transformants from the ΔEPYC1ΔRbcL complementation with D86H mutated CrRbcL were plated on TP plates and recovered in 3% CO2-air mix at ∼50 μmol photons m-2 s-1. 16 transformants from each transformation were propagated 4 times on selection plates before checking for homoplasmic integration of the genetic material.

Transformation of mCherry-tagged *Chlamydomonas* RbcS was completed with pLM03515, using a NEPA21 electroporator according to ref.69.

### Chlamydomonas growth assays

Spot test assays were completed according to ref.15.

### *Chlamydomonas* confocal microscopy

All images were captured on a Zeiss LSM880 confocal microscope in Airyscan mode with a 63x 1.4 numerical aperture (NA) Plan-Apo oil-immersion lens (Carl Zeiss, Germany). µ-Slide 18 Well chambered coverslips (ibidi, Germany) with 10 μL of cell suspension and 30 μL of 1% TP-low-melting point agarose were used for imaging.

### Transient expression *in planta*

Overnight cultures of electrocompetent GV3101 Agrobacterium harbouring the relevant plasmids were grown overnight in LB. Cultures were resuspended in 10 mM MgCl2 to OD600 of 0.8 and syringe infiltrated into the youngest fully expanded leaves of four-week old *Nicotiana benthamiana* plants. Images were captured using a Leica SP8 confocal microscope 48 hours after infiltration and incubation of plants at 25 °C, 16h light 8h dark, 170 μM photons m-2 s-1.

## Data availability

Proteomics data were deposited in MassIVE, with ProteomeXchange identifier PXD044179. Electron density maps were deposited in EMDB with accession codes EMD-18049 (D4) and EMD-18050 (C1), and their corresponding coordinates in the PDB with accession codes 8Q04 and 8Q05 respectively. Code for the Fast Linker Identification Pipeline for Pyrenoids (FLIPPer) is available in a public GitHub repository (https://github.com/james-r-barrett/FLIPPer). All other associated data will be made available in a Zenodo repository.

## Acknowledgments

J.B. and S.M. were supported by BBSRC studentships (BB/M0111511a and BB/T007222/1, respectively). J.B., L.C.M.M. and A.J.M. were supported by a CTRF grant (AP23-1_023). M.I.S.N. was supported by an NWO Rubicon 2022-1 grant (Project ID: 019.221EN.010). Y.M. was funded by a postgraduate research scholarship from the Darwin Trust of Edinburgh. C.D., M.C.L, M.J.P. and L.C.M.M. were supported by EPSRC funding (EP/W024063/1) as part of the York Physics of Pyrenoids Project (YP3) who are also thanked for fruitful discussions. A.J.M. and L.C.M.M. were supported by BBSRC-NSF/BIO funding (BB/S015337/11 and BB/Y000323/1) and Bill and Melinda Gates Foundation/Foreign Commonwealth and Development Office funding (INV-054558). J.N.B was supported by a UKRI Future Leader Fellowship (MR/T040742/1), as was L.C.M.M. (MR/TO20679/1). In addition, we wish to thank University of York Bioscience Technology Facility staff Dr. Adam Dowle, Dr. Grant Calder, Dr. Andrew Leech and Dr. Clare Steele-King for their assistance in mass spectrometry, FRAP, SPR and immunoelectron experiments respectively. Philipp Girr is thanked for his mentorship and for supplying the TEM image of the *Chlamydomonas* pyrenoid. Tobias Wunder and Oliver Mueller-Cajar are thanked for their assistance in the conception of the project. René Inckemann and Tobias Erb are thanked for the gift of the pME_Cp_2_098 plasmid. Luke Oltrogge and David Savage are thanked for allowing the use of Rubisco graphics. Katy Davis is thanked for the supply of Solanaceae plant material. We would also like to thank the BBSRC, the Wellcome Trust (grant number 206161/Z/17/Z), Tony Wild, and the Wolfson Foundation for funding the Eleanor and Guy Dodson Building and associated Cryo-EM facilities, in addition to Dr. Johan Turkenburg and Sam Hart for coordinating data collection.

## Author contributions

J.B. and L.C.M.M. conceived the study and wrote the manuscript with contributions from M.J.P., A.J.M. and M.C.L.. J.B. performed biochemical characterisations, in vivo *Chlamydomonas* experiments. M.I.S.N. performed co-immunoprecipitation, absolute quantification and immunoelectron experiments. J.B conceived and constructed FLIPPer with assistance from S.M.. C.D. performed NMR experiments. A.S. performed sequence validation. J.B. and J.N.B. performed cryo-EM experiments and analysis. Y.M. and A.J.M. designed and performed in planta experiments.

