## Supplementary Figures and Tables for "A promiscuous mechanism to phase separate eukaryotic carbon fixation in the green lineage"

CSI2_123000008654

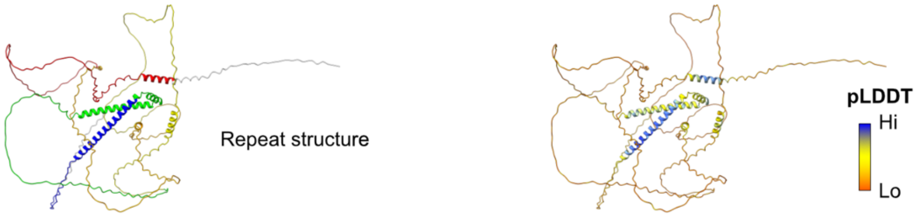


MSQCTIQHTEADAGLLSLAAPTSDAEGWLVPVAQPASRQERLSEQASLFMANLLAQAAPGTSNWLSAEPADPFAPFQQQTAAPPATSAQQRRQLTSEGSEVQPLPCVSLAGQPPRSQRRPSSVTAPQ
PRPGSALPRPASSFPRPTPKPAASASRPAMPAQAAASQAASPKPLWRPSGSAQPSPTTPRHSTPGTPTRQSLAGTPPAVRTAAAPRGLASTSQQAQQAQRPASAAAKQAPRAEELSPSTKVAQWLFS
PAHPAAPAGRLTVKRPASSAAAAAAAAASLASPSASSSEASWRIAAATGTGSTLSSPAGTPAAAAAAAAAASGFRRQASGSRIPRPGTAASAAAPAEPLPAEGTSPSARAAAPASLPTRRSELALKQ
LAQRALAAGSTSPPAPCPLHVRRPSSLRGAAQAASVAAAPASTSLPSSQPTSPFQQTFVSARSADAASVSSYLSAAESPAGETTPRRAHRLSSSSSSQPASSQPGSPAALPLSPRPGSAGSQLRQAQ
AAAVGSLAAAQAGARAAAEEVALLSTASPGAWAAAQAVAAVSPASDTSLLDAQLALLDRLSRDLSDASSQIAEVCILDTLANSRRASSSMPTARRLSLQLHSPGASSAGASSAGAAGSPPRPRSLAG
SLHGQHHAM

CSI2_123000010295

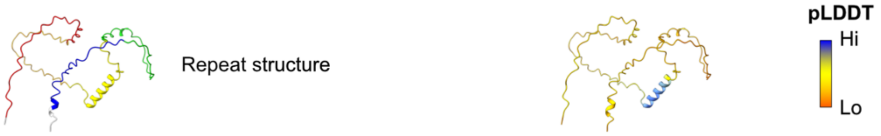


MARPRGKGSKPSTTQTAAAPSSRKPKGGRPRPASKAASGGGKPGGRPSGSSKPAQLAGAAHRIVEFAPPPGPATLAERFALLEKQRQQLGAGRQRAAAPALGVASGGVAKKAKQQQRRRKGKASSGE
AGGAGASGKPAAAKAQGAAAAGASSGSGKRRRRGGKRSQAAKMDTK

CSI2_123000010558

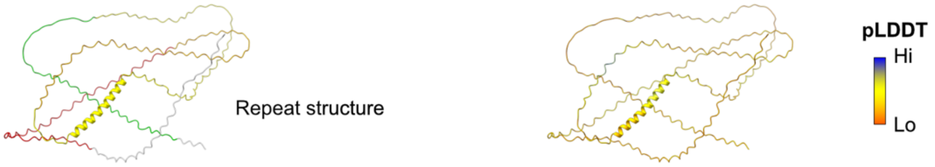


MREVAALANTSSPAREWAAAKTASPSSAAAGPPAAKRRLQDDAPQDFVIAAISGSAAQAVAPIEPAAPSQAAADAPVPPAAATPAAPAPTIPAPAAEEQSPPAVVEALFSDAASPAGTAAAAEAPGA
PPAQPARAASLNRSPAVSHAVRSLGPVAESSPAVPLPPLPTLAPVxxVAGSATSPAPARSISLTRQLSRKLSRGMSRLRASIRRRLRRADSPPVPRSRDLSGRISPTSGTAHLPAIPEGAPASAAAA
AAGSGGLGGQVELATLAPPHPAATAAVERAASVAAAPSPAAAAAPGAETAAAAGSQASRASLVEVDLSEQRPAPSSSPSRRRGLACACFA

CSI2_123000012064 (**CsLinker**)

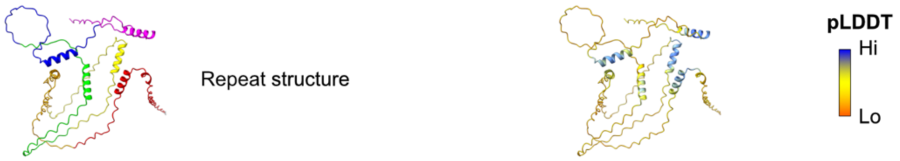


MQALSARVTLGRPATARVAKKQQRPLTVAMATAQLSAEQKAFLERKARESRSPAPVETRGRSAPRPRSASPVGRPSSRTPAPASYSAPAASYSSAPANGLSAEQREFMERKARESRSPAPVETRGRS
APRPRSASPRPTSRTPAPASYSAPAASYTAPAAAPAGGLSAEQREFLERKARESRSPAPVETRGRSAPRPRSASPRPTSRTPAPASYSAPAASYSAPAAAPAGGLSAEQREFMERKARESRSFSAPV
ETRGRSASARPSARPSSRTPTPVSNSGVRSAMSSGSSAGLSAEQREFLERKARESRSPAPAAASQSSSYGSAASGLSAEQREFLERKARESRSFSAPAPTRGRSSTPRRR

CSI2_123000002244

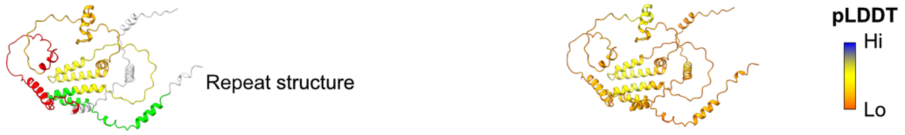


MASALRPSAACIARRQASDWNKTRVVPKSAFATALHAKASNARALPHSASSRIEAVHVEAALVELAAPAAFPPLPGTTTTRPATLAGAWAARQQAAASCAAVPAARGVSEDAEDVAPRLPAAYLPPA
RREGATATILAGCSLAYPAPAQPAQRPAAYVPPHSRTAEQQAAAAEACRPGRICAWPYLTAGPLPQPAAEAPLPLYRLALLDIDDSTEGNARMRLAGLLMLAFATMMHFLGSPQAPFLYLFAWMAGV
LGPSRAGTSEPHAPQPPPPLPPAPSDIADSSEGNAHMRLAGLLMLAFAMMMHFLGSPQAPFLYLFAWITHPQDEHVAHRCCAPEPERAAQRRHARAARPTAAAATAAGAQ

CSI2_123000002308

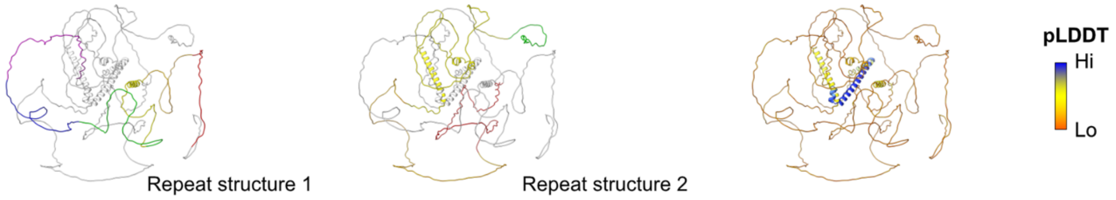


MQRGAPPRDCPEQQGGVAALAASPGARPAGPGAGTPCCPPSSPLQPIAEAGPAPAAPGCPAPALDAGAGVMRPPPPKCAPLSGSLATSSQREALTAQHSKLLSSEAATEERDEYLMRLLCKGAPPAG
VPPAPPAAQVPQGAPPPALQQQAVQAQAAQAVEAPRFRAPGPVSRSVPPSAPAPGSATPPAPSPSGLLPRATSTTLASPSSTVGAVTGTACATVPSLPLPAPSATATTTDQAVPSSGGAAAAPTTAA
PPPRAAPPLPPSARAAAAGAAAAVTALPAPQLVAVPLVAVPPELAATFDQTRRMLGDEIALSVRDTMIRQQVMFLEQLYDLHRAIAIQKLLMQNCPEVMGEAMRLMSSVGSQHGSGSGRGSKRQRTT
PNGGATVPGSGSGDPSSFRPDSQFAAVPQLPGEAQTRDDEDDDGGSGDANGSGSAGANGSGGGSTSPQQLVPSLPPPPPLPYMYLAHTAVATPSGASAAPVGWQWGMPPLVQGQVQQLQQAVGLPPV
PRGPPQMAPWVGQDAGLAAVGINTHTPQHPKHAAGAAGSKPSKTATAATAPPAPRATPPRPAAAQPSAAAAAAPAAPAPQPKQQQPVAQPQPQQAPLALAQAPAAPAMAQPAPASLAMPPPGMPPPM
PFGPPPPHxxxAxHGPPGLPCPAPGAADPMAWWYQNYYGSQQQMGGMSLATAQAAAAAAAAFMQQAAQQPGGTGCTTAGPLGTAGTAGGTGTGGPAVTKWWHDPQLTFGPPVDTEMLARRPAGGPAA
LLASAREGPEVQSAPAGQLKRPADSRRRSSMPYSDGPRMRLKRKATADADEASSGTTGHTHDPDGGAGTTAGGPFKKHESLRARSARTSPQGPAAADDPMGGGGQDENAAQLLLAIGKQ

CSI2_123000011700

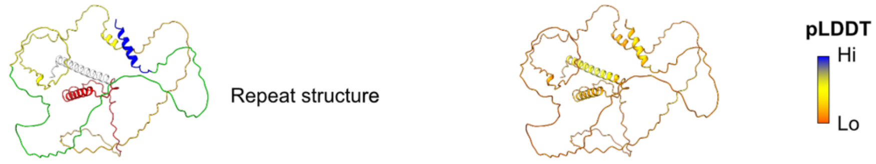


MAAPVTSSDATRNITLACIVAGTVLLVVFVAWKMRRRVRCSCSCWSLADPAAAERRRVLASAAVLYQAAAANPDFKLRLQPGPGRWPQPSPAASVYTSPTLAPPDPGSTLGTPASWRSRSPTGSYQA
ELAPVPAAPPLRSRTSPSGRPPLPPAPPRGPAQPMASPGTSPASSQQQRRSPPQRRSVRELVRAWSGGGSQNGRERSPPAVTAAAAAVAAADLEGQQLSATHEAQAFTWSQLGLSPAASPAASRRVS
ENAAAPYRLPLPPSQQGEVHSNPVFDWTAAGVAATDGGTLGPAAGSFSQQVKRTASRSRLSRGVSFAEPAGEGSAHGGREGGGHSRRGSFVTPEQGSRREAGGQMSSRPSEQALLQLHCELEGIQE

CSI2_123000004449

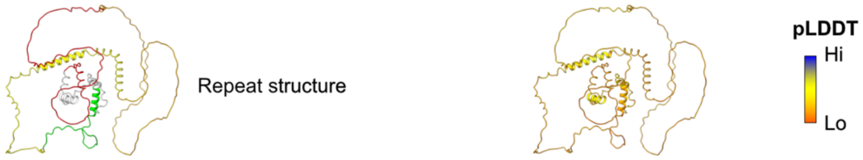


MASGRLALPPRQGGSAEQRKRVSDEAIADAVEKLSGAPVGAVERLAGAPPALTDLAAAAEAAAAGGSTGKKRKAEEASPPAAAAADGAATGERQTTGGRSRRSNAGKKPLRAEFDDDGDSWDDESEE
DSPPRGKRRGKAKADSDFEDFSEPESASESEEDSEDEFASEDEDSPPKCGKKAPARGKGAAGKSGAAAGKKSGAAAKKSGAAAAKKSAAAAKPPAKAPAAKKSAAAAKAAAPPKPAAKSGVGKKAAG
AAAAAAAAAAAAAEEAEQVPEEQQQLEQELPVQDTAQLAAAAAQQAEEEAAAEEEASAAVPPPAKKAKVPAPKPAAAPKPAAAPKPAAAPKPAAAAGGSKPVGLKAAGGGVRKPLAPAAQAKAPSPA
GGGPSPAGGGGLMAAAARVVQGAA

CSI2_123000004744

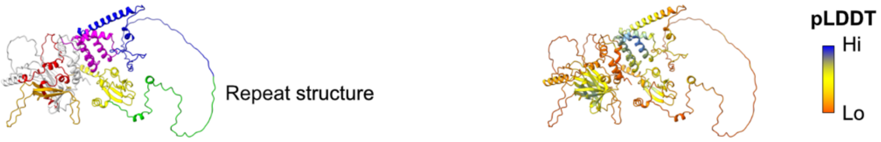


MVQPAAAAEALALQLQATELENILDARDKGRTPTILRRNLSYVDRSRRRALQQGHGAALQLQLELAGGPGPAGGPMQTSSAAAGSSSPDRATRHVLVAAVCIVAPQNGTQAYVPVSARTVRLPPERH
ARLSLCLPLGAASPAAGLAVVLLHPFAGDARWRYRYRLSNEEQRRWNAAQAAAAEGQHMQPPECGQLALLASLAPALHRRSRSSAASSGSGGGSGDEGRLLEPSITAAAAFTLLPSGSQVGYSVISL
DGLLGGRPKAEGFIQRQHGVLTTHVPQLGASAGGAERGLASPWFQLDSRGLGLHQAAAGRSSSGSGAASGGDSHPPFRYTATLVSGSGGATQQRGGQAPPAVQQQQQQQQQQQPEPQITYFFCSFD
GSCSVRSVANGFRCPLASCRGLRCHSYAALQQHIGAMHGYHASYFSGLLPDGGLEVYLRCKPAWLTPKLEFLPRQLGRVEEHPLLDLLSMSHNRYPFYFHCPRWQRRRRLGPWGAAPRDAREEELER
QAEANIAVRLGYAPPQPSSSAPAASSMYSVETLSDSGTGTELTAAEEAGPTAASGQRAAKPAGGRGRGRAGGGASGRGRGRAAAAVAGHHPASGLALLNGEGRPKFYHSRTCVAMTKEELFAEAGQA
GGDASDSEGDLSEWQASWWCLFVCRDCRERLAEHKGLCPEERAFMFEWNLFLHRRPLHADGDMPAALAAFAAGHREELGRDGPFRRCMVAHLLNLWRFRLLTPSQLHQLTRELPRVAPL

CSI2_123000000046

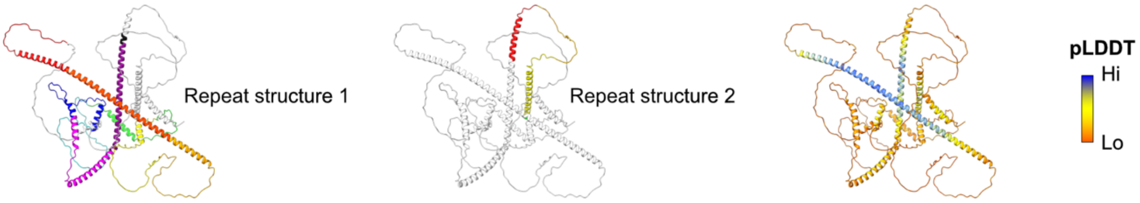


MPGTGVWDSCQQFLEAVERQQQRASSPCPQPSCPQQPPLSPQHPPQQQPQQQLGEQRPPSRTSSQPQQWPLPMASPFAAPNVQQAAPEPSGSAGAAGGEAKMVSPFGAAYLQEQAFSTGSLPAPSAP
LPDAKTLPSAEGPLHQPTRSGGESPPPLAQLQPQPAQQQGGAAAGQQQQRRSPSPDQWAAQQAQLEQQEAQEQEQQLEQHRAQAHQAALLAQQQRQKQAVAMGRLIEQQQQQQQSALLEKQLAAQQA
ALEQQRSEQQQWCTAAMRLIEQQQGQQAQQGQQGQRAQQMDWAAQRTQQAPPPPACPGCRQPVLPAVACACDGPLSLRRAGTANRSVLMVEASSLSRAGSSPGPSRAPNHISQASIAGAKVEADEDQ
QMQEVLSIARELGVFPTCPADLEAQPSWVPPSLGAPPPPPPLPRADQRRSEESAQSQLQLLQQQLLQPRSHAPGLTRGHSHPPPASTQQQQQQLLQPAPQLRREEWSAGGGAAPPALLAQPGMQQQP
AGRQLSADQLFELAHLLQRAKAAGIQLPQLASWPTQAPGPQAQPPVQPQLPPQQLHAAAQQQQLHGLVHQQQQQLAQPQPQPRRAETATFEHRSAMLAGLRRQISADAIHAHHVQQAQHAHRVQQAP
QVQQAQQAAMHRQQLMLLEAAQAEQRKQLVQAQHAELLQLEQARQARQARQAQQAQQAQHTWLAADARLAAPPAPISQHTLQFAGPQQQRSAYAAPPAPLQRLPTAQQQLQPPPLQQGAASLYAQQQ
ALLAQQRAEQQRWADAALSLLSQATR

CSI2_123000000713

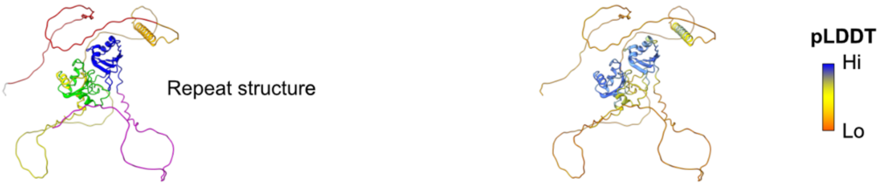


MSLFAGLFAAPKQADSPGGATLFSSSSKYAAQPLAAPAAAAAAAPEPGGGKKKKRKQEAAEAPAAPAAAAAEPSSKKARKAPQQRRQQGQEERQQQADKAAAAAAAAAAPAEGEKKANKKRKQQEGA
QQEQQGAAAPAAAAPAAAADGRGAKASSKPGKRDQASKPKPAAPAAEAAAEEGRSKAPGKAGGKATEPAAASKGKGGKAEAPADERLPRTVFVGNLPATVKTKKIARLFAAAGCGAVESVRLRSLPL
KQEPESKLPRKNAIAAGKIDSTHSAHAYVVFEEEAGVDKALALNMTEFEGHHIRVDRAAPASAKGAVAFDHMRTLFVGNLPHDCEDEELIRHFGAQAVEAVRIVRDGKTGAGKGVAFVLFRSQEASK
AALRARGSGELRGRKLRLTVAKKQAAAGGKPAAWQQGKDSGGKGGGKSGGKSGKPAAAQRKAGGKRPAVAARKAAAKMAAKGGVQKKRAA

CSI2_123000000775

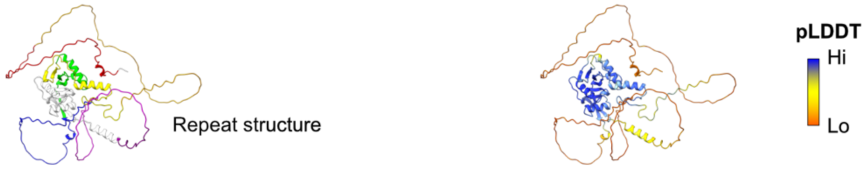


MQPPPGSIRAFFKPASKLAGAKRASGGSAAAAPAAEAPSDAIAAASEPAAAAADQLPPPPAVFARKRRRPSGGGSAGAAAEPTPAAAAAAAAGGDGAPAAAAGPSSAPATAAAAVEGGAAAGSRTRQ
LTQMYLDVGQRHFHAYRCPACGMLYARGTDADERLHAAFHASATQGPRYQGWAGERVVQLDGSKGRVLLFGSADQGRPRKQLADLCSFLEEQMGLSKGWLLQAPLTVLLYVSAARRVQGFVAAEPIK
EAYCARPSLPPAAAQGMLSCGSAVQPASGSGAAHAADAAATVAPFEGHPTAAAAAAQAHVATSGRSLLSVTLSTGAAAAECSSSAAAEGAGGGAPAAAAPAAAEAGDGEVAAAAGCTPGLPTSLQQQ
QQAQPEAQQAQQQLQQTQQPPVLPPLELDRSRRVRAVCGVRLMWVSLEARRRGLASRLLDCCRSHFMPGYVLSRHELAFSAPTDDGRAFIEQYAGSRKFLVYE

CSI2_123000001971

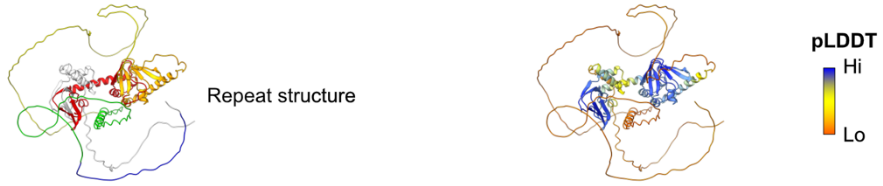


MGCGFSKGGGLRRGAVAPAPEEPAQAVGAQAGPPGGPTKRRGSVSAEPSGVSRSLSQAAGRQSVANQKALGTRERIRAIVKSTLLFKGLDEEQQGRMIDAMEERKVAAGEEIIKQGEEGDNFYVVES
GRFRATKSGAEVAAYDGAGSFGELALMYNCPRAATVTALSDGVLWALDRATFRALVVGSMAERAQRYEASLRRIPILQHLSQEQLAAIADCLEQERFEPGEYIVWEGAPLDDGAKFYMIESGRVDCF
RTFEGQKRAVKGLRAGDVFGEVALLTKAPRQADCVAATPCKCLTLSRAAFERLMGPVEQVLAQQIAQYQASNAQVTAGIDTTTASSSLPSLREDAAAAGNRGSEGAADDSKGDAGGSSQPPTSAGAG
KPPARRGSSPRSPAAGGGGSPAAGGSAAALLPGSPRPSMLSEGSAGSGSFSRKRSSPRATSNSLGARSARSTTNSDSAPAGEGDCPLGAAATGSGSGSGSGSGGAAELAVAMEEAAALSEQEPVCHL
PPIAEEPAEGATPRTSNASTQGSLQGQPLASSKERRSTAGGSTKGAAGGLSSGRRTSSGSGGKGKGPGSRPGAVKPAGSKPKAGSSKKK

**Extended Data Fig. 1 | *In silico* characterization of FLIPPer candidates.** Top ranked AlphaFold 2 structural predictions of the 13 candidate sequences identified by FLIPPer, and their accompanying full-length protein sequences below. Models on the left are colored according to the repeat sequences identified by XSTREAM, consistent with the coloring in the protein sequence. Gray sequence was not represented in the repeats. Models on the right are colored according to pLDDT score.

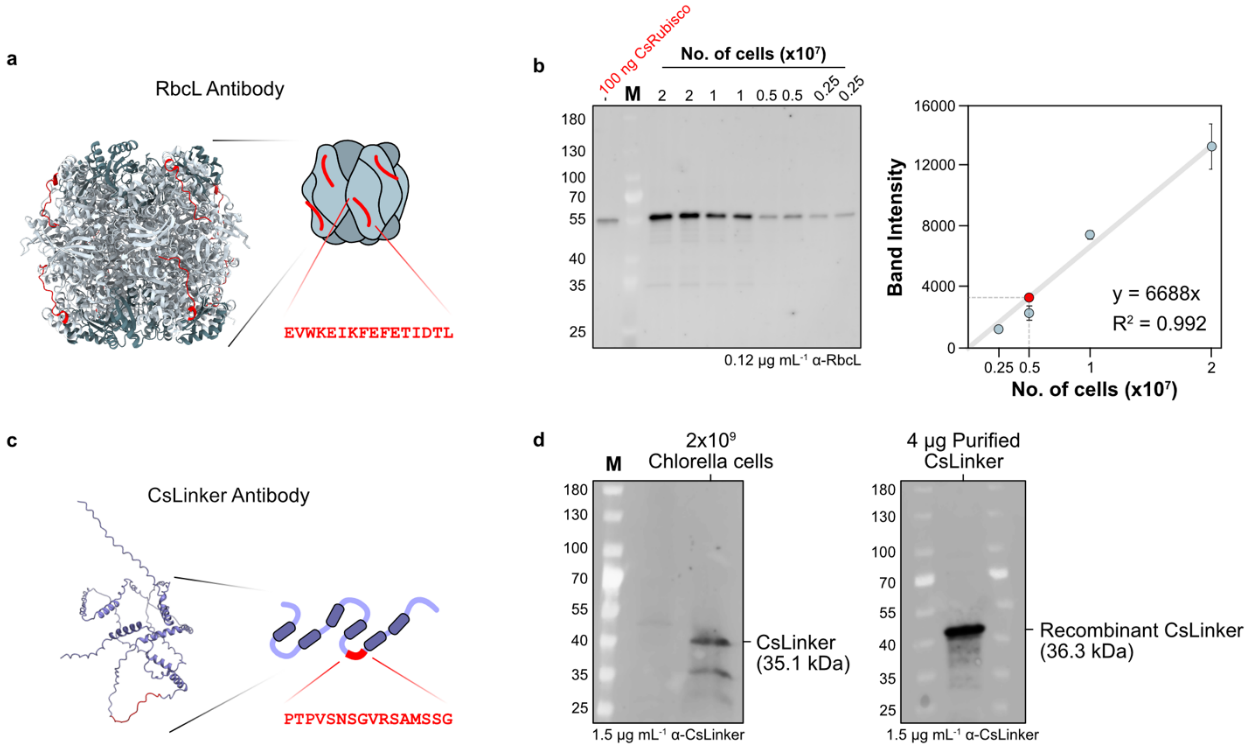


**Extended Data Fig. 2 | Antibodies raised to Rubisco large subunit and CsLinker. a**, Structure of *Chlamydomonas* Rubisco (PDB: 1EJ7[40]) with the region of the RbcL that the antibody was raised to colored red (left). Schematic representation of the location and sequence the peptide was raised to (right). **b**, Quantification of CsRubisco by western blot from lysate compared to purified CsRubisco. Band intensity was quantified and used to determine the amount of CsRubisco per cell. **c**, AlphaFold 2 structural prediction of CsLinker with the antibody peptide colored red and shown schematically adjacent. **d**, Western blot validation of the CsLinker antibody against lysate (left) and recombinantly produced CsLinker (right). Recombinant CsLinker has a higher molecular weight due to the presence of scar amino acids introduced by TEV cleavage sites.

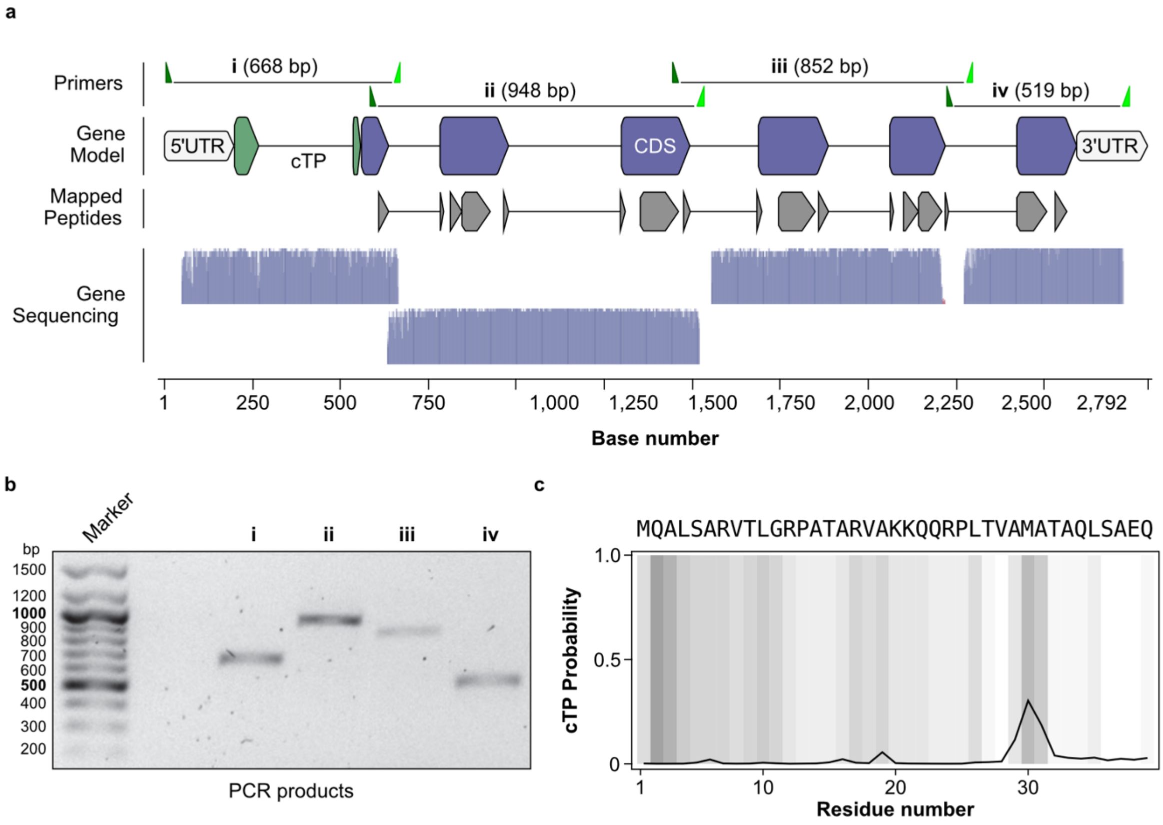


**Extended Data Fig. 3 | Validation of CsLinker annotation. a**, Schematic summary of the predicted gene model of CSI2_123000012064-RA, CsLinker. The primers used to amplify the sequences in b and their expected lengths (top) are shown relative to the gene model (below). In the gene model, exons are represented by blocks, connected by lines representing introns. The predicted chloroplast transit peptide (cTP) from c is shown in green. A selection of the mapped peptides from mass spectrometry experiments are shown below the gene model where the mapped peptide sequence is represented by blocks aligned to the relative position of the corresponding amino acids in the gene sequence. Peptides that span introns are represented with a line between adjacent blocks. Sequencing results, represented as quality chromatograms, mapped to the gene are shown at the bottom of the figure. **b**, DNA gel electrophoresis of PCR products corresponding to the mapped products in a. The same products were sequenced. **c**, TargetP 2.0 prediction of the chloroplast transit peptide location in the first 39 residues. The predicted cleavage site is after the Met30 residue.

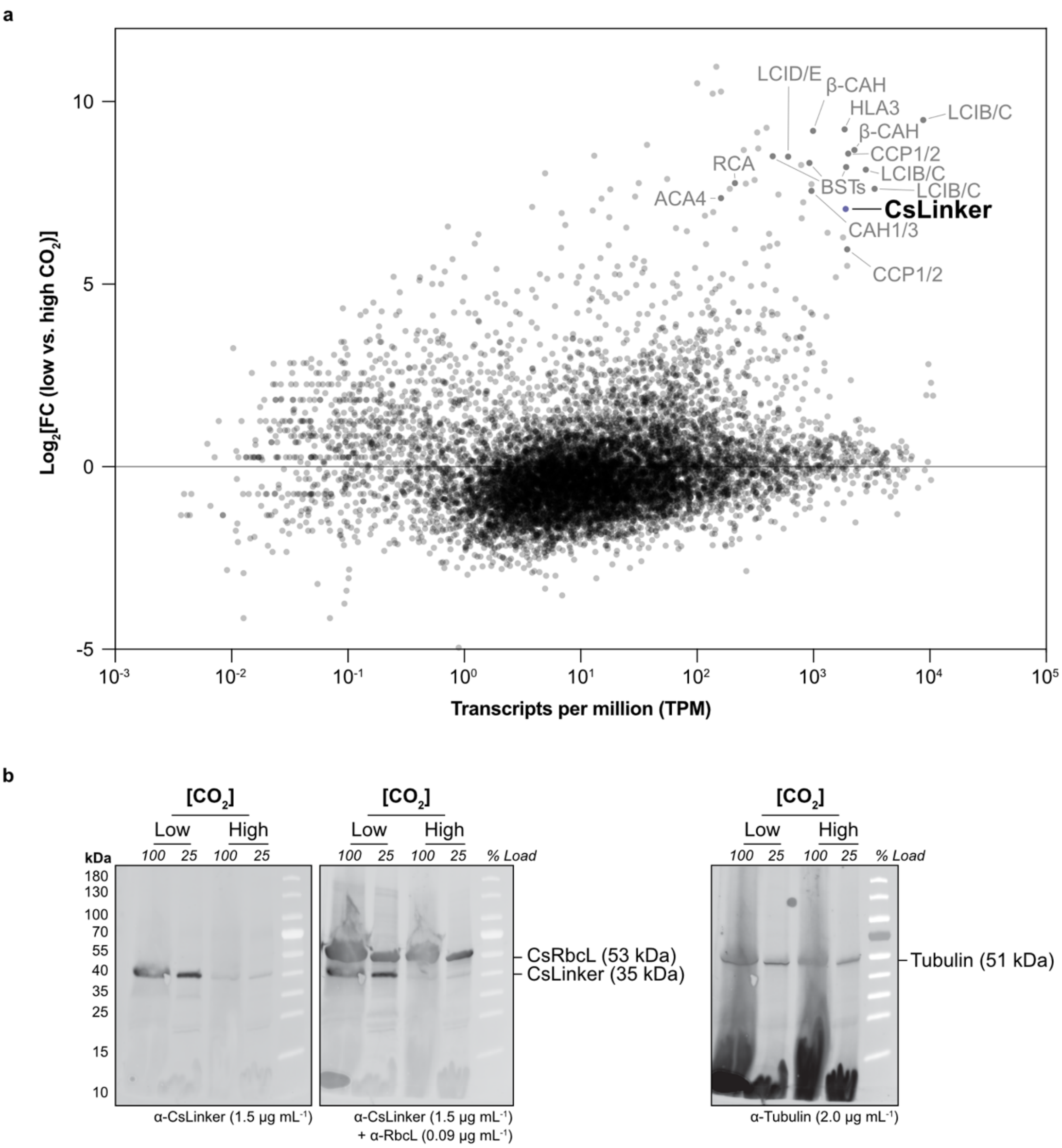


**Extended Data Fig. 4 | Low $CO_2$ inducibility of CsLinker. a**, Differential gene expression analysis completed using publicly available RNA-seq data (PRJNA343632) comparing transcript abundance at high $CO_2$ (5%) and 30 minutes after switching to 0.01% $CO_2$ culture conditions. Homologues of low $CO_2$ induced CCM genes in *Chlamydomonas* are indicated alongside CsLinker. **b**, Western blot analysis of CsLinker and CsRbcL protein abundance under low (0.04%) and high (3%) $CO_2$ conditions. Tubulin was used as a loading control.

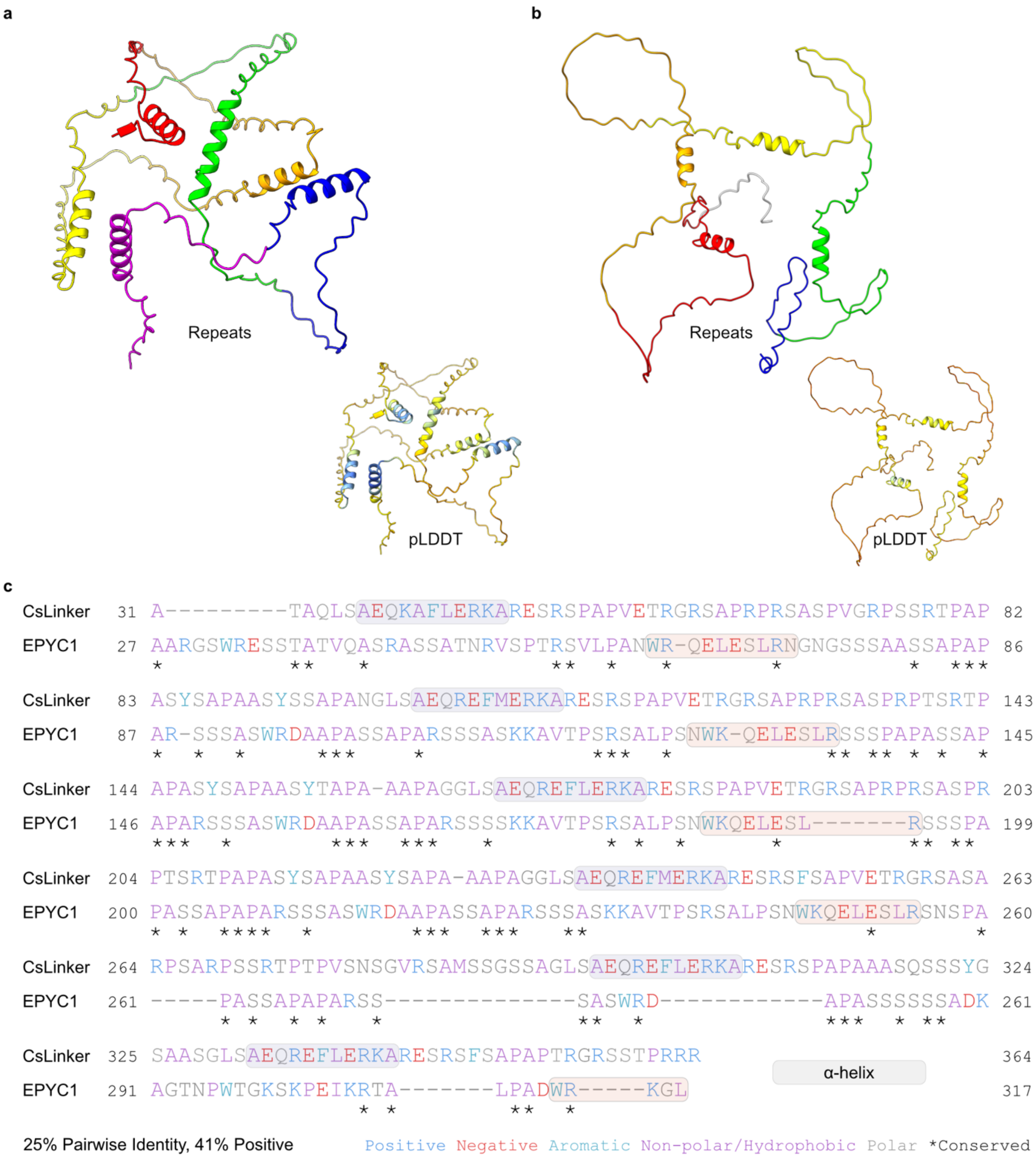


**Extended Data Fig. 5 | Comparison of CsLinker and EPYC1. a**, Top ranked AlphaFold 2 structural prediction of CsLinker displayed without the predicted chloroplast transit peptide (cTP) - colored by XSTREAM-identified repeat structure (top) and pLDDT (bottom). **b**, Top ranked AlphaFold 2 structural prediction of EPYC1 without cTP. **c**, Alignment of CsLinker and EPYC1 sequences after removal of the cTPs. Residues are colored by property; conserved residues are indicated (asterisks) and the predicted α-helices are shown. Alignment was completed with MAFFT v7.49[70] using a BLOSUM62 scoring matrix with a gap open penalty of 1.53 and an offset value of 0.123.

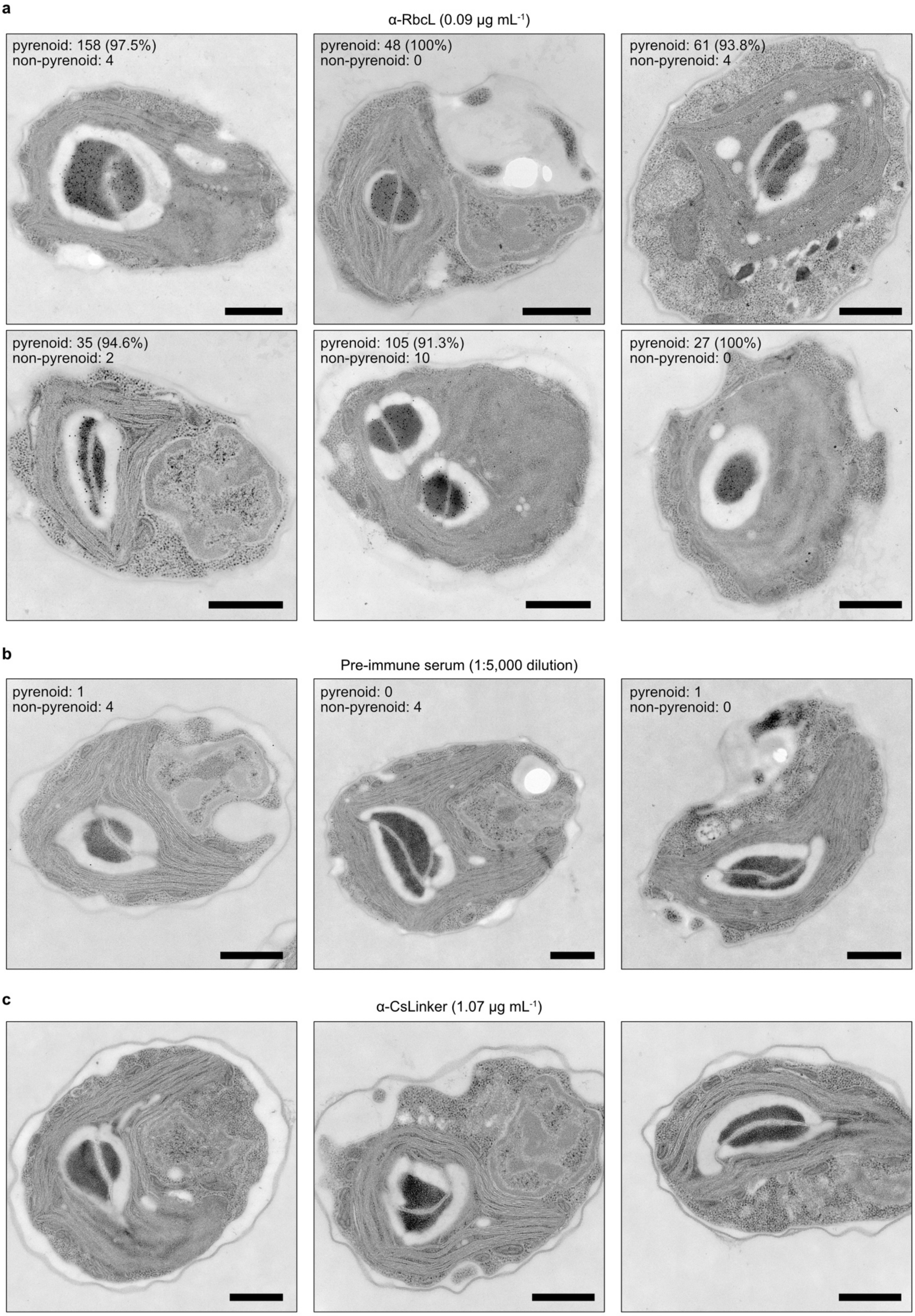
a
α-RbcL (0.09 µg mL-1)
pyrenoid: 158 (97.5%)
non-pyrenoid: 4
pyrenoid: 48 (100%)
non-pyrenoid: 0
pyrenoid: 61 (93.8%)
non-pyrenoid: 4
pyrenoid: 35 (94.6%)
non-pyrenoid: 2
pyrenoid: 105 (91.3%)
non-pyrenoid: 10
pyrenoid: 27 (100%)
non-pyrenoid: 0
b
Pre-immune serum (1:5,000 dilution)
pyrenoid: 1
non-pyrenoid: 4
pyrenoid: 0
non-pyrenoid: 4
pyrenoid: 1
non-pyrenoid: 0
c
α-CsLinker (1.07 µg mL-1)

**Extended Data Fig. 6 | Immunogold localization of CsRubisco in the pyrenoid. a,** Transmission electron micrographs of Chlorella cells that were immunogold labelled following primary incubation with RbcL antibody at 0.09 μg $mL^{-1}$. **b**, TEM images following incubation with a 1:5,000 dilution of the pre-immune serum from the same rabbit used to raise the RbcL antibody. **c**, TEM images following incubation with the CsLinker antibody at 1.07 μg $mL^{-1}$. Scale bar in all panels = 500 nm.

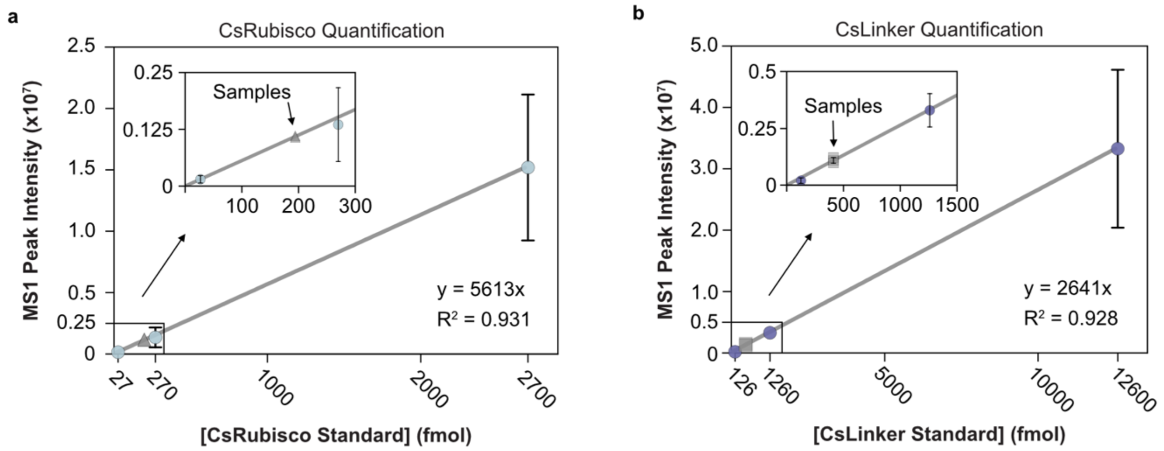


**Extended Data Fig. 7 | Absolute quantification of CsRubisco and CsLinker. a**, Standard curve obtained using purified CsRubisco (blue circles), used to quantify CsRubisco abundance *in vivo* (grey triangles). **b**, Standard curve used to quantify CsLinker abundance *in vivo* (grey squares. In both panels error bars represent S.D. and the zoomed inset shows the position of the *in vivo* samples.

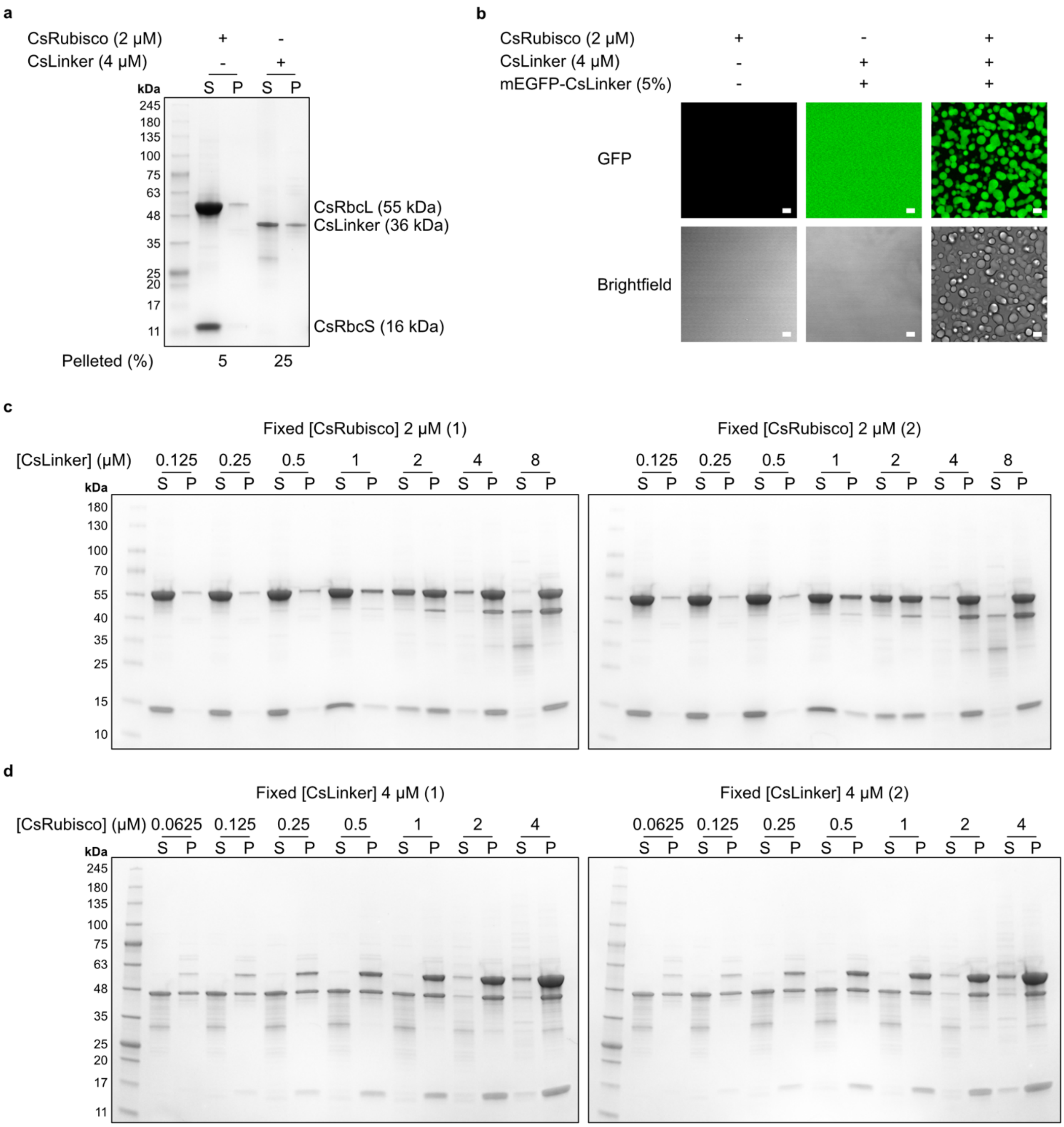


**Extended Data Fig. 8 | Characterisation of Chlorella pyrenoid reconstitution *in vivo*.** **a**, Control droplet sedimentation assays in which CsRubisco and CsLinker were incubated alone and analyzed by SDS-PAGE. **b**, Confocal fluorescence microscopy images of control droplet assays. Scale bar = 5 μm. **c**, SDS-PAGE analysis of droplet sedimentation assays with CsRubisco fixed at 2 μM and CsLinker titrated as indicated. **d**, Droplet sedimentation assays with CsLinker fixed at 4 μM and CsLinker titrated.

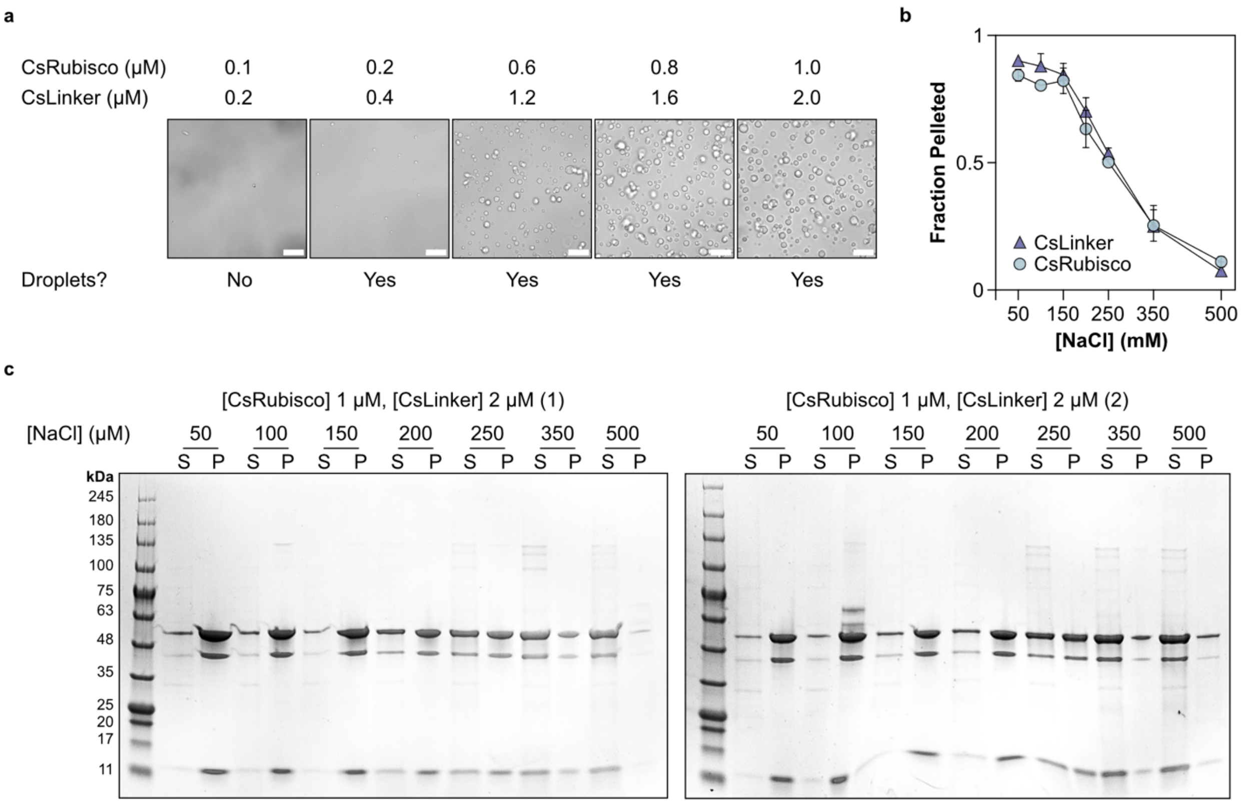


**Extended Data Fig. 9 | Indicators of droplet formation by LLPS. a**, Brightfield images of droplet assays performed at increasing global concentrations of CsLinker and CsRubisco to determine a critical concentration for LLPS. Scale bar = 5 µm. **b**, Quantification of CsLinker and CsRubisco pelleting in salt-dependency droplet sedimentation assays. **c**, SDS-PAGE analysis of salt-dependency droplet sedimentation assays.

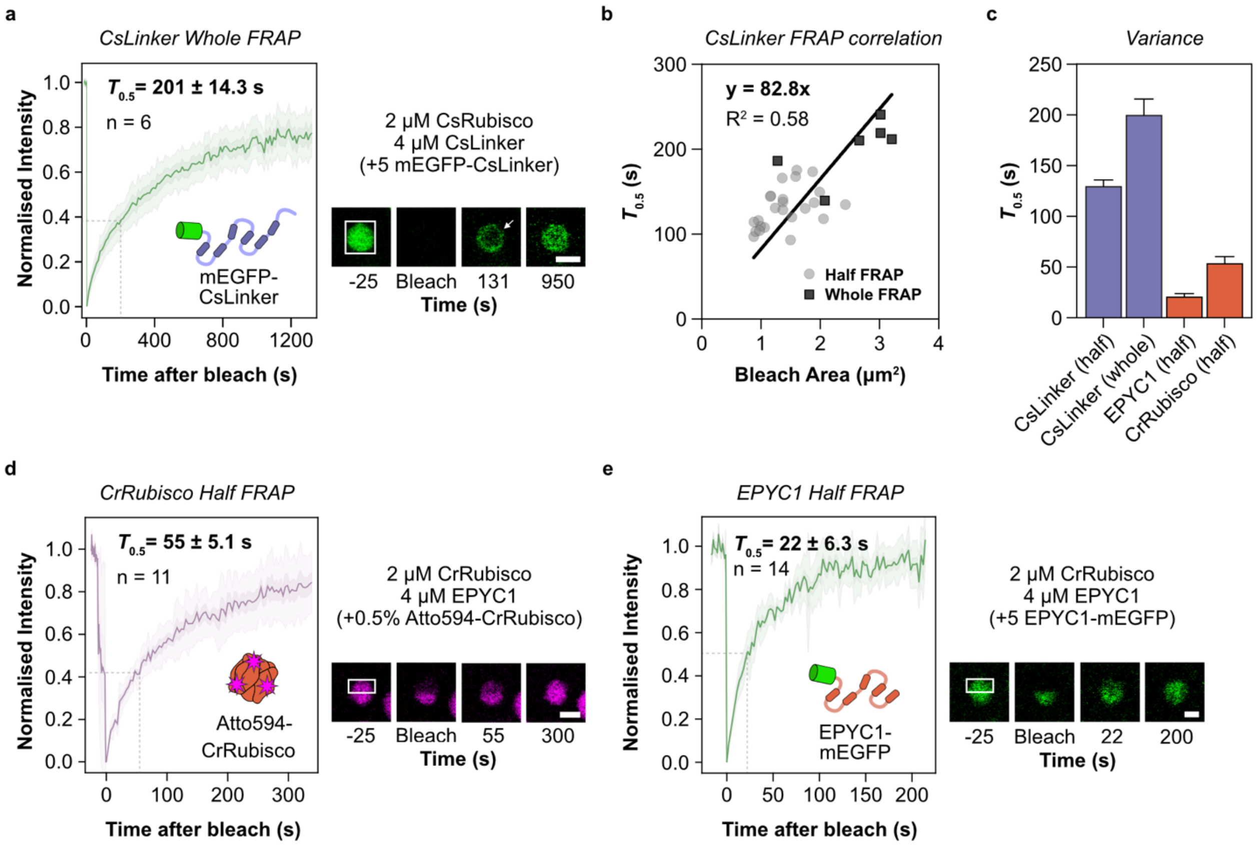


**Extended Data Fig. 10 | FRAP analysis of Chlorella and *Chlamydomonas in vitro* reconstitutions. a**, Average FRAP recovery curve from whole FRAP experiments completed according to reference images adjacent where the bleach region is indicated by the box and the scale bar = 1 µm. The arrow highlights recovery of the signal from the periphery of the droplet, indicating external exchange. The standard error of the $T_{0.5}$ is indicated in the plot and the dashed lines indicate the $T_{0.5}$ on the plot. **b**, Correlation of $T_{0.5}$ with the area of the bleached region in whole and half FRAP experiments of CsLinker, explaining the longer $T_{0.5}$ in whole FRAP experiments. **c**, Variance of $T_{0.5}$ values derived from individual fits of recovery curves. Errors bars represent S.D. **d**, Average half FRAP recovery curve of Atto594-CrRubisco in the *Chlamydomonas* reconstitution. **e**, Average half FRAP recovery curve of EPYC1-mEGFP in the *Chlamydomonas* reconstitution.

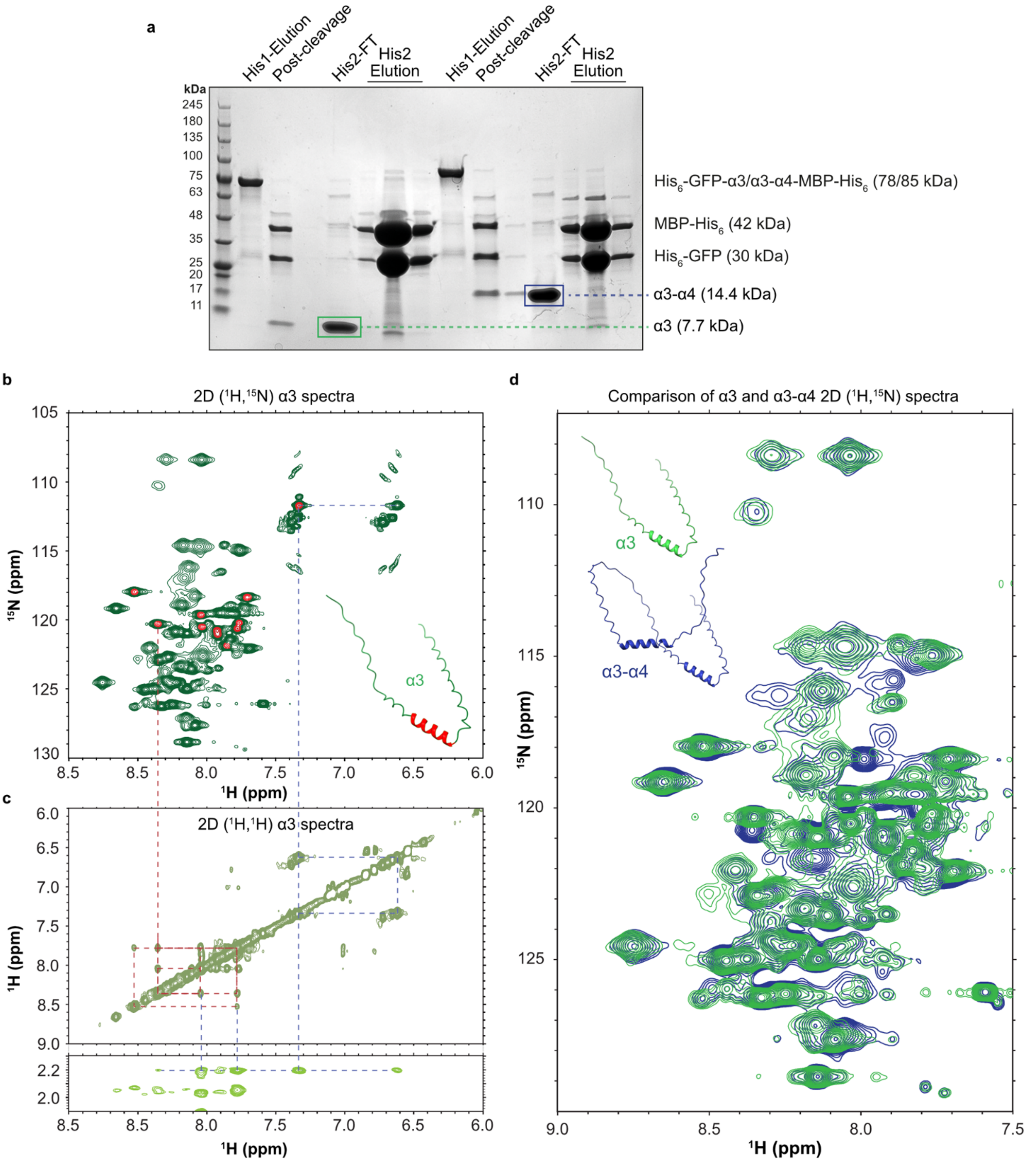


**Extended Data Fig. 11 | 2D NMR spectroscopy confirms the presence of pre-formed α-helices in CsLinker. a**, Purification of the α3 and α3-α4 CsLinker fragments. SDS-PAGE analysis of the first nickel affinity product (His1), the samples following overnight cleavage with TEV protease, the flow through from the second nickel affinity purification (His2) and the elution. Annotations show the expected migration distances of the various species. **b**, 2D ($^1$H,$^{15}$N) SOFAST HMQC spectrum of the α3 fragment in solution. The low $^1$H spectra dispersion of the amide-proton resonances (~1.5 ppm) is typical of proteins with significant intrinsic disorder. Approximately 60 of the 62 expected cross peaks are visible in the 2D spectrum (the 73 residue α3 construct has 11 prolines). **c**, Regions of a 2D ($^1$H, $^1$H) NOESY spectrum of the same fragment in a, showing approximately 10 NOE cross-peaks in the amide-proton region (red dashed lines), which is consistent with the formation of a stable α-helix, as per the predicted structure. Consistent with the primary sequence of α3, the α-helical region contains at least one glutamine residue, as evidenced by the observation of NOEs between backbone amide and side chain amide protons and aliphatic protons (blue dashed lines). **d**, Overlay of the 2D ($^1$H, $^{15}$N) SOFAST HMQC spectra of α3 (green) and α3-α4 (blue) showing significant overlap in the position of the cross-peaks. The small number of non-overlapping cross peaks observed is consistent with the small sequence differences between the α3 and α3-α4 repeat regions.

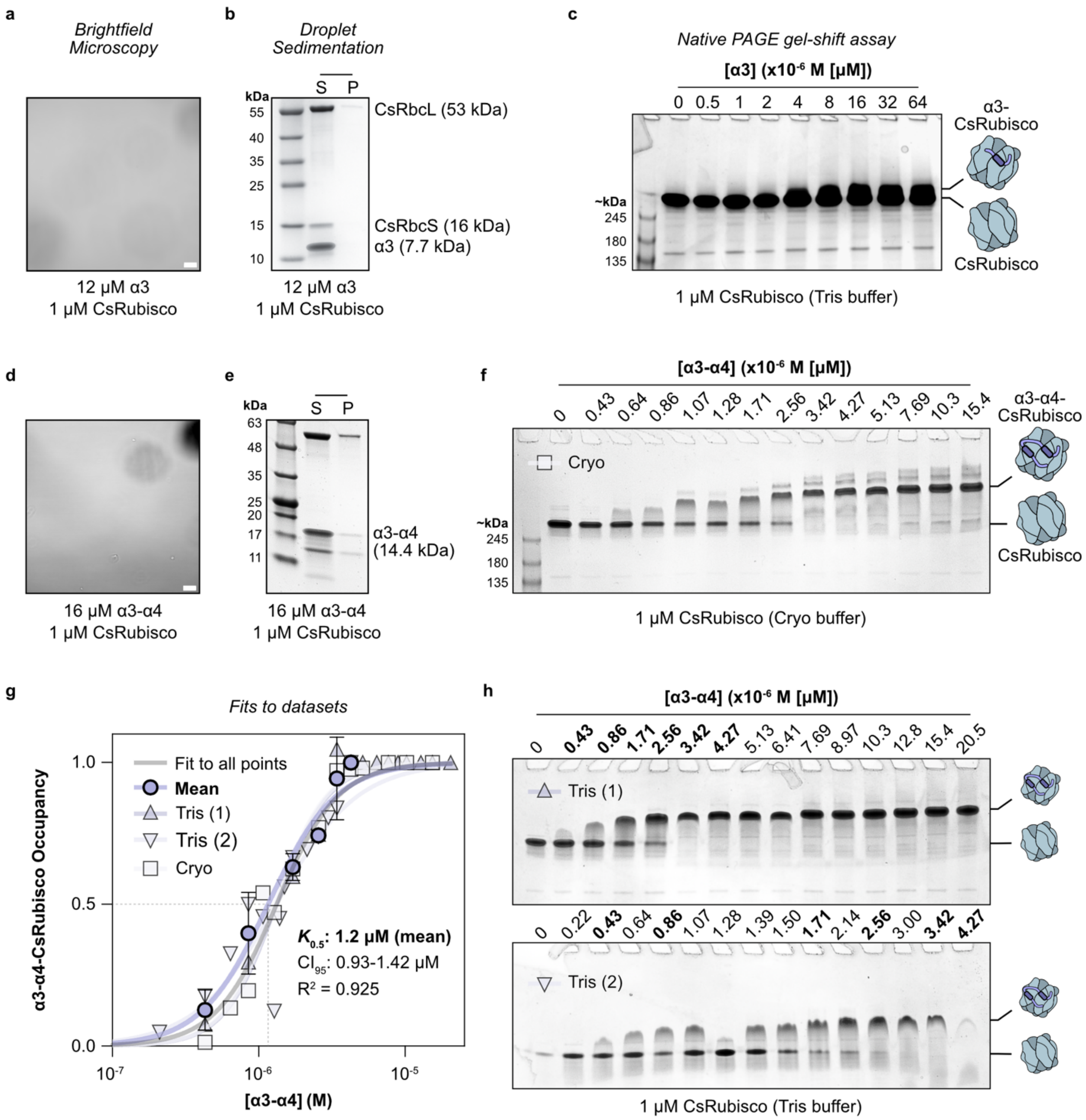


**Extended Data Fig. 12 | α3 and α3-α4 CsLinker fragments do not phase separate CsRubisco. a**, Brightfield microscopy image of α3-CsRubisco mix, scale bar 5 µm. **b**, Droplets sedimentation assay of α3-CsRubisco mix. **c**, Native PAGE gel-shift assay of CsRubisco incubated with increasing concentrations of α3 CsLinker fragment in Tris buffer (50 mM Tris-HCl, pH 8.0, 50 mM NaCl). **d**, Brightfield microscopy of α3-α4-CsRubisco mix. **e**, Droplet sedimentation of α3-α4-CsRubisco mix**. f**, Native PAGE gel-shift assay of CsRubisco incubated with increasing concentrations of α3-α4 in cryo buffer (200 mM sorbitol, 50 mM HEPES, 50 mM KOAc, 2 mM $Mg(OAc)_2 \cdot 4H_2O$ and 1 mM $CaCl_2$ at pH 6.8). **g**, Quantification of α3-α4-CsRubisco complex occupancy from native PAGE gel shift assays in f and h, derived from the inverse occupancy of the unbound CsRubisco state. Datasets were fitted separately as well as in a fit to all points using a Hill coefficient model ($Y = x^h / (K_{0.5}{}^h + x^h)$), where h is the Hill coefficient. The 'mean' dataset was derived from the bolded replicate values in the Tris (1) and Tris (2) experiments in h. A fit to this dataset was used to derive the $K_{0.5}$ of the α3-α4-CsRubisco complex as presented in Fig. 2a. **h**, Native PAGE gel-shift assays completed with CsRubisco and : α3-α4 in Tris buffer. Bolded concentrations were used to derive the 'Mean' dataset in g.

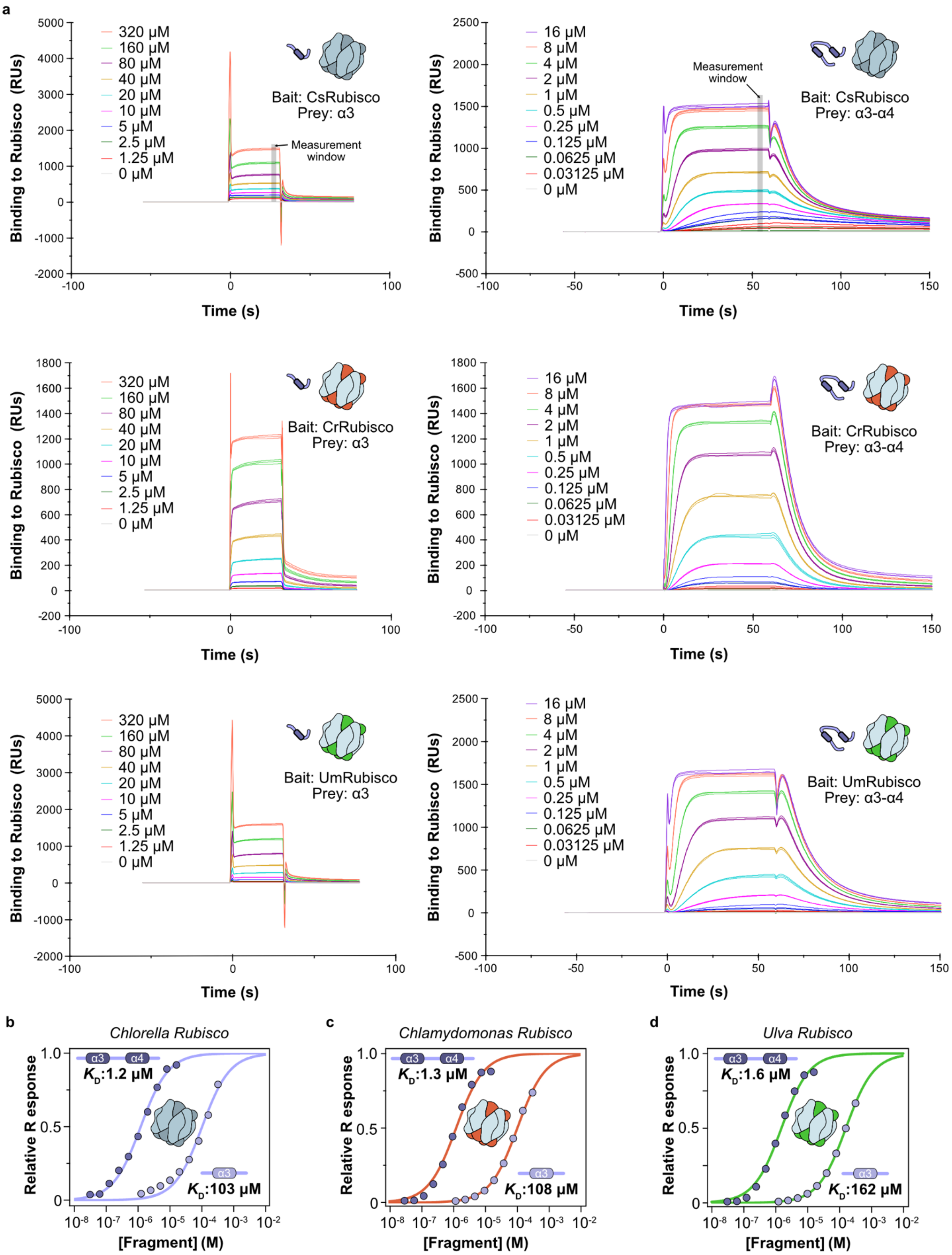
a
Bait: CsRubisco
Prey: α3
Measurement window
Bait: CsRubisco
Prey: α3-α4
Bait: CrRubisco
Prey: α3
Bait: CrRubisco
Prey: α3-α4
Bait: UmRubisco
Prey: α3
Bait: UmRubisco
Prey: α3-α4
320 μM
160 μM
80 μM
40 μM
20 μM
10 μM
5 μM
2.5 μM
1.25 μM
0 μM
16 μM
8 μM
4 μM
2 μM
1 μM
0.5 μM
0.25 μM
0.125 μM
0.0625 μM
0.03125 μM
0 μM
Binding to Rubisco (RUs)
Time (s)
b
Chlorella Rubisco
$K_D$:1.2 μM
$K_D$:103 μM
c
Chlamydomonas Rubisco
$K_D$:1.3 μM
$K_D$:108 μM
d
Ulva Rubisco
$K_D$:1.6 μM
$K_D$:162 μM
α3
α4
Relative R esponse
[Fragment] (M)

**Extended Data Fig. 13 | Sensorgrams from SPR experiments. a**, Raw sensorgrams from surface plasmon resonance experiments with the α3 and α3-α4 fragments at the indicated concentrations against Chlorella (Cs), *Chlamydomonas* (Cr) and *Ulva* (Um) Rubiscos. In the top panels, the window in which the SPR response was measured is indicated. **b**, Relative SPR response curves for α3 and α3-α4 fragments against CsRubisco as presented in Fig. 3b. SPR response is normalized to the fitted $B_{max}$ value obtained from fit of the raw data. n=3, error bars = S.D. **c**, SPR response curve against *Chlamydomonas* Rubisco. **d**, SPR response curve against *Ulva* Rubisco.

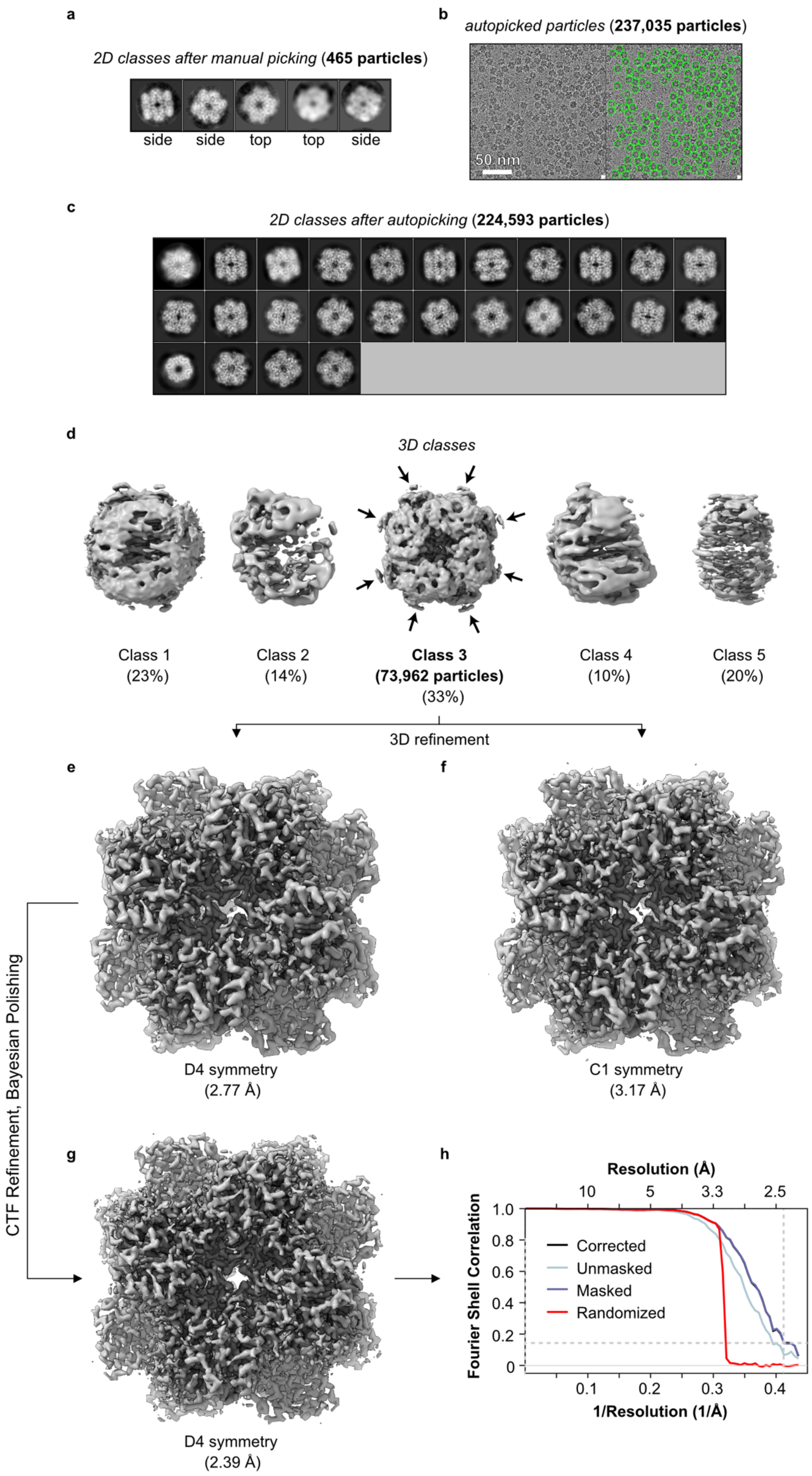
a
2D classes after manual picking (465 particles)
side
side
top
top
side
b
autopicked particles (237,035 particles)
50 nm
c
2D classes after autopicking (224,593 particles)
d
3D classes
Class 1
(23%)
Class 2
(14%)
Class 3
(73,962 particles)
(33%)
Class 4
(10%)
Class 5
(20%)
3D refinement
e
D4 symmetry
(2.77 Å)
f
C1 symmetry
(3.17 Å)
CTF Refinement, Bayesian Polishing
g
D4 symmetry
(2.39 Å)
h
Resolution (Å)
10
5
3.3
2.5
Fourier Shell Correlation
1.0
0.8
0.6
0.4
0.2
0
Corrected
Unmasked
Masked
Randomized
0.1
0.2
0.3
0.4
1/Resolution (1/Å)

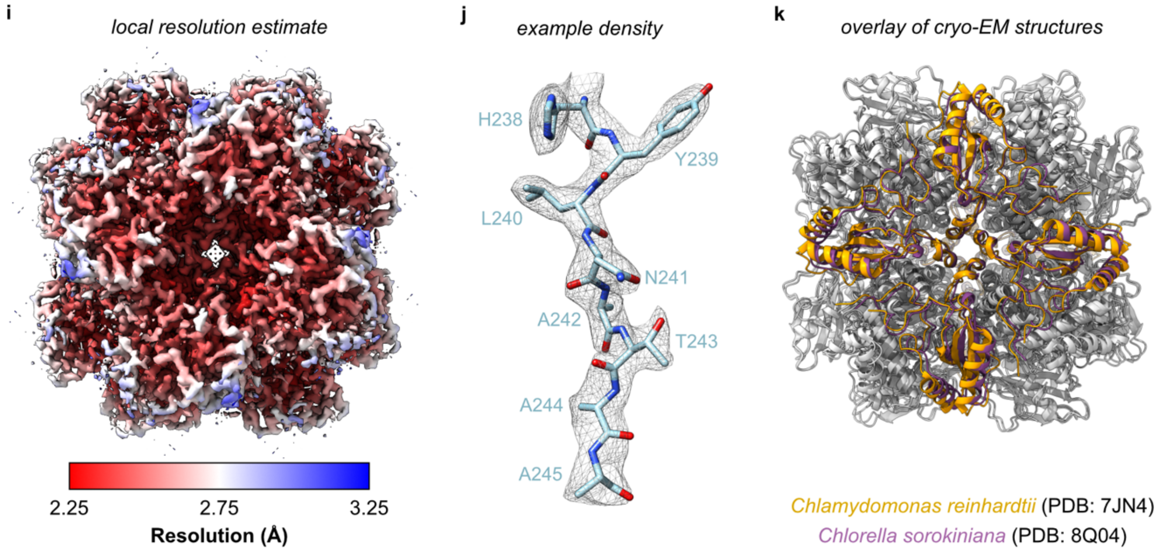


**Extended Data Fig. 14 | Cryo-EM data processing of the D4 map (PDB: 8Q04). a**, 2D classes after manual picking of 465 particles, used for autopicking from all grids. **b**, Representative micrograph shown without (left) and with (right) autopicked particles (green circles). Autopicking resulted in 237,035 particles that were used in subsequent 2D classification. **c**, Selected 2D classes after classification, resulting in 224,593 particles that were used for 3D classification. **d**, Top view of the five 3D classes following classification, shown at the same contour level (0.01). Class 3 was used for subsequent refinements to create the final D4 map. Arrows indicate the regions of additional low resolution density at the equator of Rubisco. **e**, Post-processed map following refinement of class 3 with D4 symmetry imposed during refinement. **f**, Post-processed map following refinement with C1 symmetry. Maps in e and f are shown at a contour level of 0.032. **g**, Post-processed map following CTF refinement and Bayesian polishing with D4 symmetry imposed during refinement. The map is displayed at a contour level of 0.0553. **h**, Fourier shell correlation (FSC) curve showing the resolution estimate for the D4 refined map with FSC cut-off of 0.143 (dashed lines). **i**, Phenix local resolution estimate for the D4 CsRubisco map. **j**, Example density of residues 238-245 of the CsRbcL with the corresponding model coordinates, carved with a radius of 2 at a contour level of 0.0415. **k**, Overlay of cryo-EM structures of Rubisco from Chlorella solved in this study and from *Chlamydomonas reinhardtii* solved in a previous study[32].

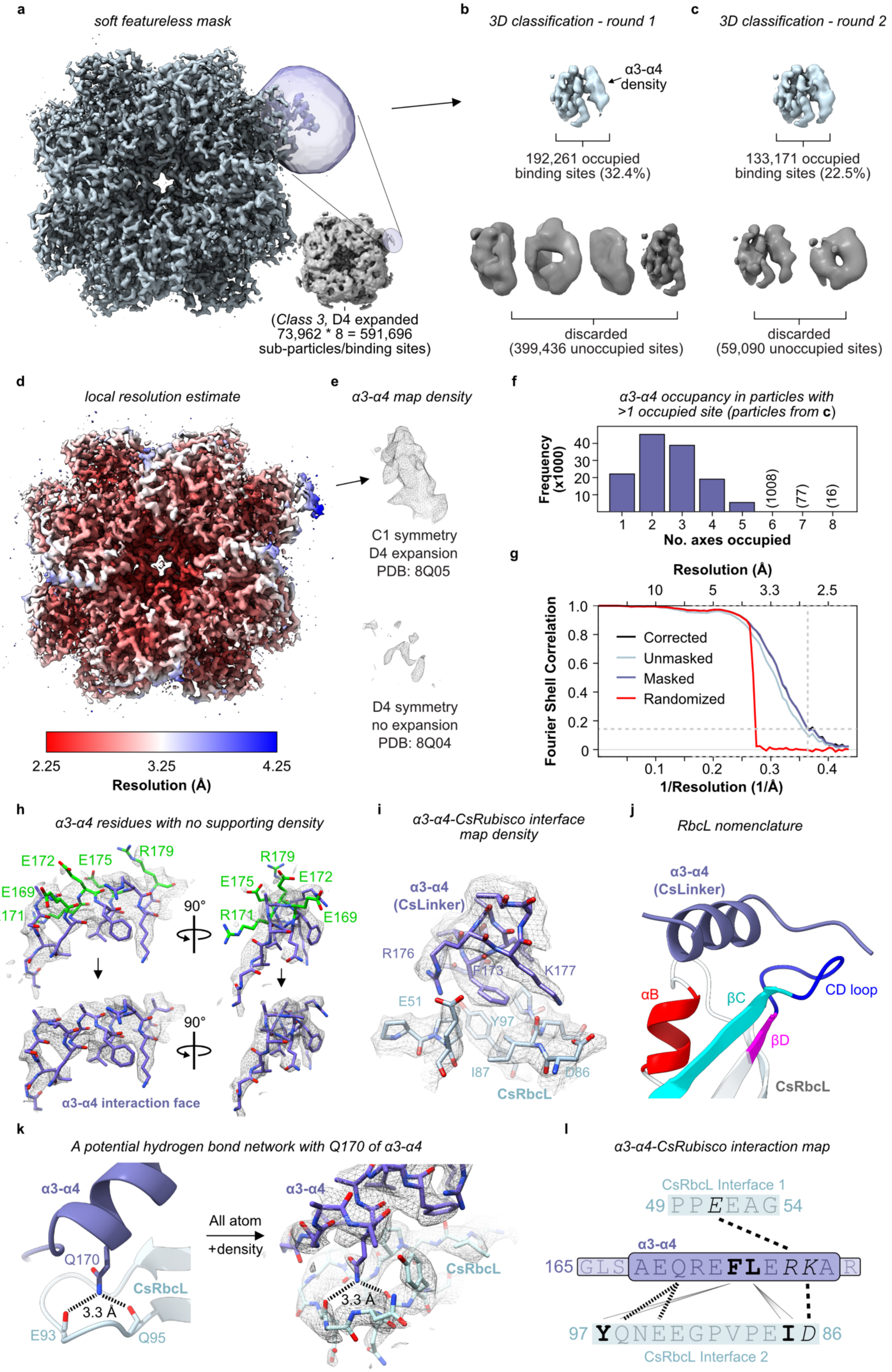

a
soft featureless mask
(Class 3, D4 expanded
73,962 * 8 = 591,696
sub-particles/binding sites)
b
3D classification - round 1
α3-α4
density
192,261 occupied
binding sites (32.4%)
discarded
(399,436 unoccupied sites)
c
3D classification - round 2
133,171 occupied
binding sites (22.5%)
discarded
(59,090 unoccupied sites)
d
local resolution estimate
2.25
3.25
4.25
Resolution (Å)
e
α3-α4 map density
C1 symmetry
D4 expansion
PDB: 8Q05
D4 symmetry
no expansion
PDB: 8Q04
f
α3-α4 occupancy in particles with
>1 occupied site (particles from c)
Frequency
(x1000)
(1008)
(77)
(16)
No. axes occupied
g
Resolution (Å)
Fourier Shell Correlation
Corrected
Unmasked
Masked
Randomized
1/Resolution (1/Å)
h
α3-α4 residues with no supporting density
E172
E175
R179
E169
R171
90°
α3-α4 interaction face
i
α3-α4-CsRubisco interface
map density
α3-α4
(CsLinker)
R176
F173
K177
E51
Y97
I87
D86
CsRbcL
j
RbcL nomenclature
α3-α4
(CsLinker)
αB
βC
CD loop
βD
CsRbcL
k
A potential hydrogen bond network with Q170 of α3-α4
α3-α4
Q170
CsRbcL
3.3 Å
E93
Q95
All atom
+density
l
α3-α4-CsRubisco interaction map
CsRbcL Interface 1
49 PPEEAG 54
α3-α4
165 GLS AEQREFLERKA R
97 YQNEEGPVPEID 86
CsRbcL Interface 2

**Extended Data Fig. 15 | Cryo-EM data processing of the α3-α4-CsRubisco map (PDB: 8Q05). a**, Sharpened (*B*-factor: 45) post-processed map of the C1 symmetry-refined α3-α4-CsRubisco complex using the D4 symmetry expanded particle dataset. The soft featureless mask used for the classification is shown in purple, over the region of additional density, into which the predicted helix of α3 is built. Shown at a contour level of 0.0496. The 3D class (class 3) from which the particle dataset was D4 symmetry expanded is shown inset, and the mask is schematically represented over a region of additional density. **b**, C1 symmetry 3D classification of sub-particles using the soft featureless mask. The selected class with α3-α4 density is shown in blue, with the discarded classes in grey below. At this resolution, the density shows clearly helical nature. **c**, Second round of 3D classification using the selected sub-particles from the first round. The selected sub-particles from this round were used for reconstruction of the α3-α4-CsRubisco map. **d**, Phenix local resolution estimate of the α3-α4-CsRubisco map, shown at a contour level of 0.0293. **e**, Map density of the α3-α4 region in the unsharpened, post-processed C1, symmetry expanded map shown at contour level 0.0293 (top), and the unsharpened, post-processed D4, non-symmetry expanded map, shown at a contour level of 0.0174. Both maps are carved with a radius of 2 around the modelled helical region. **f**, Histogram showing the distribution of α3-α4 occupancy in the sub-particles of the particles used for the reconstruction of the α3-α4-CsRubisco map. **g**, Fourier shell correlation (FSC) curve showing the resolution estimate for the α3-α4-CsRubisco map with FSC cut-off of 0.143 (dashed lines). **h**, Model of α3-α4 in the density displayed with the side chains of residues with no density support displayed in green (top) and removed in the final coordinates (bottom). **i**, Coordinates at the α3-α4-CsRubisco interface displayed in the map density at a contour level of 0.0304 and carved with a radius of 2. **j**, Nomenclature of RbcL regions at the α3-α4-CsRubisco interface. **k**, A potential hydrogen bond network between Gln170 of α3-α4 and Glu93 and Gln95 of the CsRbcL CD loop shown with and without map density shown at a contour level of 0.0396. **l**, α3-α4-CsRubisco interaction map.

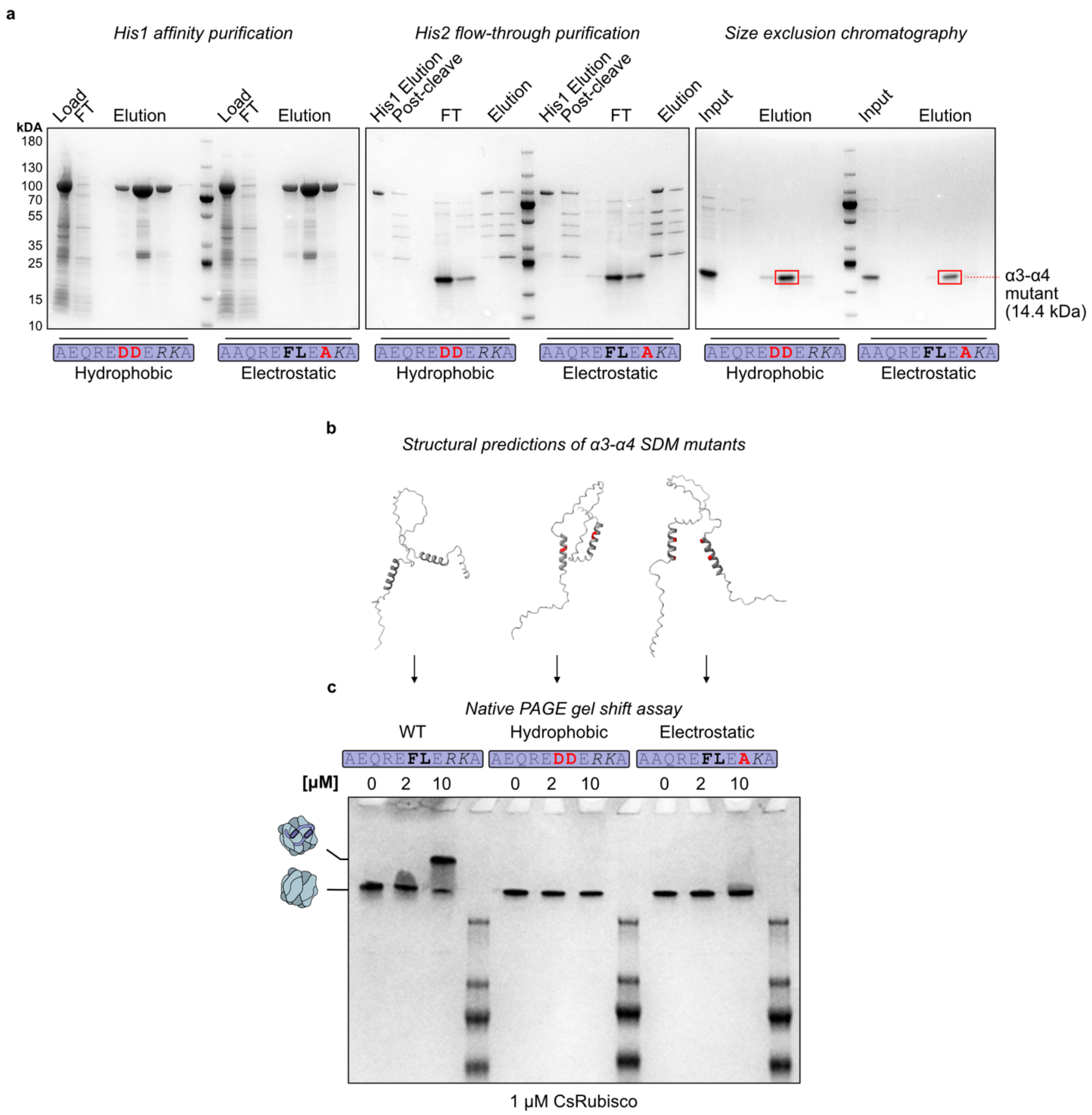


**Extended Data Fig. 16 | Purification of α3-α4 SDM variants and native PAGE analysis. a**, Purification strategy for the hydrophobic and electrostatic α3-α4 mutants produced by site-directed mutagenesis. **b**, AlphaFold 2 structural predictions of the WT and mutant α3-α4 fragments, with the mutated residues shown in red, according to the schematics in a. **c**, Native PAGE gel shift assay completed with the mutant fragments compared to WT.

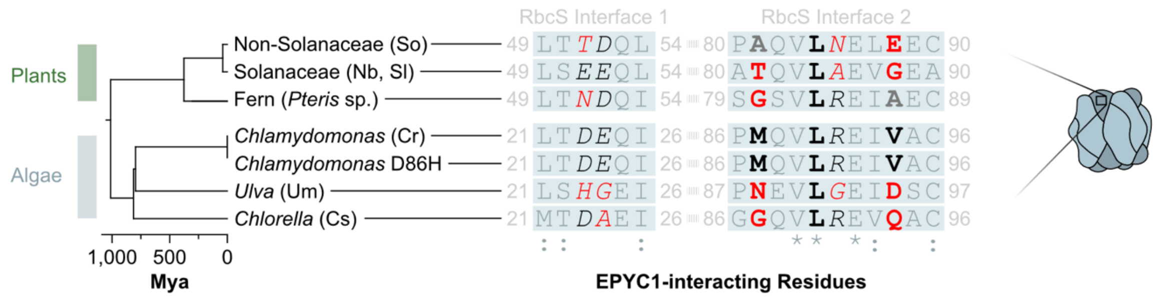


**Extended Data Fig. 17 | Comparison of EPYC1-interacting interfaces in green lineage RbcS sequences.** Residues shown to interact with EPYC1 in a previous study[32] are colored black, with hydrophobic residues in bold and electrostatic residues in italics. Residues that do not share similar properties in other species are shown in red and stylized according to the corresponding residues in the *Chlamydomonas* sequence. No sequence was available for the *Adiantum* RbcS, so a *Pteris* fern RbcS sequence is presented instead.

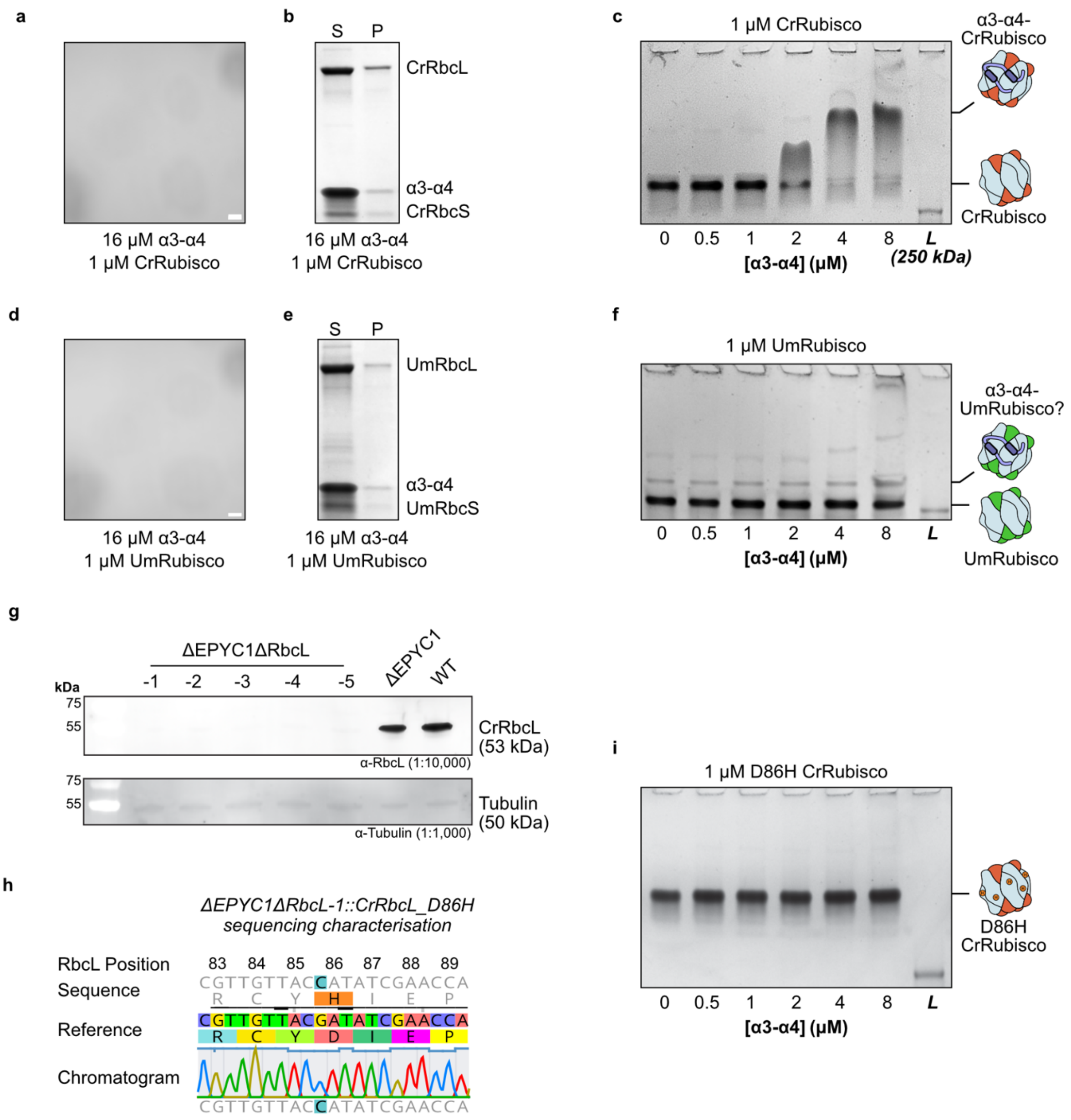


**Extended Data Fig. 18 | Cross-reactivity of α3-α4. a**, Brightfield microscopy of α3-α4-CrRubisco solution at the indicated concentrations, scale bar = 5 µm. **b**, Droplet sedimentation assay of the same solution as in a. **c**, Native PAGE gel shift assay of 1 µm CrRubisco incubated with increasing concentrations of α3-α4 demonstrating shift to a higher order α3-α4-CrRubsico complex. **d**, Microscopy of α3-α4-UmRubisco solution. **e**, Droplet sedimentation assay of the same solution as in d. **f**, Native PAGE gel shift assay of α3-α4-UmRubisco solutions demonstrating possible complex formation at higher concentrations of α3-α4. **g**, Western blot analysis of RbcL protein presence in ΔEPYC1ΔRbcL knockout lines compared to the parental ΔEPYC1 and WT lines. Tubulin is used as a loading control. **h**, Sequencing of the ΔEPYC1ΔRbcL::CrRbcL_D86H strain that was complemented with a point mutated version of CrRbcL to introduce the D86H substitution. **i**, Native PAGE gel shift assay with α3-α4 and D86H mutated CrRubisco indicating lack of complex formation.

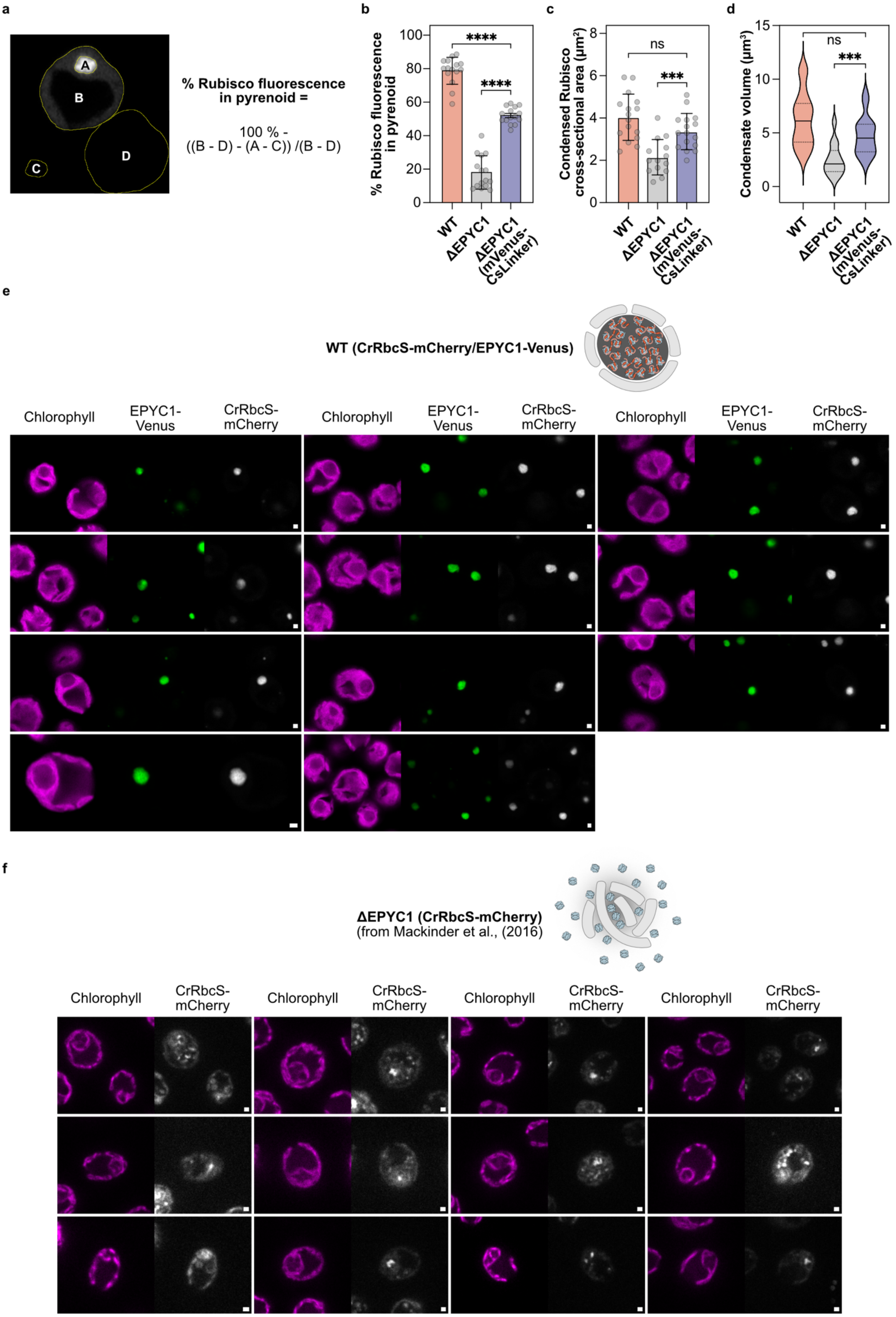
a
A
B
C
D
% Rubisco fluorescence in pyrenoid =
100 % - ((B - D) - (A - C)) /(B - D)
b
% Rubisco fluorescence in pyrenoid
0
20
40
60
80
100
****
****
WT
ΔEPYC1
ΔEPYC1 (mVenus-CsLinker)
c
Condensed Rubisco cross-sectional area (μm²)
0
2
4
6
8
ns
***
WT
ΔEPYC1
ΔEPYC1 (mVenus-CsLinker)
d
Condensate volume (μm³)
0
5
10
15
ns
***
WT
ΔEPYC1
ΔEPYC1 (mVenus-CsLinker)
e
WT (CrRbcS-mCherry/EPYC1-Venus)
Chlorophyll
EPYC1-Venus
CrRbcS-mCherry
f
ΔEPYC1 (CrRbcS-mCherry)
(from Mackinder et al., (2016)
Chlorophyll
CrRbcS-mCherry

g

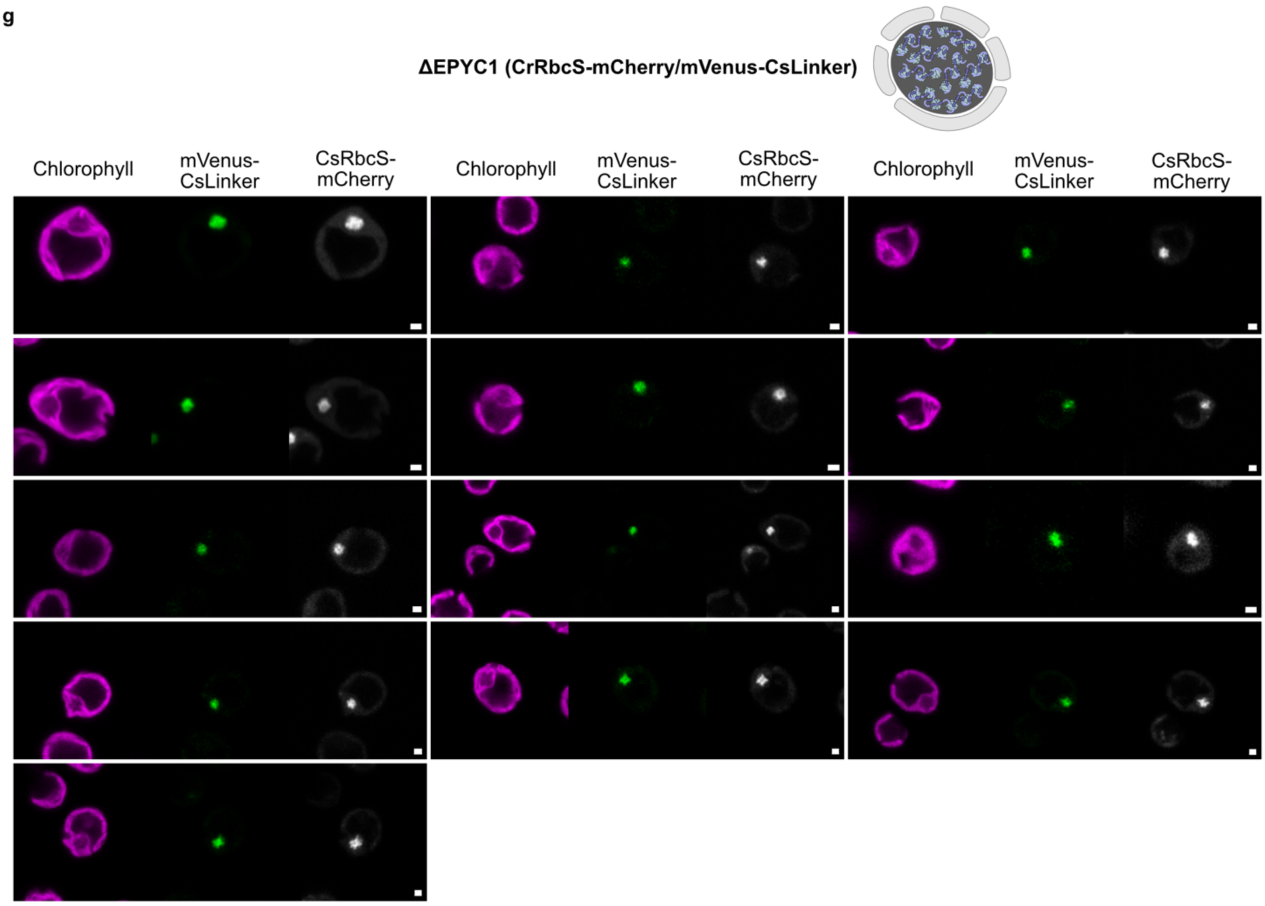


h

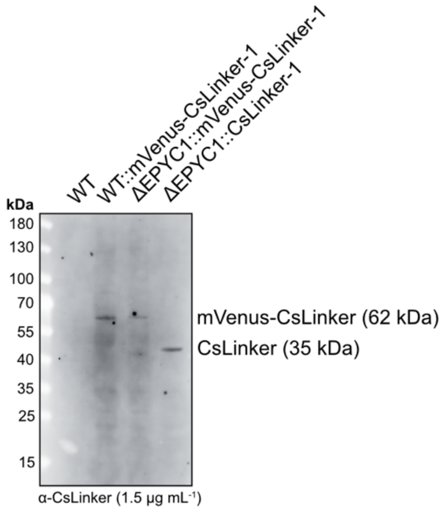


i

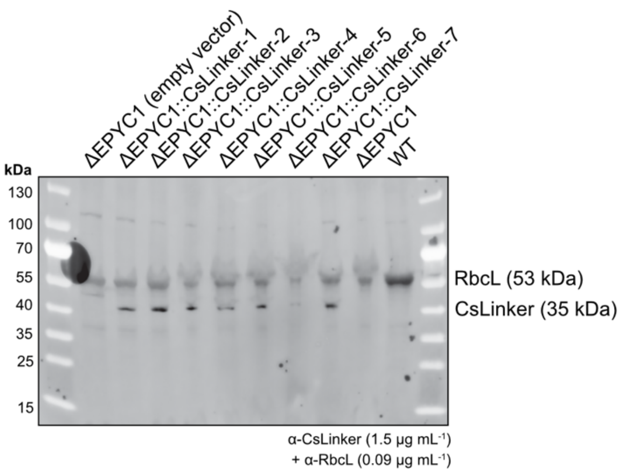


**Extended Data Fig. 19 | Cross-reactivity of α3-α4. a**, Example of Rubisco partitioning calculation using integrated density analysis in Fiji, according to ref.[15]. **b**, Rubisco partitioning in the condensate in the WT (CrRbcS-mCherry/EPYC1-Venus), ΔEPYC1 (CrRbcS-mCherry) and ΔEPYC1 (CrRbcS-mCherry/CsLinker) lines, quantified from the images in e, f and g using the method outlined in a. Statistical significance from unpaired t-tests are indicated; **** = $p < 0.0001$. **c**, Area of the Rubisco condensate as measured in Fiji, according to region 'A' in a. For the ΔEPYC1 (CrRbcS-mCherry), the largest condensed fluorescence signal at the canonical position was measured. *** = $p < 0.001$. **d**, Estimated volume distributions of the Rubisco condensates in the three lines, assuming sphericity of the condensate and calculating from the cross-sectional area in b. *** = $p < 0.001$. **e**, Confocal fluorescence microscopy images of tagged RbcS and EPYC1 in the WT background. **f**, Images of tagged RbcS in the ΔEPYC1 background strain. **g**, Images of tagged RbcS and CsLinker in the ΔEPYC1 background. Scale bars in e-g = 1 µm. **h**, Western blot confirmation of CsLinker variant expression in WT and ΔEPYC1 background lines. **i**, Western blot confirmation of CsLinker expression in ΔEPYC1 background relative to the empty vector and background strains. RbcL was used as a loading control.

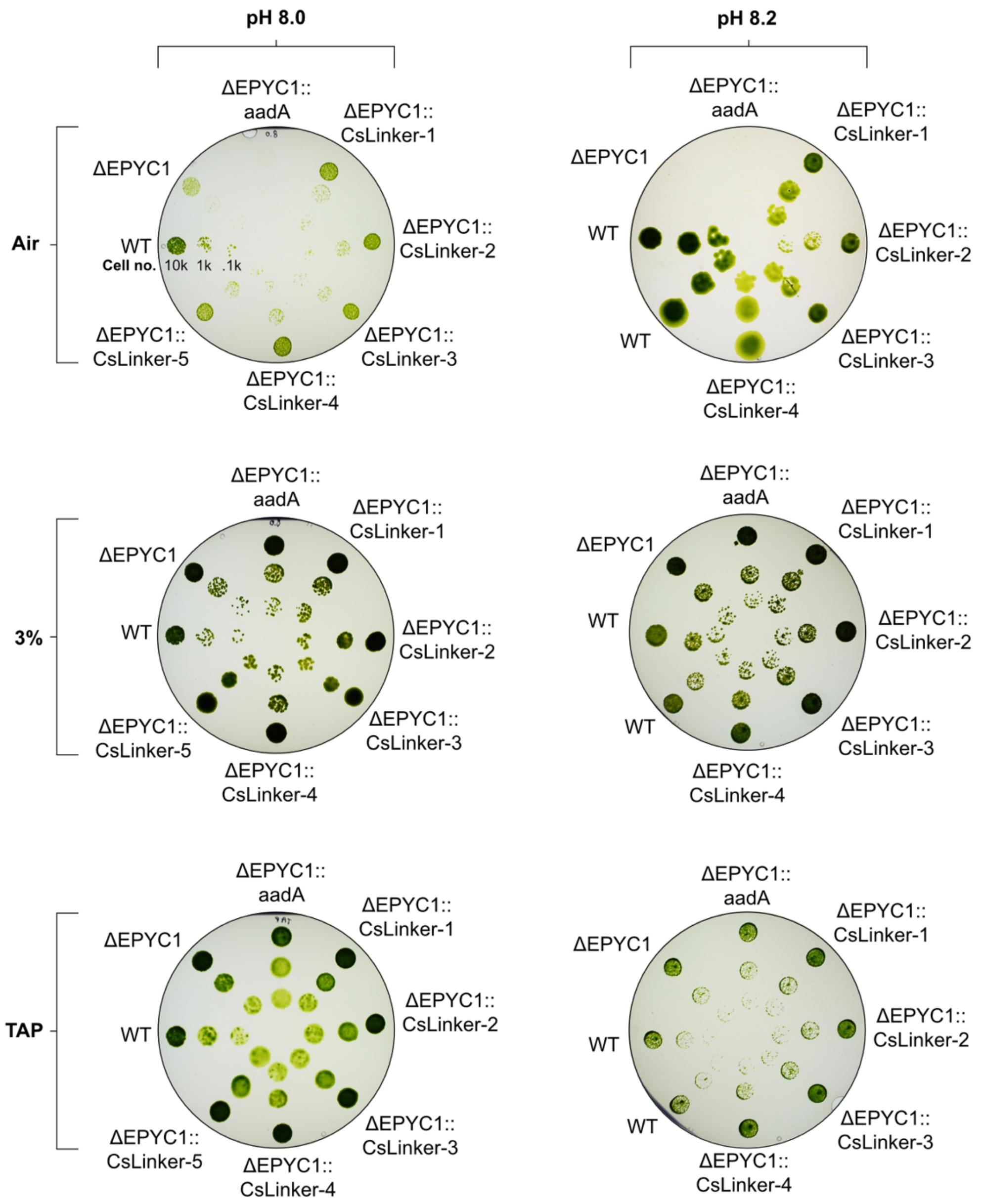


**Extended Data Fig. 20 | Spot test of CsLinker replacement lines.** Images of spot test plates following 5 days of growth at the indicated conditions. Images used in the Fig. 4 were taken from the pH 8.0 dataset.

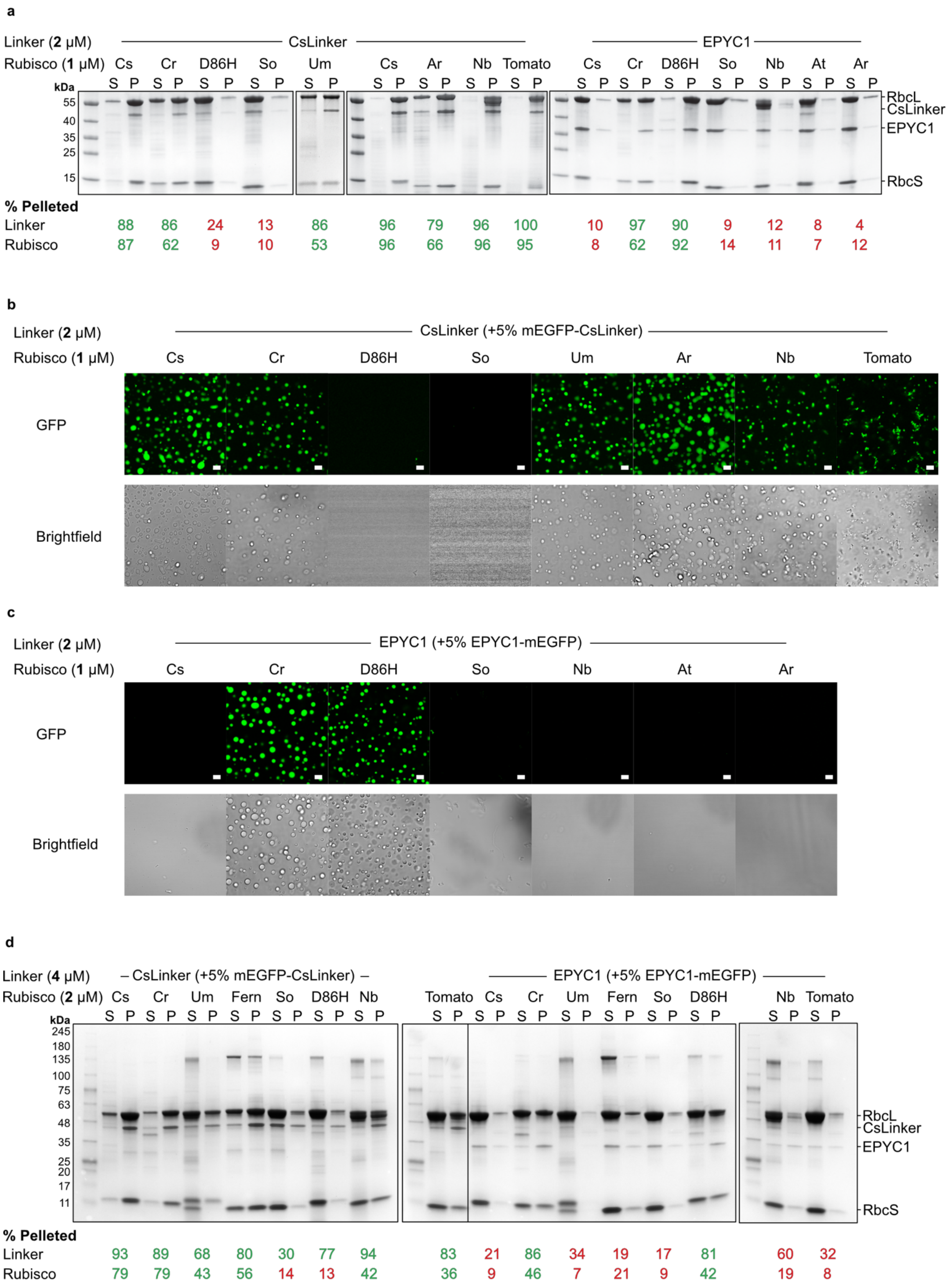

a
Linker (2 μM) CsLinker EPYC1
Rubisco (1 μM) Cs Cr D86H So Um Cs Ar Nb Tomato Cs Cr D86H So Nb At Ar
S P
kDa
55
40
35
25
15
RbcL
CsLinker
EPYC1
RbcS
% Pelleted
Linker 88 86 24 13 86 96 79 96 100 10 97 90 9 12 8 4
Rubisco 87 62 9 10 53 96 66 96 95 8 62 92 14 11 7 12
b
Linker (2 μM) CsLinker (+5% mEGFP-CsLinker)
Rubisco (1 μM) Cs Cr D86H So Um Ar Nb Tomato
GFP
Brightfield
c
Linker (2 μM) EPYC1 (+5% EPYC1-mEGFP)
Rubisco (1 μM) Cs Cr D86H So Nb At Ar
GFP
Brightfield
d
Linker (4 μM) – CsLinker (+5% mEGFP-CsLinker) – EPYC1 (+5% EPYC1-mEGFP)
Rubisco (2 μM) Cs Cr Um Fern So D86H Nb Tomato Cs Cr Um Fern So D86H Nb Tomato
S P
kDa
245
180
135
100
75
63
48
35
25
20
17
11
RbcL
CsLinker
EPYC1
RbcS
% Pelleted
Linker 93 89 68 80 30 77 94 83 21 86 34 19 17 81 60 32
Rubisco 79 79 43 56 14 13 42 36 9 46 7 21 9 42 19 8

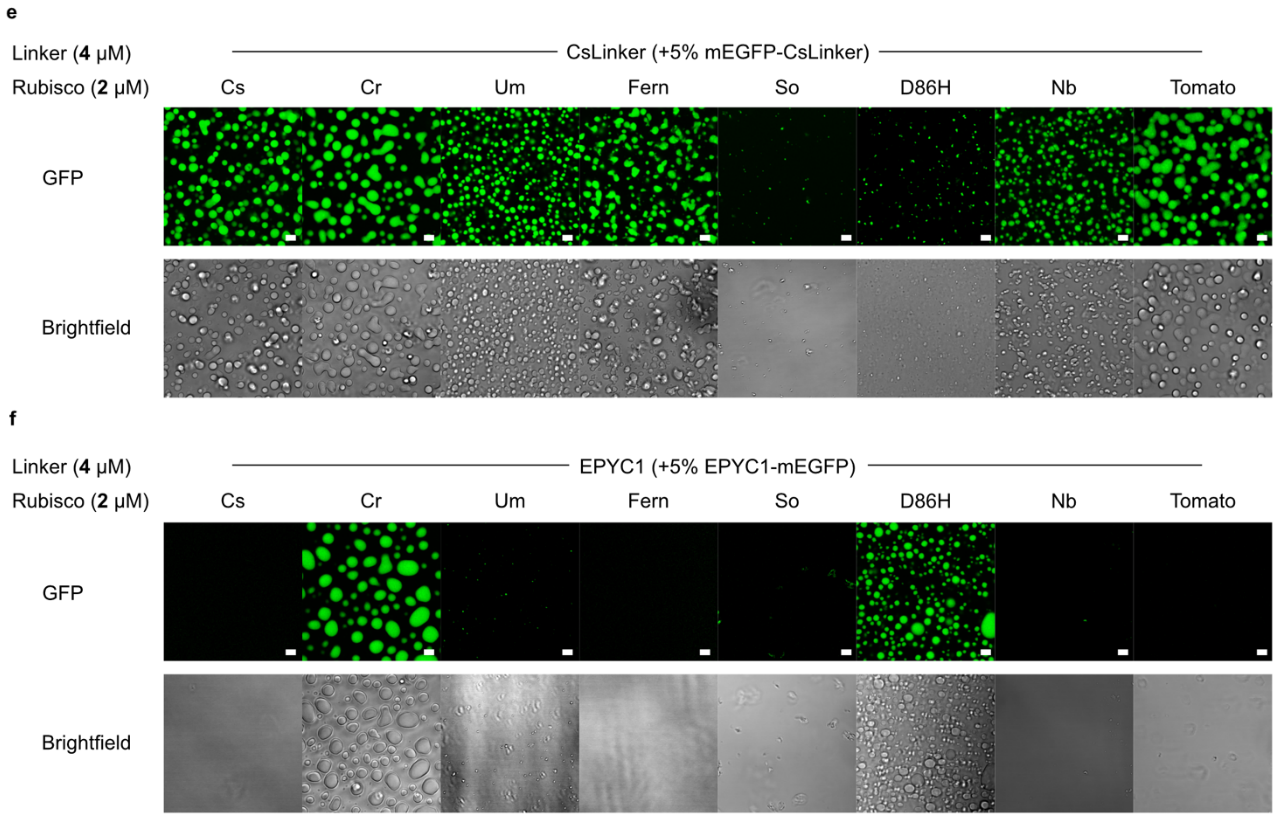


**Extended Data Fig. 21 | Cross-reactivity of CsLinker with green lineage Rubiscos. a**, Droplet sedimentation assays comparing the cross-reactivity of CsLinker and EPYC1 fixed at 2 μM, with Rubiscos from the green lineage fixed at 1 μM. The amount of Rubisco and Linker pelleted in each reaction is indicated below, with the numbers colored green if droplet formation was observed in the accompanying microscopy experiments in b. **b**, Confocal fluorescence microscopy and brightfield images of droplets formed with CsLinker under the same conditions as analyzed by droplet sedimentation assays in a, except mEGFP-CsLinker was included at a 5% molar ratio. **c**, Images of droplets formed with EPYC1 (+5% EPYC1-mEGFP molar ratio). Scale bar in b and c is 5 μm. **d**, Droplet sedimentation assays completed with CsLinker and EPYC1 fixed at 4 μM and green Rubiscos fixed at 2 μM. In these experiments, tagged linker was also included at 5% molar ratio as completed in the accompanying microscopy experiments. **e**, Images of droplets formed with CsLinker (+5% mEGFP-CsLinker molar ratio) at 4 μM. **f**, Droplet images with EPYC1 (+5% EPYC1-mEGFP) at 4 μM. Scale bar in e and f = 5 μm. Abbreviations: Cs = *Chlorella sorokiniana*, Cr = *Chlamydomonas reinhardtii*, Um = *Ulva mutabilis*, Ar = *Adiantum raddianum* (Fern), So = *Spinacia oleracea*, Nb = *Nicotiana benthamiana*, D86H = *Chlamydomonas reinhardtii* with D86H mutation made in RbcL.

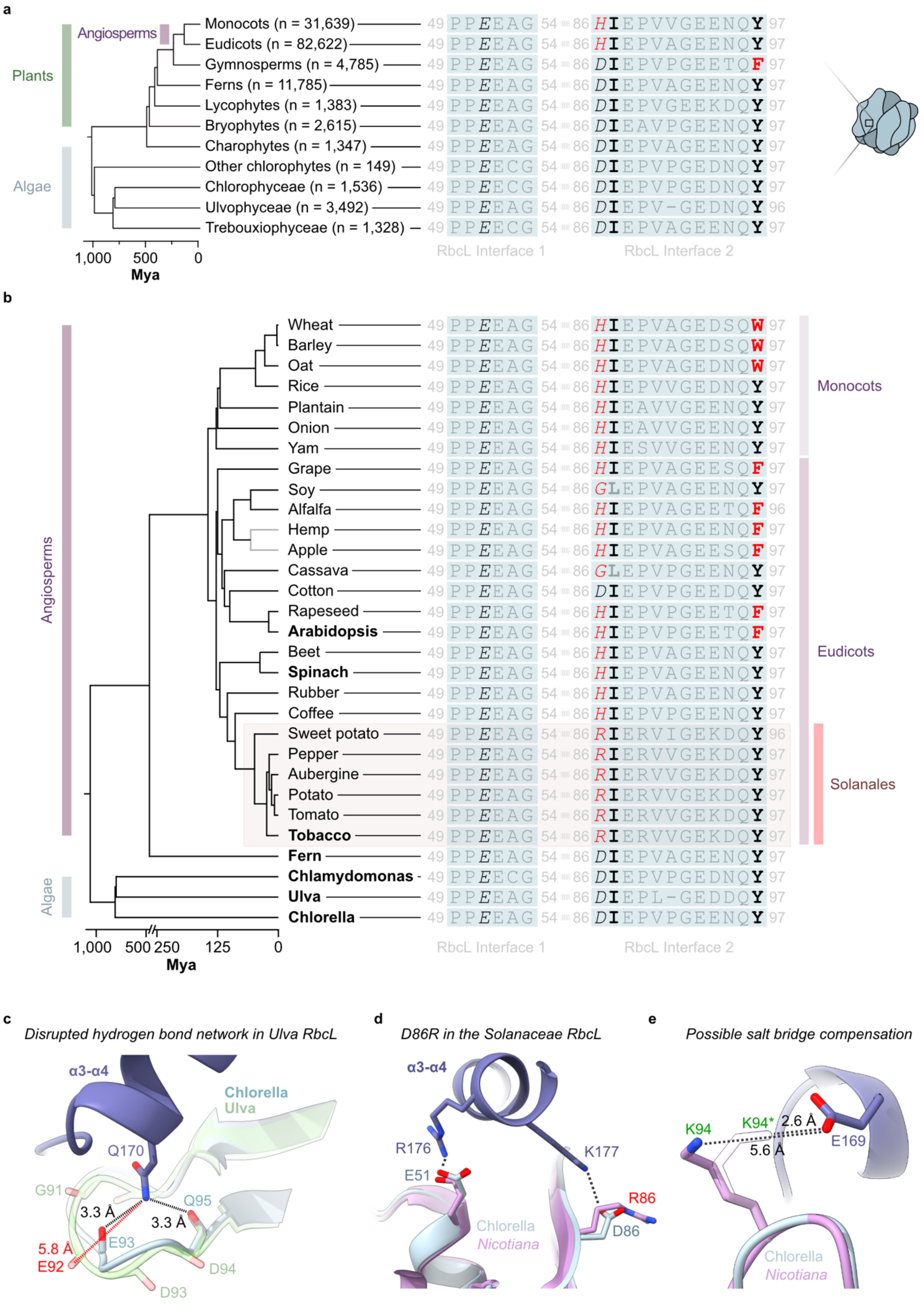
a
Plants
Angiosperms
Algae
Monocots (n = 31,639)
Eudicots (n = 82,622)
Gymnosperms (n = 4,785)
Ferns (n = 11,785)
Lycophytes (n = 1,383)
Bryophytes (n = 2,615)
Charophytes (n = 1,347)
Other chlorophytes (n = 149)
Chlorophyceae (n = 1,536)
Ulvophyceae (n = 3,492)
Trebouxiophyceae (n = 1,328)
1,000 500 0
Mya
RbcL Interface 1
RbcL Interface 2
b
Angiosperms
Algae
Wheat
Barley
Oat
Rice
Plantain
Onion
Yam
Grape
Soy
Alfalfa
Hemp
Apple
Cassava
Cotton
Rapeseed
Arabidopsis
Beet
Spinach
Rubber
Coffee
Sweet potato
Pepper
Aubergine
Potato
Tomato
Tobacco
Fern
Chlamydomonas
Ulva
Chlorella
Monocots
Eudicots
Solanales
1,000 500 250 125 0
Mya
RbcL Interface 1
RbcL Interface 2
c
Disrupted hydrogen bond network in Ulva RbcL
α3-α4
Chlorella
Ulva
Q170
G91
Q95
3.3 Å
3.3 Å
E93
5.8 Å
E92
D94
D93
d
D86R in the Solanaceae RbcL
α3-α4
R176
K177
E51
R86
Chlorella
Nicotiana
D86
e
Possible salt bridge compensation
K94
K94*
2.6 Å
5.6 Å
E169
Chlorella
Nicotiana

**Extended Data Fig. 22 | Variation of the CsLinker-interacting interface in plant RbcLs and potential alternative interactions at the interface.** **a**, Alignment of the consensus sequences of the CsLinker-binding interface in the RbcLs of major plant groups. Consensus sequences were produced from alignment of the indicated number of sequences in each class from NCBI. **b**, Alignment of the CsLinker-binding interfaces in the RbcLs of the 24 most valuable $C_3$ crop plants (FAOSTAT data) alongside the algal and fern Rubiscos tested in this study. **c**, The possibly disrupted hydrogen bond network in the βC-D loop of the *Ulva* RbcL. The AlphaFold 2 structural prediction of the *Ulva* RbcL (green) is shown aligned with the Chlorella RbcL coordinates solved in this study (blue). The disrupted hydrogen bond is annotated in red, with the corresponding lengths of the cognate and disrupted hydrogen bonds shown in red and black respectively. **d**, Demonstration of the disrupted salt bridge in the *Nicotiana* RbcL[40] due to the D86R substitution. **e**, A possible compensatory salt bridge in the K94 residue of the βC-D loop in the *Nicotiana* RbcL if an alternative residue conformer is occupied (K94*).

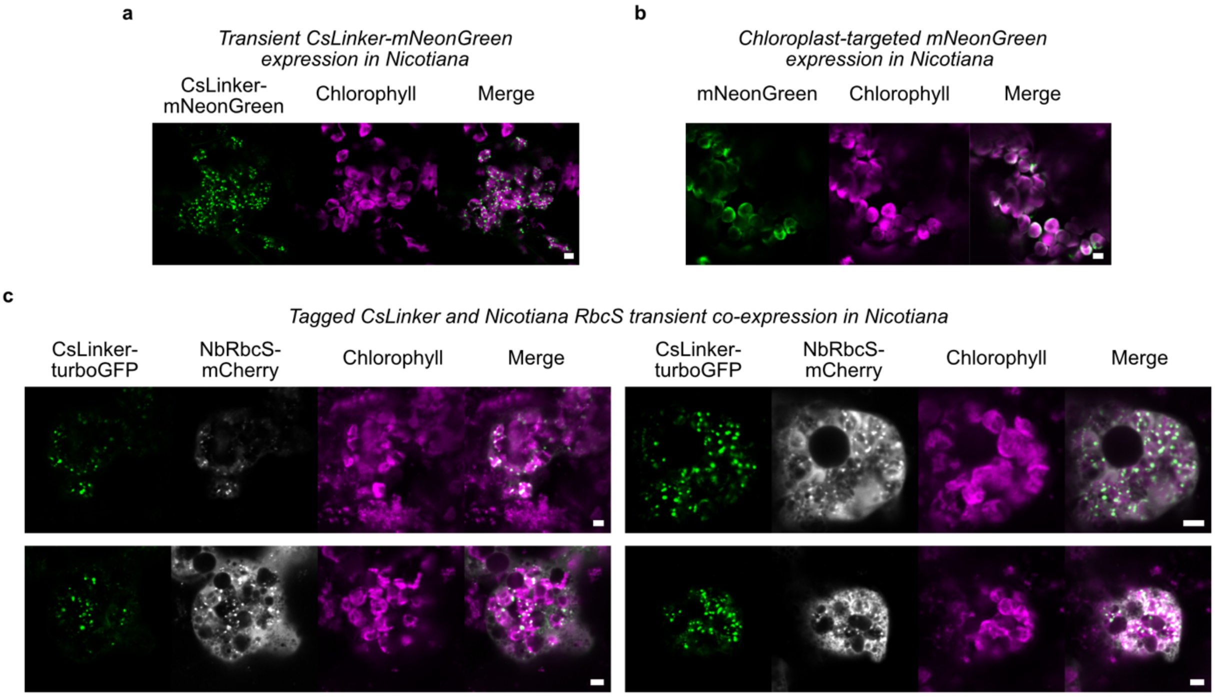


**Extended Data Fig. 23 | Transient expression of CsLinker in *Nicotiana benthamiana*. a**, Confocal microscopy image of turboGFP-tagged CsLinker transiently expressed alone in *Nicotiana*. **b**, Image of chloroplast-targeted (fused to *Arabidopsis* RbcS signal peptide; SP1A) turboGFP expressed alone, demonstrating a lack of puncta. **c**, Images of transient co-expression of CsLinker-turboGFP and mCherry-tagged *Nicotiana benthamiana* RbcS. Scale bars in a-c = 5 µm.

**Supplementary Table 3 | Quantification of immunogold-labelled RbcL in the pyrenoid.**

| Image number | Pyrenoid | Non-pyrenoid | % Pyrenoid | |
|---|---|---|---|---|
| 1 | 35 | 2 | 94.6 | |
| 2 | 48 | 0 | 100.0 | |
| 3 | 105 | 10 | 91.3 | |
| 4 | 27 | 0 | 100.0 | |
| 7 | 158 | 4 | 97.5 | |
| 8 | 61 | 4 | 93.8 | |
| | | | 96.2 | **Average** |
| | | | 3.5 | **S.D.** |

**Supplementary Table 4 | Standard curve values from absolute quantification experiments.**

| Purified Protein Standards | | | | | |
|---|---|---|---|---|---|
| **Sample ID** | **[Input] (pM)** | **MS1 Peak Area** | **Sample ID** | **[Input] (pM)** | **MS1 Peak Area** |
| CsLinker 1 | 126000 | 2.86E+05 | CsRbcL 1 | 27000 | 9.31E+04 |
| CsLinker 2 | 126000 | 1.14E+05 | CsRbcL 2 | 27000 | 2.07E+05 |
| CsLinker 1 | 1260000 | 2.77E+06 | CsRbcL 1 | 270000 | 7.80E+05 |
| CsLinker 2 | 1260000 | 3.81E+06 | CsRbcL 2 | 270000 | 1.93E+06 |
| CsLinker 1 | 12600000 | 4.24E+07 | CsRbcL 1 | 2700000 | 1.94E+07 |
| CsLinker 2 | 12600000 | 2.42E+07 | CsRbcL 2 | 2700000 | 1.10E+07 |

**Supplementary Table 5 | Measured values from whole cell lysate in absolute quantification.**

| Chlorella cell analysis (from 6E+6 cells) | | | | |
|---|---|---|---|---|
| **Sample ID** | **Analyte** | **MS1 Peak Area** | **Analyte** | **MS1 Peak Area** |
| Chlorella 1a | CsLinker | 1.11E+06 | CsRbcL | 9.57E+05 |
| Chlorella 1b | CsLinker | 1.34E+06 | CsRbcL | 1.26E+06 |
| Chlorella 2a | CsLinker | 9.45E+05 | CsRbcL | 9.60E+05 |
| Chlorella 2b | CsLinker | 1.19E+06 | CsRbcL | 1.23E+06 |
| Chlorella 3a | CsLinker | 8.92E+05 | CsRbcL | 1.03E+06 |
| Chlorella 3b | CsLinker | 1.06E+06 | CsRbcL | 1.10E+06 |

**Supplementary Table 6 | Calculation of CsRubisco chloroplast concentration from absolute quantification data.**

| | | | |
|---|---|---|---|
| CsRubisco MS1 intensity (rep 1) | 1110011.8 | fmol (/5613) | 197.8 |
| CsRubisco MS1 intensity (rep 2) | 1095989.3 | fmol (/5613) | 195.3 |
| CsRubisco MS1 intensity (rep 3) | 1067985 | fmol (/5613) | 190.3 |
| | | average fmol per sample | 194.4 |
| | | average mol per sample | 1.94E-13 |
| | | mol / cell (/6E+6) | 3.24E-20 |
| | | [chloroplast] (M) (/1.5E-14) | 2.16E-6 |
| | | **[chloroplast] (μM)** | **2.16** ± 0.04 S.D. |
| | | molecules / cell (av. mol*NA) | 1.95E+04 |

**Supplementary Table 7 | Calculation of CsLinker chloroplast concentration from absolute quantification data.**

| | | | |
|---|---|---|---|
| CsLinker MS1 intensity (rep 1) | 1224976 | fmol (/2641) | 463.8 |
| CsLinker MS1 intensity (rep 2) | 1064847 | fmol (/2641) | 403.2 |
| CsLinker MS1 intensity (rep 3) | 975129.7 | fmol (/2641) | 369.2 |
| | | average fmol per sample | 412.1 |
| | | average mol per sample | 4.12E-13 |
| | | mol / cell (/6E+6) | 6.87E-20 |
| | | [chloroplast] (M) (/1.5E-14) | 4.58E-6 |
| | | **[chloroplast] (μM)** | **4.58** ± 0.53 S.D. |

| | |
|---|---|
| molecules / cell (av. mol*NA) | 4.13E+04 |

**Supplementary Table 8 | Calculation of CsRubisco chloroplast concentration from western blot data.**

| | |
|---|---|
| Intensity of 100 ng band | 3349.175 |
| Intersect no. of cells (/6688) | 5.008E+06 |
| ng per cell (/5.008E+06) | 1.997E-05 |
| pg per cell (*1000) | 1.997E-02 |
| mg per cell (/1E-19) | 1.997E-11 |
| volume chloroplast (mL) | 1.500E-11 |
| [chloroplast] (mg mL$^{-1}$) | 1.33 |
| **[chloroplast] (µM) (/0.55)** | 2.42 |

**Supplementary Table 10 | Hill model fit parameters for α3-α4 native PAGE gel-shift assay data.**

| | Experiment | | | | |
|---|---|---|---|---|---|
| | Tris (1) | Tris (2) | Cryo | 'Mean' | All data |
| $K_{0.5}$ (μM) | 1.323 | 1.319 | 1.292 | 1.161 | 1.327 |
| $CI_{95}$ ($K_{0.5}$) | 1.138-1.528 | 1.009-1.656 | 1.163-1.438 | 0.9311-1.422 | 1.206-1.454 |
| Hill coefficient | 2.465 | 1.567 | 2.47 | 1.876 | 2.075 |
| $CI_{95}$ (hill coeff.) | 1.968-3.172 | 0.9824-2.449 | 1.971-3.161 | 1.374-2.568 | 1.747-2.428 |
| $R^2$ | 0.9809 | 0.8488 | 0.9791 | 0.925 | 0.9441 |

**Supplementary Table 11 | Fit parameters for SPR data.**

| | Experiment | | | | | |
|---|---|---|---|---|---|---|
| | α3-CsRubisco | α3-CrRubisco | α3-UmRubisco | α3-α4-CsRubisco | α3-α4-CrRubisco | α3-α4-UmRubisco |
| $K_D$ | 102.5 | 108.3 | 162.3 | 1.213 | 1.300 | 1.556 |
| $CI_{95}$ ($K_D$) | 79.88-132.5 | 102.3-114.8 | 157.6-167.2 | 1.107-1.329 | 1.157-1.460 | 1.404-1.723 |
| Fitted $B_{max}$ | 1860 | 1648 | 2378 | 1634 | 1683 | 1886 |
| $CI_{95}$ ($B_{max}$) | 1684-2075 | 1608-1689 | 2345-2413 | 1593-1677 | 1626-1742 | 1827-1946 |
| $R^2$ | 0.9797 | 0.9988 | 0.9998 | 0.9955 | 0.9922 | 0.9943 |

**Supplementary Table 12 | Collection, refinement, and validation of cryo-EM models.**

| **Data Collection and Processing** | | |
|---|---|---|
| Magnification | 240,000 | |
| Voltage (kV) | 200 | |
| Electron fluence (e-/Å$^2$) | 50 | |
| Pixel Size (Å$^2$) | 0.574 | |
| Symmetry | D4 | C1 |
| Figure for map | Ext. Data Fig. 14g | Ext. Data Fig. 15a |
| Initial Particles | 237,035 | 591,696 (8x expanded) |
| Final Particles | 73,962 | 133,171 |
| Map resolution (Å) | 2.39 | 2.77 |
| FSC Threshold | 0.143 | 0.143 |
| Map resolution range (Å) | 8.05-2.23 | 9.42-2.29 |
| **Refinement** | **CsRubisco** | **α3-α4-CsRubisco** |
| Map sharpening *B* factor (Å$^2$) | 50 | 45 |
| *Model Composition* | | |
| Non-H atoms | 35979 | 35836 |
| Residues | 4424 | 4440 |
| Water | 1267 | 1011 |
| *Mean B factors (Å$^2$)* | | |
| Protein | 44.00 | 44.10 |
| Water | 42.40 | 47.97 |
| *RMS deviations* | | |
| Bond length (Å) | 0.008 | 0.009 |
| Bond angles (°) | 1.164 | 1.171 |
| **Validation** | | |
| MolProbity Score | 1.30 | 1.30 |
| Clashscore | 2.09 | 2.20 |
| Poor rotamers % | 0 | 0 |
| *Ramachandran plot* | | |
| Favored % | 95.41 | 95.61 |
| Allowed % | 4.59 | 4.39 |
| Outliers | 0 | 0 |
| PDB Code | 8Q04 | 8Q05 |

**Supplementary Table 13 | PDBePISA analysis of α3-α4-CsRubisco complex interface.**

| **Salt bridges** | | | |
|---|---|---|---|
| α3-α4 residues (atom) | CsRbcL residue (atom) | Distance (Å) | Secondary structure |
| **Arg176 (NE)** | **Glu51 (OE2)** | **3.04** | **αB** |
| Arg176 (NE) | Glu51 (OE1) | 3.93 | αB |
| Arg176 (NH2) | Glu51 (OE2) | 2.96 | αB |
| **Lys177 (NZ)** | **Asp86 (OD1)** | **3.56** | **βC** |
| **Hydrogen bonds** | | | |
| Gly165 (O) | Gly92 (N) | 2.66 | CD loop |
| ***Gln170 (NE2)*** | ***Glu93 (O)*** | ***3.29*** | ***CD loop*** |
| ***Gln170 (NE2)*** | ***Gln96 (O)*** | ***3.30*** | ***CD loop*** |
| Arg176 (NE) | Glu51 (OE2) | 3.04 | αB |
| Arg176 (NH2) | Glu51 (OE2) | 2.96 | αB |
| Lys177 (NZ) | Ile87 (O) | 2.68 | βC |
| Lys177 (NZ | Asp86 (OD1) | 3.56 | βC |

| **α3-α4 Interface** | | | |
|---|---|---|---|
| α3-α4 residue | Accessible surface area (Å$^2$) | Buried surface area (Å$^2$) | % Buried |
| Gly164 | 121.72 | 28.64 | 23.5 |
| Gly165 | 84.64 | 31.20 | 36.9 |
| **Leu166** | **105.80** | **36.00** | **34.0** |
| Ser167 | 63.01 | 15.15 | 24.0 |
| Ala168 | 87.15 | 0.00 | 0.0 |
| Glu169 | 73.76 | 23.02 | 31.2 |
| ***Gln170*** | ***95.46*** | ***95.46*** | ***100.0*** |
| Arg171 | 46.76 | 0.00 | 0.0 |
| Glu172 | 48.95 | 0.00 | 0.0 |
| **Phe173** | **90.09** | **90.09** | **100.0** |
| **Leu174** | **82.49** | **12.39** | **15.0** |
| Glu175 | 63.88 | 0.00 | 0.00 |
| Arg176 | 158.68 | 62.48 | 39.4 |
| Lys177 | 149.89 | 64.24 | 42.9 |
| Ala178 | 89.52 | 0.00 | 0.0 |
| Arg179 | 124.39 | 0.00 | 0.0 |
